# *IL18* and pyroptosis-related genes show contrasting expression in human cardiac pericytes and endothelial cells

**DOI:** 10.64898/2026.09.06.749635

**Authors:** Daisuke Oryoji, Goro Doi, Sho Fujimoto, Naoya Nishimura, Kyoko Otsuka, Ayako Kuwahara, Masahiro Ayano, Yasutaka Kimoto, Koichi Akashi, Hiroaki Niiro, Hiroki Mitoma

## Abstract

Interleukin-18 (IL-18) contributes to cardiac inflammation, fibrosis and contractile dysfunction, and its maturation and release involve inflammatory caspases and gasdermin D (GSDMD). In two discovery cohorts (42 and 38 donors), donor-paired comparisons of public single-nucleus RNA-sequencing data from human hearts showed *IL18* relatively enriched in pericytes and *CASP1*, *CASP4* and *GSDMD* in endothelium. We specified these four directions as predictions and tested them against prespecified confirmation criteria in 22 adult non-failing donors from two additional resources. All four directions held in all 22 donors, each gene meeting the criteria. In prediction-based evaluations planned after the initial confirmation, endothelial enrichment of *IL18BP* and *CARD8* was also confirmed in all 22 donors. Among 52 cardiomyopathy donors, *IL18* favoured pericytes in 51, and *CASP1*, *CASP4* and *GSDMD* favoured endothelium in all 52. In exploratory analyses of three ischaemic-heart resources planned after confirmation, the four median directions were preserved in all five groups. In both acute myocardial infarction cohorts, pericyte-class *CASP4* relative RNA abundance was higher than in controls and the difference from endothelium was smaller. We propose the configuration hypothesis of intercellular division of labour: in the cardiac microvasculature, pericytes supply the IL-18 precursor and endothelium carries out its processing and release. This hypothesis reframes IL-18 maturation and release not only as a process completed within a single cell but also as a reaction accomplished through cooperation between cells with different roles.

## Introduction

Inflammasome activation and subsequent pyroptosis, an inflammatory form of cell death, have recently attracted attention as mechanisms that drive tissue injury and inflammation in cardiovascular diseases, including heart failure and myocardial infarction [26,31]. IL-18 is an important inflammatory cytokine that promotes cardiac inflammation and contributes to fibrosis and contractile dysfunction. Its inhibition improved recovery of contractile function in an ex vivo ischaemia–reperfusion model of human atrial tissue [7] and reduced cardiac inflammation and fibrosis in a mouse model of β-adrenergic stimulation [8]. This cytokine is produced intracellularly as the precursor pro-IL-18, which is cleaved and matured by caspase-1 activated within the inflammasome. Human caspase-4 and caspase-5 can also process this precursor directly [1,2]. These inflammatory caspases also cleave gasdermin D (GSDMD). Membrane pores formed by its N-terminal fragment release mature IL-18 and initiate the membrane injury of pyroptosis [3,4]. Studies using macrophages and keratinocytes have established that the sequence linking stimulus sensing, inflammatory caspase activation, and cytokine maturation and release can proceed within the same cell [5,6]. In the human heart, however, it remains necessary to establish which cells carry out these steps and how closely sensing, caspase activation, GSDMD-mediated membrane injury, and IL-18 maturation and release are coupled within individual cells.

This is because the cells initiating an inflammatory response in the heart have been suggested to differ from those carrying out downstream reactions. In experiments using mouse cardiac cells, inflammasome activation initiated in cardiomyocytes led to caspase-1 activation and inflammatory cell death in fibroblasts connected by membrane nanotubes. This response required NLRP3 in cardiomyocytes but did not require NLRP3 in the fibroblasts in which caspase-1 was activated [9]. Outside the heart, experiments using human monocytic cells showed the release of microparticles containing active caspase-1 and cleaved GSDMD that induced cell death in pulmonary microvascular endothelial cells [28]. Extracellular vesicles released by pyroptotic cells have also been shown to transfer GSDMD pores to the membranes of other cells and induce lytic cell death [29]. These findings show that understanding inflammatory caspase activation and GSDMD-mediated membrane injury requires attention both to intracellular reactions and to the relationships between cells supplying the relevant molecules and cells receiving their effects.

The human heart contains vascular and immune cells in addition to cardiomyocytes and fibroblasts. Microvascular endothelial cells and pericytes occupy adjacent compartments while maintaining distinct cellular properties and transcriptional profiles [10]. In failing ischaemic hearts, immunostaining identified IL-18 protein mainly in endothelium and macrophages [18]. Protein localisation, however, does not identify the cells that produced it. Nucleus-level atlases can address which cell classes contain transcripts for the precursor and its processing machinery. To investigate how IL-18-related inflammatory responses could be coordinated in such tissue, the starting point is to identify the cell classes expressing transcripts for the precursor, the enzymes that process it and the pore-forming protein. Whether these biochemically related transcripts favour the same or different cell classes determines the combinations of cells and molecules to examine next in studies of protein localisation and cell interactions. Single-cell and single-nucleus atlases provide a basis for describing these distributions [10]. Comparing cell classes within donors and testing whether the pattern recurs in independent donors can establish the reproducibility of the transcript pattern.

We examined *IL18* and related transcripts across major cardiac cell classes using donor-paired comparisons in two public single-nucleus datasets. This exploration identified a configuration in which *IL18* was relatively enriched in pericytes and *CASP1*, *CASP4* and *GSDMD* in endothelial cells. We then specified these four directions as predictions and tested them against prespecified confirmation criteria in donors from additional resources who were separate from the discovery donors. Prespecified secondary analyses examined the scope of support. In exploratory analyses planned after confirmation, we also examined these transcript distributions in acute myocardial infarction and ischaemic cardiomyopathy and compared cell-class transcript abundance and the endothelial–pericyte difference with controls in acute myocardial infarction. Through these analyses, we revealed the cell-class transcript distributions of the IL-18 precursor and the molecules involved in its processing and release and examined how this configuration relates to ischaemic myocardial injury.

## Methods

### Study design, data sources and units of analysis

This study reanalysed publicly available, de-identified human-heart data. The discovery survey used only the single-nucleus components of SCP1303 and GSE183852 [11,12]. SCP1303 included 42 donors: 16 non-failing, 11 with dilated cardiomyopathy and 15 with hypertrophic cardiomyopathy. The GSE183852 nuclei dataset included 38 donors: 25 classified as Donor and 13 with dilated cardiomyopathy. Figure 1A shows the study design, and Table 1 lists the resources, donors and nuclei used in the analyses. For *GSDMD*, endothelial cells were also compared with cardiomyocytes using these two single-nucleus datasets and the GeoMx spatial dataset GSE271676 [13].

**Figure 1.**
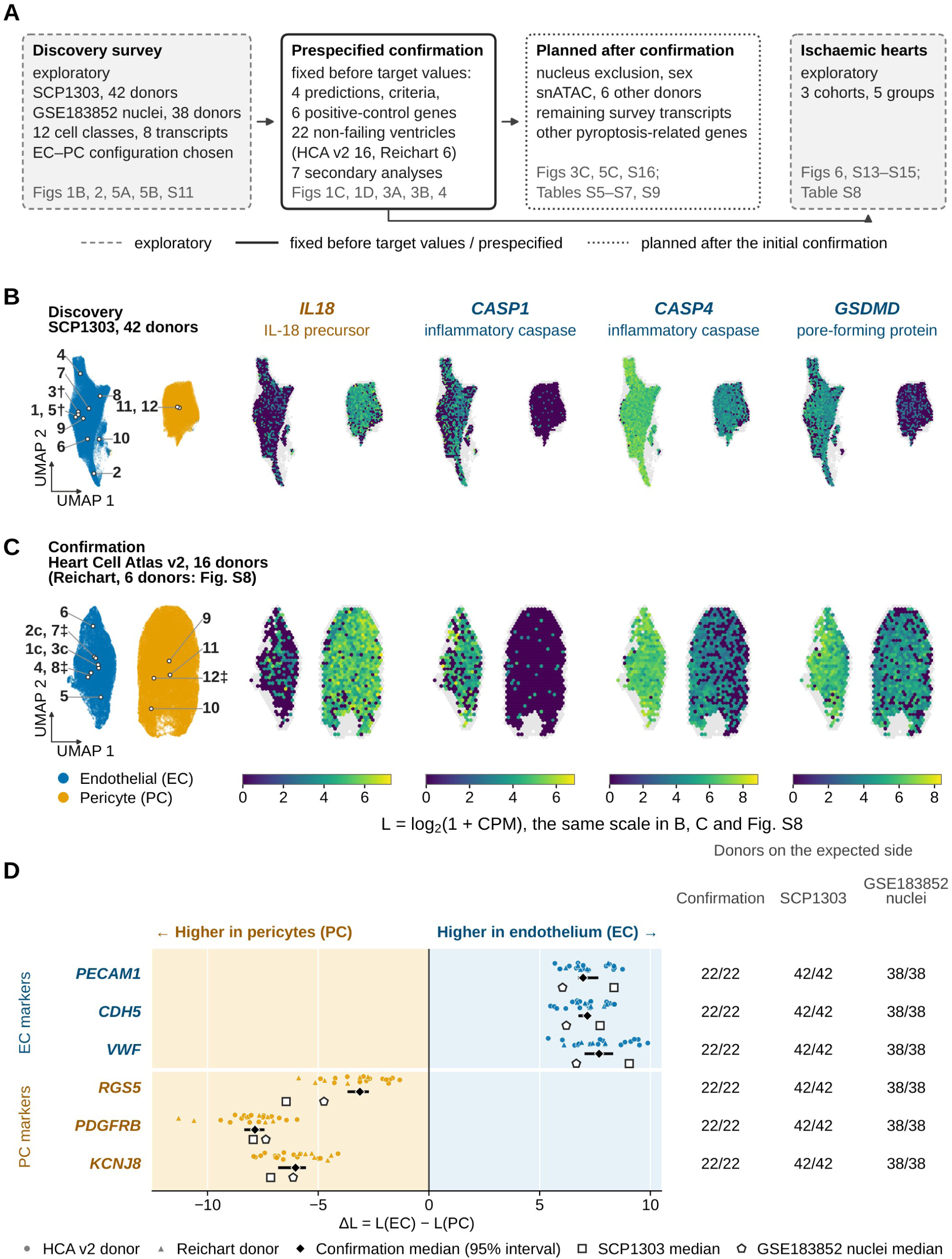
Study design, endothelial and pericyte nuclei in shared embeddings with transcript bin maps, and cell-class positive controls. (A) Study design. Solid boxes indicate stages fixed before the target values were read and prespecified confirmation, dashed boxes indicate exploratory analyses, and dotted boxes indicate checks planned after the initial confirmation. Corresponding figures and tables are shown below each box. Arrows indicate the order of the analyses. (B) Discovery cohort SCP1303: 42 donors, 101,307 endothelial and 69,304 pericyte nuclei. (C) Confirmation cohort Heart Cell Atlas v2: 16 of the 22 confirmation donors, 19,740 endothelial and 42,718 pericyte nuclei. The six Reichart donors are shown in Supplementary Figure S8. The left panels in B and C show UMAPs of the nuclei used for the donor-level comparisons. Numbered points mark the median position of each source cell state. The symbol † denotes endocardial and lymphatic states included in the discovery endothelial class, c denotes states used in the capillary secondary analysis, and ‡ denotes states excluded in the secondary analysis removing ‘-like’ states. The right panels show hexagonal bins of the same embeddings coloured by L for *IL18*, *CASP1*, *CASP4* and *GSDMD* (Methods). Grey bins contain fewer than 30 nuclei. Each transcript has one colour scale, shared by B, C and Supplementary Figure S8. (D) ΔL = L(EC) − L(PC) for the six cell-class positive-control genes fixed before the target values were read. Points represent the 22 confirmation donors, with circles for Heart Cell Atlas v2 and triangles for Reichart. Diamonds and horizontal bars show pooled medians and 95% bootstrap intervals from 10,000 donor resamples stratified by dataset. Open squares and pentagons show the discovery medians for SCP1303 and GSE183852 nuclei, in that order. The numbers at right give the number of donors on the side predicted by the class annotation and the total number of donors, in the order confirmation, SCP1303 and GSE183852 nuclei. Values are provided in Supplementary Data 1. State numbers in B are: 1 EC-ADAMTS9, 2 EC-DKK2, 3 EC-Endocardial†, 4 EC-LHX6, 5 EC-Lymphatic†, 6 EC-NEBL, 7 EC-PKD1L1, 8 EC-TMEM163, 9 EC-X1, 10 EC-X2, 11 PC- S1, 12 PC-S2. State numbers in C are: 1 EC1_cap (c), 2 EC2_cap (c), 3 EC3_cap (c), 4 EC4_immune, 5 EC5_art, 6 EC6_ven, 7 EC9_FB-like (‡), 8 EC10_CMC-like (‡), 9 PC1_vent, 10 PC2_atria, 11 PC3_str, 12 PC4_CMC-like (‡). EC, endothelium; PC, pericyte.

**Table 1.**
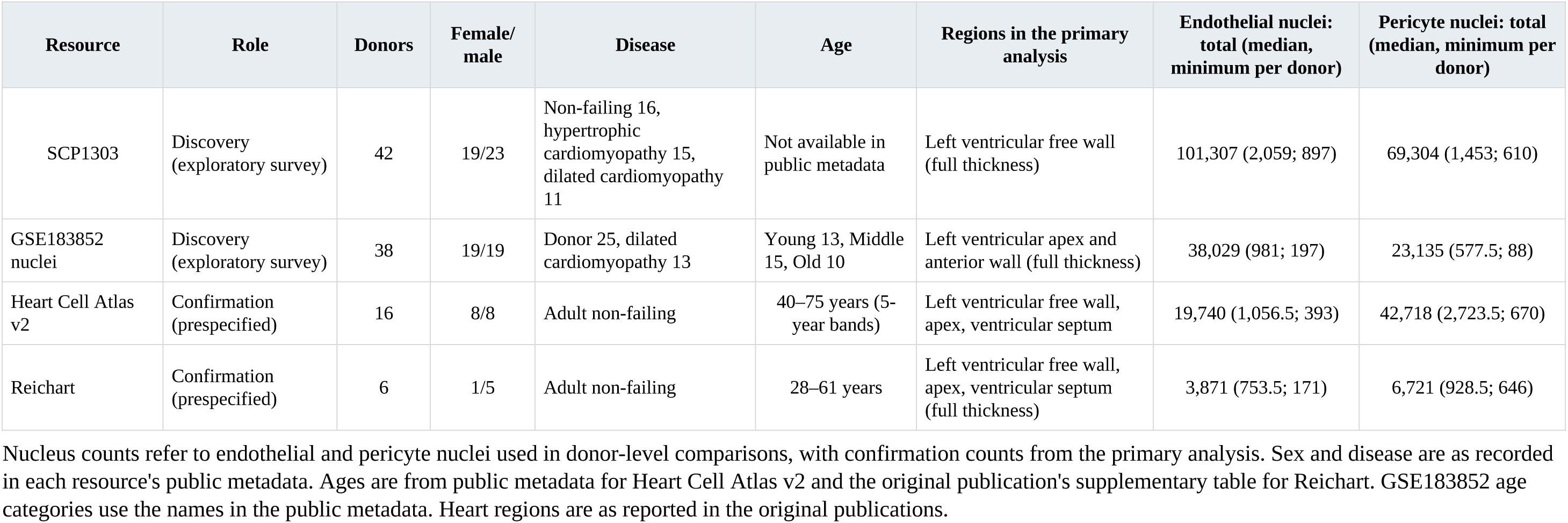
Resources, donors and nuclei used in the analyses.

Donors, rather than individual nuclei, were the units of inference [14,15]. Source-provided class annotations were used as supplied. Vascular endothelium was defined by the source primary class, cardiac endothelial cell, with lymphatic endothelium and endocardium treated as separate classes according to the source annotations. Pericytes and vascular smooth-muscle cells were treated separately. Twelve corresponding class definitions were used for cross-cohort comparisons.

### Relative RNA abundance and paired differences

For gene g, donor d and class c, raw counts were summed over the nuclei assigned to that donor–class group. The library denominator was the sum over all available features in that group. Relative abundance was defined as L = log2(1 + 10⁶ × gene count / library count). A donor–class group was eligible when it contained at least 30 nuclei and had a positive library total. Groups with 0 target counts were assigned L = 0.

The principal difference was ΔL = L(endothelium) − L(pericyte), calculated only when both classes were eligible within the same donor. Negative values favour pericytes. This quantity compares proportions of measured class libraries after transformation.

The eight-transcript survey comprised *IL18*, *CASP1*, *CASP4*, *CASP5*, *GSDMD*, *IL18R1*, *IL18RAP* and *IL18BP*. Sixty-six pairs among the 12 matched classes produced 528 cross-cohort gene–pair records. The endothelial–pericyte set of four transcripts was selected from the survey results. Previously inspected marker information from candidate confirmation atlases and other organs was documented in Supplementary Information.

### Shared embedding of endothelial and pericyte nuclei

For display, endothelial and pericyte nuclei used in the donor-level comparisons were placed in one shared two-dimensional embedding per cohort. The embedding was divided into hexagonal bins, and each bin was summarised using the same transformation L as the donor–class aggregates (Supplemental Methods). The embedding was used for display, and inference was based on donor-level comparisons.

### Summaries, precision and discovery support criteria

We summarised each paired contrast by its median, a percentile donor-bootstrap 95% confidence interval, and the numbers of positive, negative and tied donors. Ties were defined by |ΔL| ≤ 10^-12^ and excluded from the directional fraction. The principal summaries used 10,000 bootstrap samples.

The discovery precision criteria required, in each cohort, at least 16 paired donors, at least eight donors with detection in either class, at least 12 non-tied differences and a confidence-interval width no greater than 1.0. A supported cross-cohort contrast additionally required matching median directions, an absolute median of at least 0.25 in each cohort, intervals excluding zero, a directional fraction of at least 0.75, and a ratio of absolute cohort medians no greater than four. Records failing precision were classified as undeterminable, whereas records meeting precision but not the remaining conditions were classified as not supported.

### Transcriptomic calibration

We reconstructed endothelial–pericyte contrasts for the 21,932 genes with non-duplicated names shared by both nucleus datasets. We compared the four selected transcripts with expression-matched reference genes and counted how often all four directions were simultaneously supported among the reference genes (Supplemental Methods).

### Sensitivity analyses

All additional analyses were specified and documented after the discovery data had been seen but before calculation. We evaluated disease strata, every leave-one-donor-out sample, minimum class sizes of 50 and 100 nuclei, and alternative denominators excluding mitochondrial genes, the ten highest-count non-target genes in each cohort, or both.

Library-depth sensitivity was examined by binomial thinning of aggregated counts. Five macrophage-associated reference genes (*C1QB*, *SPI1*, *TYROBP*, *FCER1G* and *C1QA*) were selected from 16 predefined candidates before association testing. We tested donor-level Spearman associations between pericyte *IL18* L and the L of each of the 16 candidates or the mean L of the five markers (Supplemental Methods). Endothelial–smooth-muscle and pericyte– smooth-muscle contrasts were also evaluated with the same four target genes.

### Prespecified independent confirmation

The confirmation analysis used adult non-failing ventricular donors from Heart Cell Atlas v2 and the Reichart resource [16,17]. Donor and library overlap was identified from documented tissue sources and identifiers and excluded. The four directions identified in discovery were used as predictions and evaluated under prespecified eligibility and decision rules. The cell-class positive controls were fixed before the target values were read: *PECAM1*, *CDH5* and *VWF* for the endothelial side and *RGS5*, *PDGFRB* and *KCNJ8* for the pericyte side. As a condition for reading the target values, each dataset required at least 90% of non-tied donors on the predicted side and a median in the predicted direction with an absolute value of at least 1.0 (Figure 1D).

For the Reichart resource, the input specification was amended before target evaluation to use 93 public filtered integer matrices. Eligibility assessment, cell-state assignment, donor ordering and calculation of per-nucleus library totals were completed before target evaluation.

The confirmation set comprised 16 Heart Cell Atlas donors and six Reichart donors. The initial confirmation evaluated four genes: *IL18*, *CASP1*, *CASP4* and *GSDMD*.

For each gene, confirmation required at least 16 donors overall and 12 non-tied contrasts, at least 75% of non-tied donors in the predicted direction, a pooled median whose 95% bootstrap interval lay wholly on the predicted side of zero, and an absolute median difference of at least 0.25. A contributing dataset with at least five donors contradicted confirmation when both its median and interval were in the reverse direction. Confirmation of the complete configuration additionally required all four genes to have no oppositely directed dataset median.

Intervals were calculated using 10,000 donor resamples stratified by dataset. Pooled estimates were the medians of the donor contrasts combined across the two datasets.

### Prespecified secondary confirmation analyses

Seven secondary analyses were performed. They comprised capillary endothelial–pericyte contrasts, exclusion of predefined ‘-like’ states, right ventricular samples, atrial samples, adult cardiomyopathy ventricular samples from Reichart, differences in the fraction of nuclei with count ≥1, and dataset-specific primary summaries. The Heart Cell Atlas/Reichart donor counts (Supplementary Table S3) were, in that order, 16/6, 16/6, 16/3, 15/0, 0/52, 16/6 and 16/6.

### Evaluation of the remaining survey transcripts

Of the eight survey transcripts, *CASP5*, *IL18R1*, *IL18RAP* and *IL18BP* had not been evaluated in the initial confirmation, and their evaluation in the confirmation set was planned after the initial confirmation results had been seen. A directional prediction was made only when the discovery endothelial–pericyte record was classified as supported under the discovery support criteria. Only *IL18BP* had a prediction under this rule, towards endothelium, whereas the other three transcripts were summarised descriptively. We used the same confirmation set, donor–class groups, confirmation criteria, saved resampling indices and seven secondary analyses as in the initial confirmation. Before this evaluation, values for these transcripts had been calculated in ischaemic-heart analyses that excluded the confirmation resources (Supplemental Methods).

### Evaluation of other pyroptosis-related genes

After the initial confirmation and evaluation of the remaining survey transcripts, we planned to evaluate ten other pyroptosis-related genes in the confirmation set. The scope covered molecules acting within a pyroptotic cell from sensing to plasma-membrane rupture, selected from sensors, adaptors, inflammatory caspases, substrates, and molecules mediating pore formation and membrane rupture. Proteases produced by other cells, regulators of activity or transcription, membrane repair, released molecules and molecules in apoptotic pathways were outside this scope. The selection was non-exhaustive, and the selected and excluded genes are listed by category in Supplemental Methods. Directional predictions were made only for genes whose endothelial–pericyte records in both cohorts were classified as supported in the discovery genome-wide calibration table. Under this rule, only *CARD8*, an inflammasome sensor that triggers pyroptosis in human cells [38,39], had a prediction, towards endothelium, whereas the other nine genes were summarised descriptively. We used the same confirmation set, donor–class groups, confirmation criteria, saved resampling indices and seven secondary analyses as in the initial confirmation. The predictions, criteria and interpretation rules were specified before values for the ten genes were read in this evaluation. Values in the confirmation set for the nine genes other than *IL1B* had already been calculated for a genome-wide comparison before this evaluation (Supplemental Methods).

### Comparison of deposited experimental structures

A quantitative comparison of deposited experimental structures was planned after the transcript analyses to relate transcript enrichment in the confirmation set to the molecular stages involving IL-18. Recognition residues on mature IL-18 (UniProt Q14116, residues 37–193) were defined in the IL-18–IL-18BP complex (PDB 7AL7) [41] and the complex of IL-18 with IL-18Rα (the product of *IL18R1*) and IL-18Rβ (the product of *IL18RAP*) (PDB 3WO4) [42]. A residue was included if any of its heavy atoms lay less than 4.5 Å from a protein heavy atom in the binding partner. The two complexes were superposed using the Cα atoms of mature IL-18, and close atom pairs were counted with each binding partner in its respective binding pose. Receptor assembly was assessed using all copies of IL-18 alone (PDB 3WO2) and in complex with IL-18Rα (PDB 3WO3) [42]. Comparison of pro-IL-18 and mature IL-18 used all NMR models in PDB 8URV and 1J0S [43,44]. The arrangement of recognition residues was assessed by comparing Cα distances between residue pairs at least four positions apart in the sequence with the corresponding distances in the bound structures (Supplemental Methods).

### Ischaemic-heart data and analyses

GSE270788 [19] included five donors with acute myocardial infarction (AMI), five with ischaemic cardiomyopathy (ICM) and seven controls. From Kuppe [20], we used seven AMI, eight ICM and four control donors, excluding one donor whose clinical group was coronary artery disease and heart failure. SCP1849 [21] included seven ICM and eight non-failing control donors. In Kuppe, counts from all specimens from each donor were summed, except in region-specific analyses. Cell classes followed the published annotations. Details of data import and class correspondence are provided in Supplemental Methods.

Definitions of L, ΔL, ties and donor–class eligibility were as above. Each group was summarised by the median ΔL, the number of donors in the predicted direction and the number of ties. The prespecified criterion was that at least 75% of non-tied donors had the predicted direction. For all quantities, percentile 95% bootstrap intervals were obtained by resampling donors with replacement 10,000 times within groups, or units within strata for region-specific analyses. Count-pooled L was calculated after separately summing the counts and library totals of eligible donors for each group and class. Count-pooled ΔL was calculated using donors eligible for both endothelium and pericytes, and its difference between AMI and controls was calculated. The change in each class was the AMI-minus-control difference in count-pooled L among donors eligible for that class. Changes in *IL18*, *CASP1* and *GSDMD* for each class were calculated in the same way. For each class-level change, two-sided P values were obtained by enumerating all donor-label assignments, and Benjamini–Hochberg q values used all classes within each cohort and gene as the family, comprising 14 classes in GSE270788 and 11 in Kuppe. Changes in *CASP4* were compared with changes in genes of similar abundance (Supplemental Methods).

In Kuppe, the unit was the patient–region combination, with specimens within the same unit pooled. Units were divided into five strata according to the recorded specimen region: control, AMI remote myocardium, border zone, infarct zone and ICM. Units eligible for both endothelium and pericytes were used to calculate count-pooled L within each stratum and its difference from controls. The contrast ratio κ was defined as the stratum’s count-pooled ΔL divided by the control count-pooled ΔL. At both stratum and unit levels, κ was compared with that of genes biased towards endothelium in controls and of similar abundance to *CASP4* (Supplemental Methods). In Kuppe, single-nucleus ATAC data from the same units were also used to examine chromatin accessibility around transcription start sites in endothelium and pericytes (Supplemental Methods, Supplementary Table S8).

### Checks for contamination in pericyte *CASP4*

We examined contamination from other cells as an explanation for higher pericyte *CASP4* in AMI using three approaches. These were (i) a contamination model based on five myeloid markers, (ii) subtraction of contributions from other classes using genes specific to myeloid cells, endothelium, fibroblasts and cardiomyocytes, and (iii) exclusion of pericyte nuclei bearing myeloid or endothelial markers. The pericyte-class change before nucleus exclusion was also estimated after leaving out one AMI or control donor at a time. Interpretation rules were specified before calculation (Supplemental Methods).

### Reproducibility and ethics

Independent implementations reconstructed the eight central cohort–gene summaries from integer pseudobulk arrays and compared them with the original results. Analysis records included input checksums, code, random-number initialisation values and output locations. Tables for the ischaemic-heart analyses were recalculated from source data using separately written import and calculation code, and agreement with the original numerical results was verified. The corresponding author had full access to all the data in the study and takes responsibility for the integrity of the data and the data analysis.

This secondary analysis used publicly available, de-identified data and involved no participant recruitment, intervention or tissue collection. Ethics and consent arrangements are reported by the source studies [11–13,16,17,19–21,32,33]. The source GSE183852 study reports Washington University Institutional Review Board approval 201104172.

### Use of generative artificial intelligence

ChatGPT and Codex (OpenAI) and Claude (Anthropic) were used to assist with manuscript preparation, development and checking of analysis and plotting code, and review of results. AI-assisted code and numerical outputs were checked against the source data using the verification procedures described above.

## Results

### *IL18* and three caspase–gasdermin transcripts show opposite endothelial–pericyte directions

Endothelial and pericyte nuclei occupied separate regions of a shared embedding in each of the three resources for which embeddings were generated (Figure 1B, 1C, Supplementary Figure S8). When the embedding was divided into bins and relative abundance was assessed within each bin, *IL18* was higher in the pericyte region, whereas *CASP1*, *CASP4* and *GSDMD* were higher in the endothelial region. This configuration was the same in the discovery cohort SCP1303 and the two confirmation resources. Coarser and finer bin grids showed the same configuration (Supplementary Figures S9, S10). The annotated classes also agreed with the six positive-control genes fixed before the target values were read. The endothelial controls *PECAM1*, *CDH5* and *VWF* and the pericyte controls *RGS5*, *PDGFRB* and *KCNJ8* were on the side predicted by the class annotation in all 22 confirmation donors and all 42 and 38 discovery donors (Figure 1D). The donor-level comparisons below quantify this configuration.

The exploratory class survey identified opposing endothelial–pericyte directions for *IL18* and *CASP1*, *CASP4* and *GSDMD*. This selected configuration met the discovery support criteria in both SCP1303 and GSE183852 nuclei (Figure 2A, 2B, Supplementary Table S1). *IL18* had median ΔL values of −2.121 (95% CI, −2.438 to −1.763) in SCP1303 and −1.317 (−1.751 to −0.850) in GSE183852 nuclei. The negative direction, favouring pericytes, occurred in 39/41 and 33/34 non-tied donors, with one and four ties, respectively.

**Figure 2.**
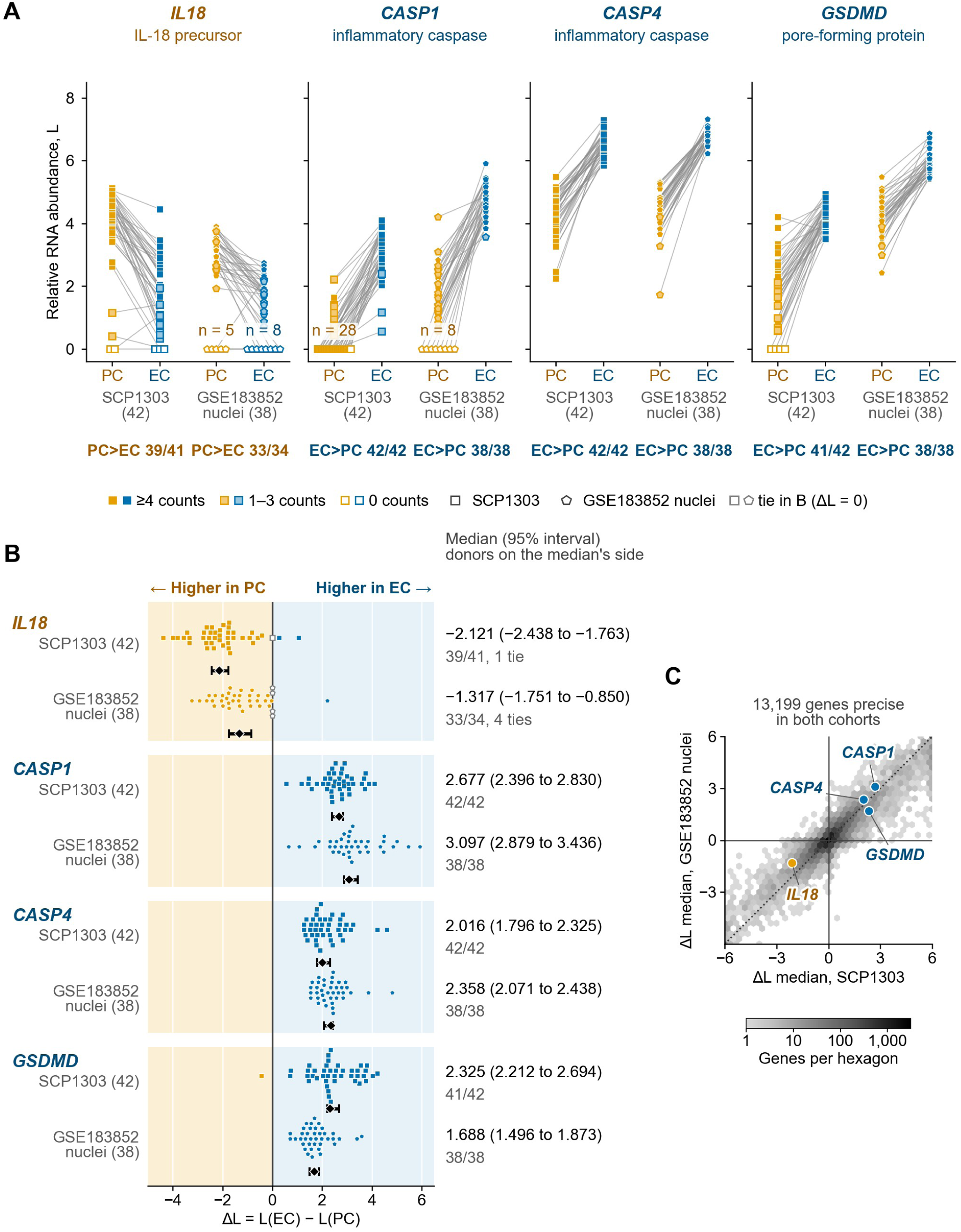
Opposing endothelial–pericyte transcript differences in the two discovery cohorts. (A) Donor-level pericyte (PC) and endothelial (EC) L values in SCP1303 (squares, 42 donors) and GSE183852 nuclei (pentagons, 38 donors). Lines join the two points from the same donor. Fill indicates group counts (≥4, 1–3 or 0). Zero-count points were spread horizontally, with the donor count n added when there were at least five donors. Numbers below each panel give the number of donors on the side of the median and the number of non-tied donors, in the order SCP1303 and GSE183852 nuclei. There were one and four *IL18* ties in the same cohort order. (B) Donor-paired differences, ΔL = L(EC) − L(PC). Points represent individual donors, with grey open points indicating ties at ΔL = 0. Diamonds and horizontal bars show medians and 95% donor-bootstrap intervals. Points were offset vertically to avoid overlap. Medians with 95% intervals and the numbers of donors on that side are shown at right. (C) Median endothelial-minus-pericyte ΔL values for the 13,199 genes meeting the precision criteria in both cohorts. The horizontal axis represents SCP1303 and the vertical axis GSE183852 nuclei. Hexagon intensity indicates gene counts on a logarithmic scale, and the dotted line marks y = x. Medians are taken from the calibration table using 1,000 bootstrap samples, and only the four primary genes are labelled. The 182 genes outside the axis range (±6) were placed at the edges of the plotting area. The four-gene configuration was identified through the exploratory class survey in these two discovery cohorts. EC, endothelium; PC, pericyte.

The other three transcripts favoured endothelium (Figure 2A, 2B). *CASP1* medians were 2.677 (2.396–2.830) in SCP1303 and 3.097 (2.879–3.436) in GSE183852 nuclei, with positive differences in 42/42 and 38/38 donors. *CASP4* medians were 2.016 (1.796–2.325) and 2.358 (2.071–2.438), also positive in every donor. *GSDMD* medians were 2.325 (2.212–2.694) and 1.688 (1.496–1.873), positive in 41/42 and 38/38 donors. The precursor transcript *IL18* and the transcripts for the three molecules involved in its processing and release favoured opposite sides in both cohorts.

Median donor-aggregated *IL18* counts in endothelium and pericytes were 7 and 31 in SCP1303, and 3 and 8 in GSE183852 nuclei. In SCP1303, pericyte *CASP1* had a median count of zero and was detected in 14/42 donors. By contrast, it was detected in all 42 endothelial groups.

### Independent confirmation of opposing endothelial–pericyte enrichment

All four directional predictions from discovery were met in all 22 eligible adult non-failing donors in the prespecified confirmation set. This set comprised 16 Heart Cell Atlas donors and six Reichart donors. *IL18* was relatively enriched in pericytes, whereas *CASP1*, *CASP4* and *GSDMD* were relatively enriched in endothelial cells (Figure 3A).

**Figure 3.**
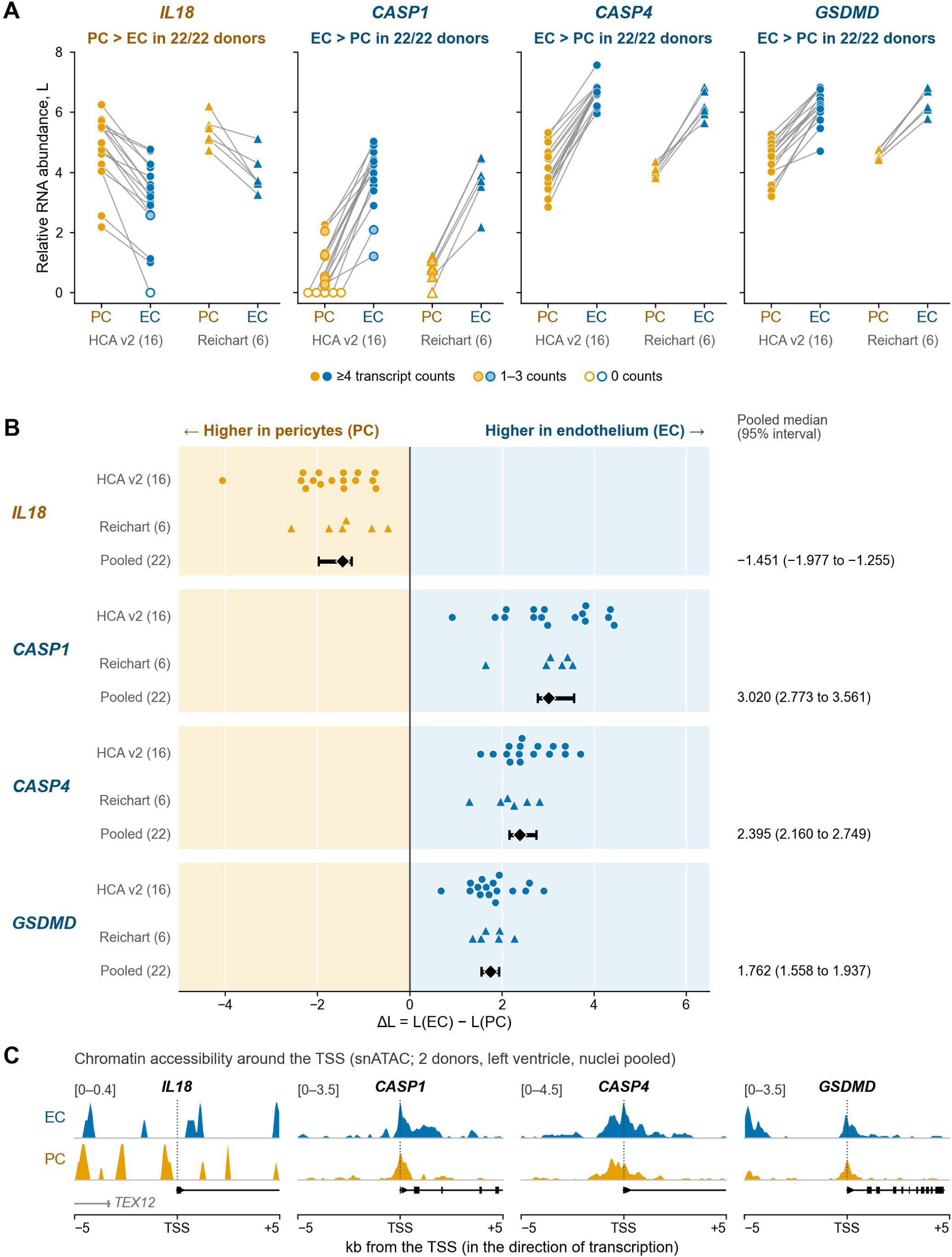
Donor-level confirmation of the opposing endothelial–pericyte configuration and chromatin accessibility around transcription start sites in two separate donors. (A) Class-relative RNA abundance of *IL18*, *CASP1*, *CASP4* and *GSDMD* in the 22 primary confirmation donors, comprising 16 Heart Cell Atlas v2 donors and six Reichart donors. Lines join the pericyte (PC) and endothelial (EC) values of each donor. After donor-by-class pseudobulking, L = log2(1 + 10⁶ × gene count / total count across all available features). Point fill indicates the aggregated count of the donor–class group. Zero-count *CASP1* pericyte points in Heart Cell Atlas v2 were spread horizontally for visibility. Circles represent Heart Cell Atlas v2 and triangles Reichart. (B) Donor-paired differences, ΔL = L(EC) − L(PC), for the same 22 donors on one shared axis. Points represent individual donors. Within each row, points were offset vertically to avoid overlap. Diamonds and horizontal bars show medians across the pooled 22 donors and percentile 95% bootstrap intervals obtained by resampling donors. All four genes met the prespecified pooled confirmation criteria, and all 22 donors showed the predicted direction. Dataset-specific medians and intervals are provided in Supplementary Table S2 and Supplementary Figure S7. (C) In a check planned after confirmation, single-nucleus ATAC data from the left ventricles of two ENTEx donors separate from those in the RNA analysis were used to display accessibility within 5 kb on either side of each gene’s transcription start site, with endothelial and pericyte nuclei pooled separately. Tracks show fragment overlap counts per million total fragments in each class, in 50-base intervals with a 150-base moving average (Supplemental Methods). The vertical-axis range is shared by the two tracks for each gene and is indicated at the upper left. The range for *IL18* is approximately one-tenth that for the other three genes, and most peaks represent 1–2 fragments. Vertical dotted lines mark transcription start sites. MANE Select gene structures are shown below, with boxes indicating exons and arrows indicating transcriptional direction. The grey feature to the left of *IL18* is the neighbouring gene *TEX12*, which is transcribed in the opposite direction. The comparison across six donors is shown in Supplementary Figure S16. EC, endothelium; HCA v2, Heart Cell Atlas v2; PC, pericyte; snATAC, single-nucleus ATAC; TSS, transcription start site.

The pooled median endothelial-minus-pericyte ΔL was −1.451 for *IL18* (95% bootstrap interval, −1.977 to −1.255), 3.020 for *CASP1* (2.773–3.561), 2.395 for *CASP4* (2.160–2.749) and 1.762 for *GSDMD* (1.558–1.937). Each gene met all prespecified confirmation criteria, and all dataset-specific medians were in the predicted direction. The complete four-direction configuration was therefore confirmed in the pooled set of 22 donors (Figure 3B).

Dataset-specific estimates were concordant. The Heart Cell Atlas medians for *IL18*, *CASP1*, *CASP4* and *GSDMD*, in that order, were −1.565, 2.957, 2.416 and 1.762. The corresponding Reichart medians were −1.419, 3.174, 2.194 and 1.792. Every donor within each dataset had a difference in the predicted direction, and all dataset-specific intervals lay wholly on that side. Complete estimates and marginal bootstrap intervals are provided in Supplementary Table S2 and Supplementary Figure S7.

In checks planned after confirmation, all four directions were also preserved in the 22 confirmation donors after exclusion of nuclei with suspected contamination and in each cohort stratified by sex (Supplementary Tables S5, S6).

In another check planned after confirmation, single-nucleus ATAC data from six non-failing heart donors different from those in the RNA analysis were used to compare chromatin accessibility around transcription start sites in endothelium and pericytes (Figure 3C, Supplementary Figure S16). In ventricles, *CASP1*, *CASP4* and *GSDMD* each showed greater accessibility in endothelium in five of six donors, and their median differences ranked in the top approximately 5% of reference genes (Supplementary Figure S16). *IL18* favoured pericytes in five of six donors, but this bias was not pronounced relative to reference genes. Across the gene body, *CASP1*, *CASP4* and *GSDMD* showed greater accessibility in endothelium in all six donors, whereas *IL18* favoured pericytes in two of the six. The region around the *IL18* transcription start site was more accessible in macrophages than in either endothelium or pericytes in the same six donors. In the descriptive comparison of RNA and gene-body accessibility differences measured in different donors, *IL18* was among the genes with the largest discrepancies (top approximately 2%), with RNA more pericyte-directed than expected from accessibility. The median RNA difference for genes with macrophage-specific accessibility was zero, matching the median across all genes, with no indication of macrophage RNA preferentially entering pericyte profiles. The two tiles with pericyte-specific accessibility within 100 kb on either side of *IL18* were both upstream of the transcription start site and overlapped neighbouring genes (Supplementary Table S10).

### The four directions are also preserved in prespecified secondary analyses

All four predicted directions were preserved in all 22 primary confirmation donors both when the endothelial comparison was restricted to capillary states and when predefined ‘-like’ states were excluded (Figure 4, Supplementary Figures S1, S2). The same four directions were observed in all 22 donors when differences in the fraction of transcript-positive nuclei were used instead of ΔL (Supplementary Figure S6).

**Figure 4.**
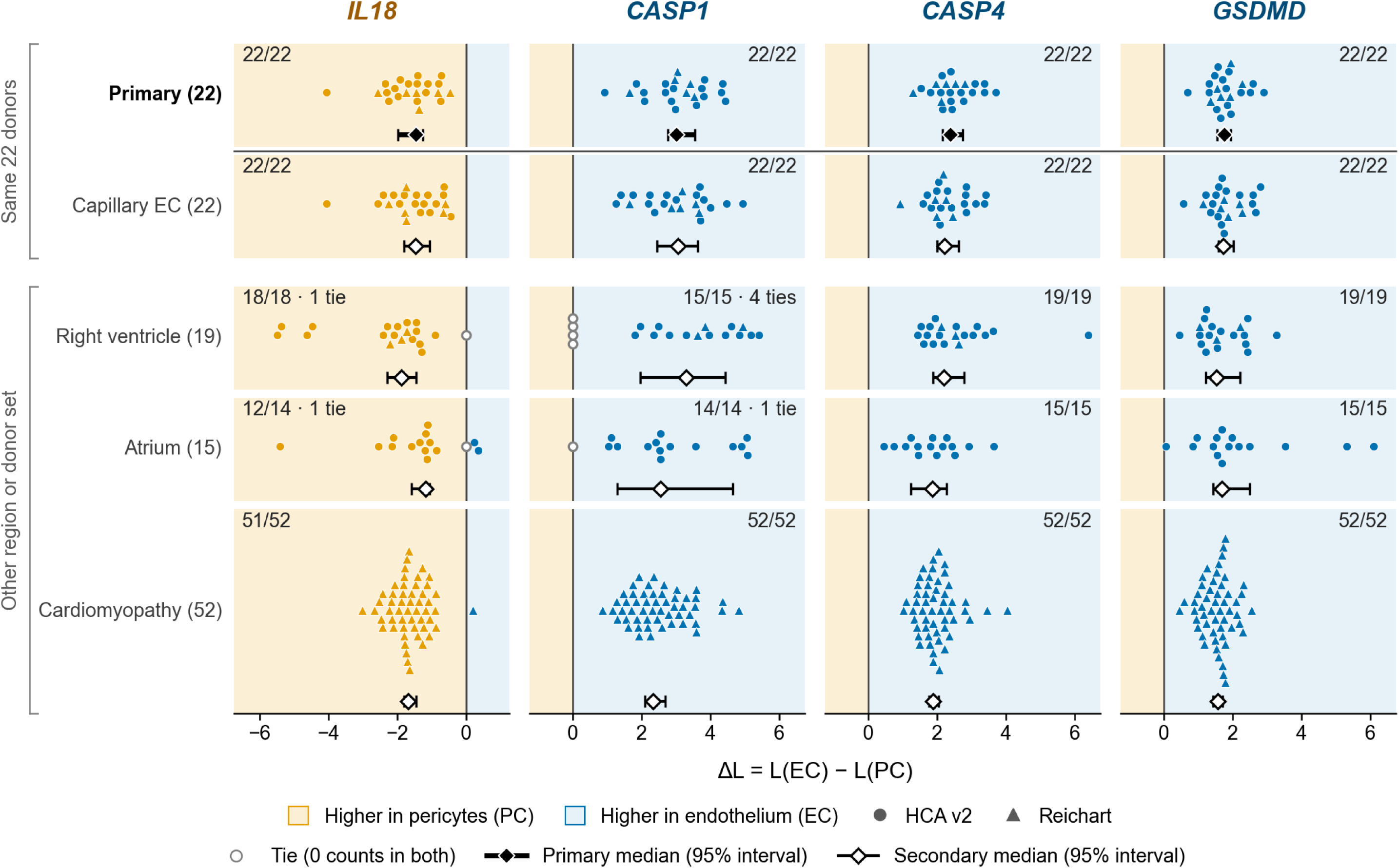
Directions of the four contrasts in prespecified secondary analyses. Each panel shows donor-paired differences, ΔL = L(EC) − L(PC), for one analysis and one gene, with L defined as in Figure 3. Points represent individual donors and are filled orange for negative differences and blue for positive differences. Points were offset vertically to avoid overlap. Diamonds and horizontal bars show medians and 95% donor-bootstrap intervals. Numbers give the number of donors in the predicted direction divided by the number of non-tied donors. The x-axis scale is shared within each gene column. Analyses excluding predefined ‘-like’ states and differences in the fraction of nuclei with detected transcripts are shown in Supplementary Figures S2 and S6. EC, endothelium; HCA v2, Heart Cell Atlas v2; PC, pericyte.

In the right ventricular analysis, the pooled medians from 19 donors preserved all four directions. At the donor level, *IL18* had 18 negative contrasts and one tie. *CASP1* had 15 positive contrasts and four ties, and *CASP4* and *GSDMD* were positive in all 19 donors. In the atrial analysis of 15 Heart Cell Atlas donors, *IL18* was negative in 12, positive in two and tied in one. *CASP1* was positive in 14 with one tie, and *CASP4* and *GSDMD* were positive in all 15 donors. All four atrial median directions also matched the predictions (Figure 4, Supplementary Figures S3, S4).

In the prespecified 52-donor Reichart cardiomyopathy set, *IL18* favoured pericytes in 51 donors and endothelium in one, with a median ΔL of −1.693 (95% bootstrap interval, −1.802 to −1.466). *CASP1*, *CASP4* and *GSDMD* favoured endothelium in all 52 donors, with median differences of 2.347 (2.099–2.683) for *CASP1*, 1.893 (1.766–2.052) for *CASP4* and 1.574 (1.435–1.716) for *GSDMD* (Figure 4, Supplementary Figure S5).

### The survey across cell classes shows the highest relative *IL18* abundance in macrophages

Across the 528 cross-cohort records for matched class pairs and genes, the discovery rules classified 65 as supported, 63 as not supported and 400 as undeterminable. Class-relative RNA abundance and eligible donor numbers for the eight transcripts varied substantially across classes (Figure 5).

**Figure 5.**
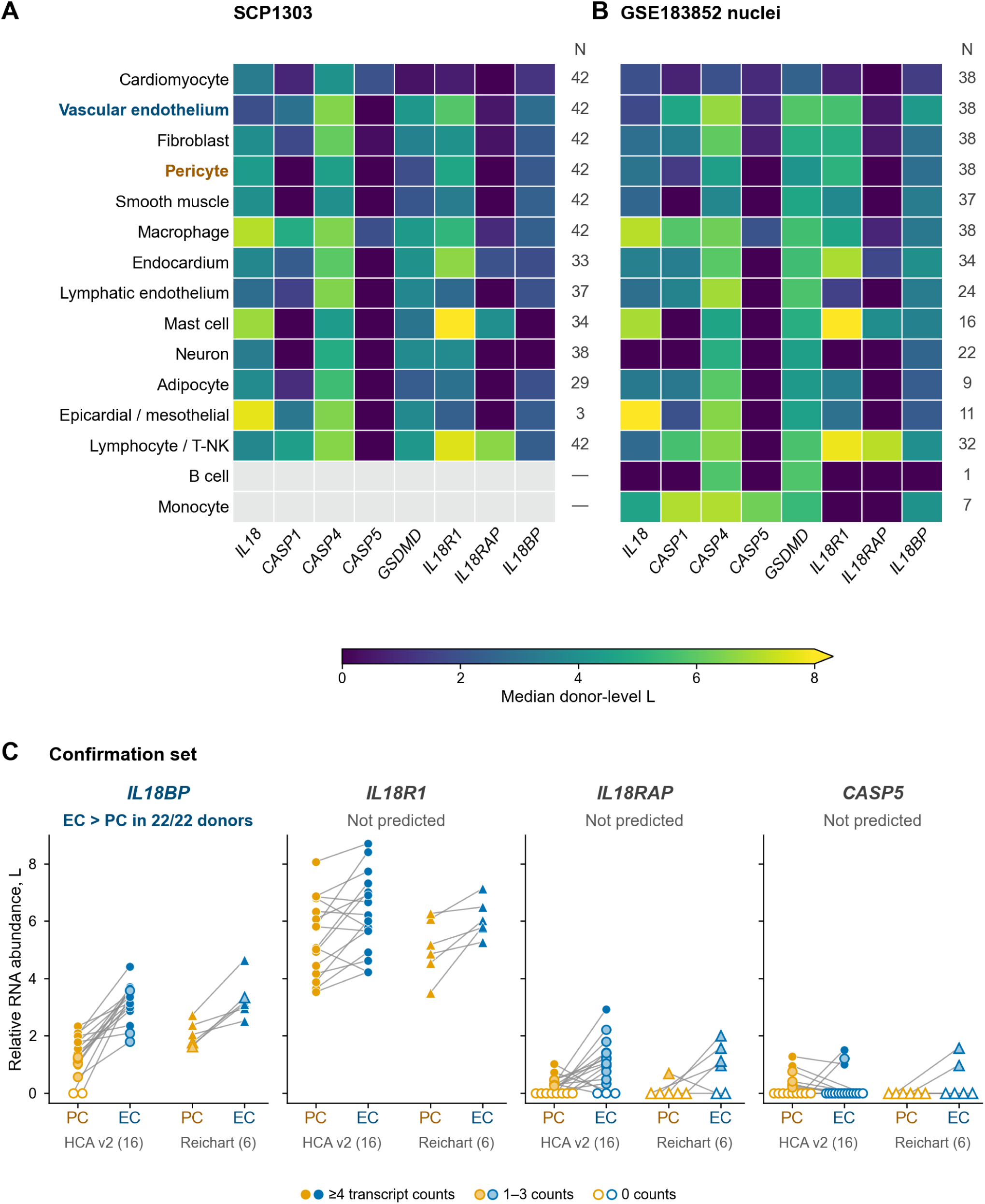
Class-relative RNA abundance in the discovery cohorts and the remaining survey transcripts in the confirmation set. (A, B) Heatmaps show the median donor-level L values for eight transcripts in eligible donor–class groups in SCP1303 (A) and GSE183852 nuclei (B). N beside each row is the number of eligible donors, including donors with zero target counts. Grey denotes no corresponding eligible summary. The colour scale ends at L = 8. The three cells above 8, comprising mast-cell *IL18R1* in both cohorts and epicardial/mesothelial *IL18* in GSE183852, are shown in the top colour. Lymphocyte and T/NK rows are shown as annotated in the sources and were treated separately from the 12 classes matched across cohorts. (C) Pericyte (PC) and endothelial (EC) L values for *IL18BP*, *IL18R1*, *IL18RAP* and *CASP5* in the 22 confirmation donors, comprising 16 Heart Cell Atlas v2 donors and six Reichart donors, with L defined as in Figure 3. Lines join the two values from each donor. Circles represent Heart Cell Atlas v2 and triangles Reichart. Point fill indicates the aggregated count of the donor–class group. Zero-count points were spread horizontally for visibility. The vertical axis is shared by all four transcripts. A prediction was made only for *IL18BP*, and the other three transcripts were evaluated descriptively. EC, endothelium; HCA v2, Heart Cell Atlas v2; PC, pericyte.

Macrophages had the highest measured class-relative *IL18* abundance among the eligible candidate classes in 26/42 SCP1303 and 27/38 GSE183852 donors.

### Calibration shows common individual differences and a limited frequency of all four directions occurring together

For the endothelial–pericyte contrast, 13,199 of the 21,932 genes shared by both cohorts met the precision criteria in both cohorts (Figure 2C). Among these, 4,272 (32.37%) had supported differences: 1,768 favoured pericytes and 2,504 favoured endothelium. This supported set represented 19.48% of all common genes. Individual class differences were therefore common.

Within precision-eligible reference sets matched to the expression of each selected transcript, the specified direction was supported for 39/186 *IL18*-matched genes (20.97%), 51/228 *CASP1*-matched genes (22.37%), 233/958 *CASP4*-matched genes (24.32%) and 41/162 *GSDMD*-matched genes (25.31%). Distinct reference genes were sampled into four labelled slots corresponding to *IL18*, *CASP1*, *CASP4* and *GSDMD*. Simultaneous support for the specified direction in every slot occurred in 608 of 200,000 draws (0.304%). Individual directions were common among reference genes, whereas combinations with all four directions occurring together were infrequent. Support classifications for the four selected transcripts were the same with 1,000 and 10,000 bootstrap samples.

### Sensitivity analyses and comparisons with macrophages and smooth muscle

All four median directions were preserved in each disease stratum, every leave-one-donor-out analysis, the 50- and 100-nucleus thresholds, and the alternative library denominators. In the non-failing subsets, the *GSDMD* interval in SCP1303 and the *IL18* interval in GSE183852 exceeded the allowed width of 1.0. Smaller disease strata failed the minimum donor criterion (Supplementary Figure S11A).

Thinning each donor’s larger class library to the expected depth of the smaller class library preserved the *IL18* median direction in every replicate in both cohorts. *IL18* in GSE183852 met the single-cohort support criteria in 111 of 200 replicates (55.5%), and the remaining replicates were undeterminable (Supplementary Figure S11C). *CASP1*, *CASP4* and *GSDMD* met the criteria in all 200 replicates in both cohorts. Under the more stringent setting that thinned all groups to the cohort’s smallest library, the median *IL18* ΔL in GSE183852 was zero in 116 of 200 replicates.

The five selected macrophage markers were sparsely detected in SCP1303 pericyte groups and more broadly detected in GSE183852. The Spearman correlation between pericyte *IL18* and mean marker abundance was 0.012 in SCP1303 and 0.263 in GSE183852. None of the 34 association tests was significant after correction. *IL18* favoured pericytes even in SCP1303, where marker detection was sparse.

*IL18* also favoured smooth muscle over endothelium. Endothelial-minus-smooth-muscle medians were −1.943 in SCP1303 and −0.667 in GSE183852. Pericyte-minus-smooth-muscle medians were 0.303 and 0.320, and these contrasts were classified as not supported under the discovery rules (Supplementary Figure S11B). In both cohorts, the differences between pericytes and smooth muscle were smaller than the differences between these mural cells and endothelium.

### *GSDMD* also favours endothelium in comparisons with cardiomyocytes

The endothelial-minus-cardiomyocyte *GSDMD* contrast was positive in 42/42 SCP1303 donors, with a median of 3.738 log2(CPM + 1) units and an exact 95% CI of 3.430–3.995. In GSE183852 nuclei, it was positive in 38/38 donors, with a median of 3.272 and an exact 95% CI of 3.089–3.540. In the GeoMx spatial dataset GSE271676, 27/32 donors had a positive difference, with a median of 1.591 upper-quartile-normalised z-score units and an exact 95% CI of 0.666–2.044 (Supplementary Figure S12, Supplementary Table S4). At the protein level, Deep Visual Proteomics data from the Human Protein Atlas for one healthy human heart showed higher GSDMD abundance at all five sites in the endothelial-rich capillary compartment than at all five sites in the cardiomyocyte compartment, with an approximately 22-fold difference between the medians (Supplementary Figure S12E).

When genes in each cohort were ordered by their signed endothelial-minus-cardiomyocyte contrasts, *GSDMD* ranked at the 94.9th, 95.1st and 97.4th percentiles (Supplementary Figure S12A–C). Many other genes also had positive contrasts or complete donor concordance. In the GeoMx composition diagnostic, the *GSDMD* difference was associated with the myeloid marker score (coefficient 0.492, 95% CI 0.097–0.887, Supplementary Figure S12D). In the same diagnostic, the interval for the measured-admixture sensitivity intercept included zero.

### Among the remaining survey transcripts, *IL18BP* also favours endothelium in the confirmation set

The survey transcripts that had not been evaluated in confirmation were *CASP5*, *IL18R1*, *IL18RAP* and *IL18BP*, of which only *IL18BP* had a discovery endothelial–pericyte record classified as supported. In discovery, *CASP5* and *IL18R1* were undeterminable, whereas *IL18RAP* was not supported. *IL18BP* favoured endothelium in both cohorts, with median ΔL values of 0.555 in SCP1303 and 1.548 in GSE183852 nuclei.

In an evaluation planned after the initial confirmation, *IL18BP* was assessed in the confirmation set with a prediction towards endothelium. It favoured endothelium in all 22 donors, with a pooled median ΔL of 1.575 (95% bootstrap interval, 1.377–1.867), and met the prespecified confirmation criteria (Figure 5C). Dataset-specific medians were 1.584 in the 16 Heart Cell Atlas donors and 1.409 in the six Reichart donors. Among the three transcripts without directional predictions, *IL18R1* favoured endothelium in 17/22 donors and *IL18RAP* in 17/21 non-tied donors, whereas *CASP5* was tied in 11/22 donors (Figure 5C, Supplementary Table S7).

In secondary analyses, *IL18BP* favoured endothelium in 21/22 donors in the capillary endothelial–pericyte comparison and in 43/52 Reichart cardiomyopathy donors (Supplementary Table S7). In the right ventricle, it favoured endothelium in 12/18 non-tied donors, with a pooled median of 1.477 (95% bootstrap interval, −0.240 to 2.706) (Supplementary Table S7).

In a single-nucleus ATAC check planned after confirmation, ventricles from six donors different from those in the RNA analysis showed greater accessibility around the *IL18BP* transcription start site in endothelium in four of six donors, with a median accessibility difference of 0.20 in the top approximately 19% of reference genes (Supplementary Figure S16). The criterion for an endothelial direction, specified before the values were counted, required at least five of six donors on the endothelial side and a positive median difference. Under this criterion, accessibility around the *IL18BP* transcription start site could not be classified as endothelial-biased. When all sampled heart regions were included, two of six donors favoured endothelium.

### Other pyroptosis-related genes

In an evaluation planned after the initial confirmation and the assessment of the remaining survey transcripts, we examined ten other pyroptosis-related genes in the confirmation set. These were the inflammasome sensors *NLRP1*, *NLRP3*, *NLRC4*, *AIM2*, *CARD8* and *MEFV*, the adaptor *PYCARD*, the substrate *IL1B*, and *GSDME* and *NINJ1*, which are involved in executing cell death. Of these, only *CARD8* had a discovery endothelial–pericyte record classified as supported, favouring endothelium in both cohorts, with medians of 2.646 in SCP1303 and 2.881 in GSE183852 nuclei. When the prediction of endothelial enrichment was evaluated in the confirmation set, *CARD8* favoured endothelium in all 22 donors, with a pooled median ΔL of 2.728 (95% bootstrap interval, 2.564–3.001), and met the prespecified confirmation criteria (Supplementary Table S9). Among the nine genes without directional predictions, the sensors *NLRP1* and *AIM2* and the adaptor *PYCARD* favoured endothelium, whereas *GSDME* favoured pericytes, in nearly all non-tied donors. Intervals for *NLRC4* and *NINJ1* included zero, whereas *NLRP3*, *MEFV* and *IL1B* had zero counts in many groups (Supplementary Table S9).

### Comparisons of deposited experimental structures distinguish IL-18 maturation, IL-18BP sequestration and receptor assembly

In a comparison planned after the transcript analyses, we applied the same quantitative rules to deposited experimental IL-18 structures to relate transcript enrichment in the confirmation set to the molecular stages involving IL-18 (Figure 7, Supplementary Table S11). IL-18BP and IL-18Rα (the product of *IL18R1*) shared some recognition residues on mature IL-18. With contacts defined by protein heavy-atom distances of less than 4.5 Å, IL-18BP (PDB 7AL7) [41] contacted 23 IL-18 residues and IL-18Rα (PDB 3WO4) [42] contacted 40, with 16 shared residues (Figure 7C). Superposition of the two structures using the 157 Cα atoms of mature IL-18 gave a root-mean-square deviation (RMSD) of 1.27 Å. In this hypothetical overlay, 1,199 atom pairs between IL-18BP and IL-18Rα were less than 2 Å apart, and the proteins overlapped substantially in their respective binding poses. This result is consistent with the report that IL-18BP interferes with receptor recognition [41].

The mature IL-18 backbone changed little across the stages of receptor assembly. We superposed the structures onto IL-18 in 3WO4, which also contained IL-18Rβ (the product of *IL18RAP*), using the 156 Cα atoms shared by all copies. The RMSD values were 0.66–0.74 Å for IL-18 alone (four copies in PDB 3WO2) and 0.37–0.45 Å for the IL-18–IL-18Rα complex (six copies in PDB 3WO3) [42]. In 3WO4, IL-18Rβ contacted both IL-18 and IL-18Rα: of its contact residues, 6 contacted only IL-18, 15 contacted only IL-18Rα and 4 contacted both (Figure 7C).

In contrast, the arrangement of the same recognition residues differed before and after maturation. For IL-18BP recognition residues, the median root-mean-square difference in pairwise Cα distances relative to the bound structure was 12.9 Å in pro-IL-18 (10 NMR models in PDB 8URV) [43] and 1.9 Å in mature IL-18 (20 NMR models in PDB 1J0S) [44]. The corresponding precursor and mature values were 8.8 Å and 2.8 Å for IL-18Rα recognition residues and 7.2 Å and 1.8 Å for IL-18Rβ recognition residues. When restricted to residues within the well-ordered ranges reported for the precursor, the precursor and mature values were 3.2 Å and 0.8 Å for IL-18BP, 6.2 Å and 1.5 Å for IL-18Rα, and 6.0 Å and 0.9 Å for IL-18Rβ (Figure 7C). In pro-IL-18, atoms in the pro-region covered part of the solvent-accessible surface area of these recognition residues (Supplementary Table S11). This result is consistent with the report that receptor- and IL-18BP-binding sites form after the structural changes accompanying cleavage [43].

### All four median directions are preserved in ischaemic hearts, and the endothelial–pericyte *CASP4* difference is smaller and pericyte *CASP4* higher in acute myocardial infarction than in controls

In all five acute myocardial infarction (AMI) and ischaemic cardiomyopathy (ICM) groups across the three cohorts GSE270788, Kuppe and SCP1849, median ΔL values were positive for *CASP1*, *CASP4* and *GSDMD* and negative for *IL18*, matching the four predicted directions. The ischaemic-heart analyses were planned as exploratory analyses after confirmation, and the five groups were compared with their respective cohort controls (Figure 6A). The prespecified criterion assessed whether at least 75% of non-tied donors showed the predicted direction. This criterion was met in all five groups for *CASP4*, in the four groups other than SCP1849 ICM for *CASP1* and *GSDMD*, and only in Kuppe ICM for *IL18*.

**Figure 6.**
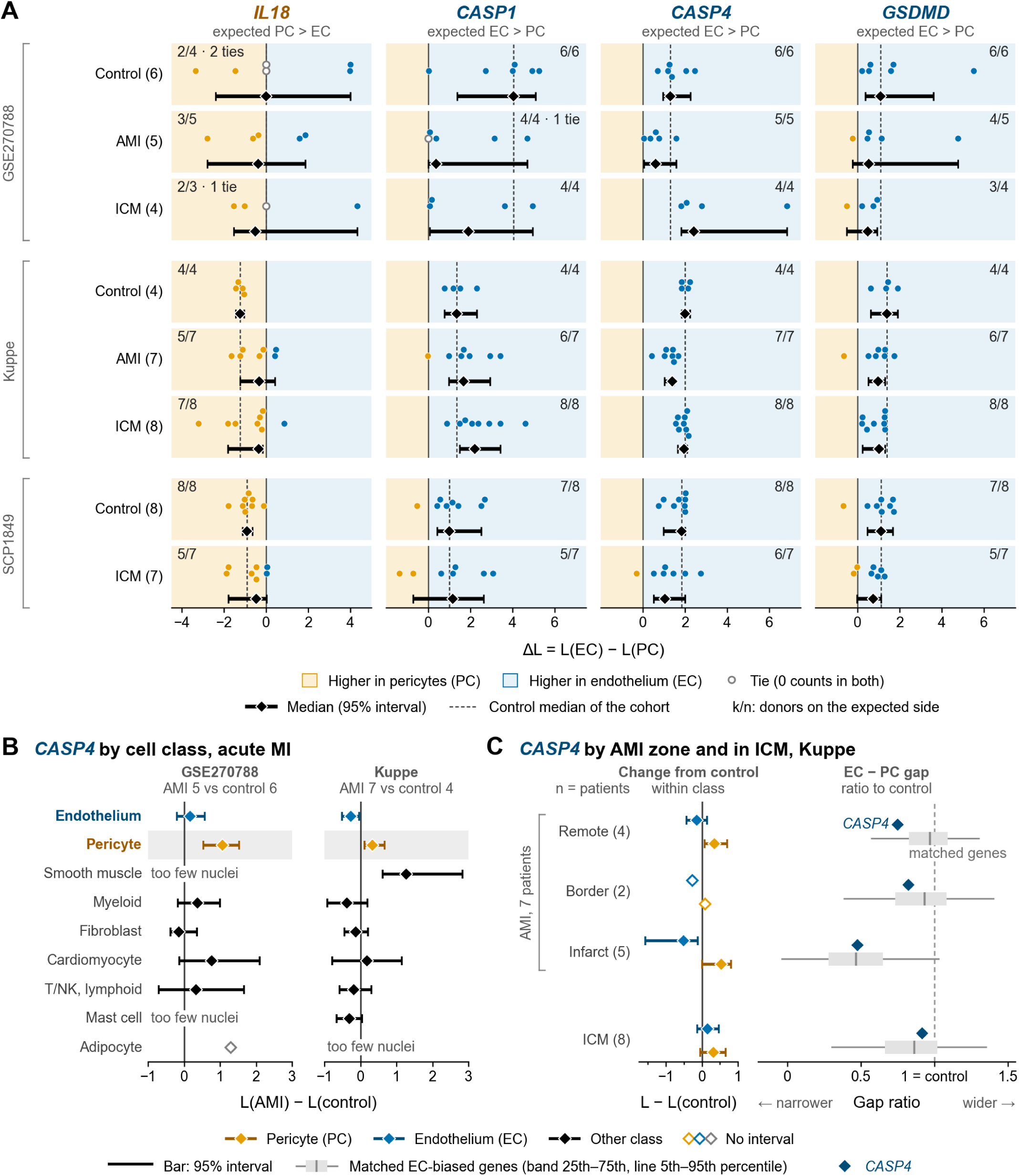
Four endothelial–pericyte contrasts in ischaemic hearts and class- and region-level changes in *CASP4*. (A) Donor-paired differences, ΔL = L(EC) − L(PC), for *IL18*, *CASP1*, *CASP4* and *GSDMD* in three ischaemic-heart cohorts, with L defined as in Figure 3. Numbers in parentheses in the row labels indicate eligible donor counts. Points represent individual donors and are filled orange for negative differences and blue for positive differences. Points were offset vertically to avoid overlap. Black diamonds and horizontal bars show medians and 95% bootstrap intervals. The same resamples were used for all four genes. The k/n annotation in the upper corner of each panel gives the number of donors in the predicted direction divided by the number of non-tied donors. Predicted directions are those in Figures 2 and 3. One horizontal-axis unit has the same plotted length across all four columns, with only the *IL18* column shifted towards negative values. (B) Class-level changes in *CASP4* in acute myocardial infarction, L(AMI) − L(control), using count-pooled L (Methods). Donor counts above the panels refer to the endothelial and pericyte rows, which include the same donors as A. Counts for other classes may differ according to class eligibility. The pericyte row has a pale grey band. (C) *CASP4* in the AMI regions and ICM in Kuppe. Numbers in parentheses in the row labels give the number of patients eligible for both endothelium and pericytes. The same patient can contribute to multiple rows. The left part shows class-level changes in count-pooled L relative to controls. The right part shows the contrast ratio κ, the endothelial–pericyte difference calculated from count-pooled L relative to the control difference (Methods). The grey band and lines show the distribution of κ for 307 endothelial-biased reference genes. The vertical line within the band marks the median. Selection of the endothelial-biased reference genes is described in Supplemental Methods. The same genes were used in all four rows. One horizontal-axis unit has the same plotted length in B and the left part of C. AMI, acute myocardial infarction; EC, endothelium; ICM, ischaemic cardiomyopathy; MI, myocardial infarction; PC, pericyte; T/NK, T and NK cells.

The endothelial–pericyte *CASP4* difference was smaller in AMI than in controls in both GSE270788 and Kuppe (Figure 6A). Using counts pooled by class across donors, the AMI-minus-control difference in count-pooled ΔL was −0.907 (95% bootstrap interval, −1.352 to −0.483) in GSE270788 and −0.597 (−0.978 to −0.358) in Kuppe (Supplementary Figure S13A). The smaller contrast in AMI was accompanied by higher pericyte *CASP4* (Figure 6B). Pericyte *CASP4* was higher in AMI than in controls, with differences of 1.074 (0.537–1.526) in GSE270788 and 0.337 (0.117–0.668) in Kuppe. Pericytes were the only class with an interval above zero in both cohorts. The AMI-minus-control changes in endothelial *CASP4* were 0.167 (−0.199 to 0.578) in GSE270788 and −0.260 (−0.512 to −0.028) in Kuppe. The smooth-muscle change was available only in Kuppe and was 1.269 (0.614–2.839).

We compared the *CASP4* changes with those of genes with pericyte abundance close to *CASP4*. In these sets of 2,364 genes for GSE270788 and 3,066 for Kuppe, 9.31% and 17.58% had pericyte AMI-minus-control differences at least as large as that of *CASP4*, and 7.06% and 3.52% had reductions in the endothelial–pericyte difference at least as large as that of *CASP4* (Supplementary Figure S13A).

In Kuppe, AMI specimens were divided into remote myocardium, border zone and infarct zone and compared with controls alongside ICM (Figure 6C). In remote myocardium, pericyte *CASP4* was 0.352 (0.079–0.699) higher than in controls, whereas the endothelial change was −0.149 (−0.429 to 0.143) (Supplementary Figure S13B). The endothelial–pericyte *CASP4* difference in remote myocardium was 0.751 times the control difference. We compared this contrast ratio with those of 307 genes that had an endothelial bias of at least 1 in controls and abundance close to *CASP4*. The median contrast ratio for these endothelial-biased genes was 0.970. There were 53 genes (17.26%) with ratios equal to or smaller than that of *CASP4*.

The reduction in the *CASP4* difference was greatest in the infarct zone, with a contrast ratio of 0.477. Endothelial-biased genes showed a similar reduction in the infarct zone, with a median contrast ratio of 0.468. There were 157 genes (51.14%) with ratios equal to or smaller than that of *CASP4*. In Kuppe ICM, the *CASP4* contrast ratio was 0.918, close to the control value, and the median ratio for endothelial-biased genes was 0.864. At the patient level, the *CASP4* ratio was smaller than the median ratio for endothelial-biased genes in the same patient in 4/4 patients in remote myocardium and 2/5 in the infarct zone (Supplementary Figure S13C). In one of these patients in remote myocardium, the *CASP4* ratio was 0.862 and the median ratio for that patient’s endothelial-biased genes was 0.864.

We used three approaches to check whether higher pericyte *CASP4* resulted from contamination by other cells (Supplementary Figure S14). Expressed in CPM, the observed AMI-minus-control difference in pericyte *CASP4* was 54.9 CPM in GSE270788 and 5.4 CPM in Kuppe (Supplementary Figure S14A). After subtracting marker-estimated contamination using myeloid nuclei as the contamination source, the remaining differences were 50.2 CPM (26.1– 86.9) in GSE270788 and 5.2 CPM (1.8–9.7) in Kuppe. Using all nuclei with public annotations as the source gave remaining differences of 49.7 CPM (26.4–100.3) and 3.0 CPM (−6.7 to 13.6). Contributions from four classes, myeloid cells, endothelium, fibroblasts and cardiomyocytes, were also estimated using genes specific to each class. In Kuppe, three classes were used, excluding the myeloid class because it had fewer than five specific genes. The AMI-minus-control differences expected from contamination by these classes were 47.3 CPM in GSE270788 and 2.6 CPM in Kuppe. Subtracting these contributions left 7.6 CPM (−68.8 to 45.1) and 2.8 CPM (−3.4 to 8.9) (Supplementary Figure S14B). Excluding nuclei bearing myeloid or endothelial markers (9.85–12.13% of pericyte nuclei) changed the AMI-minus-control difference from 1.074 to 0.694 (0.039–1.303) in GSE270788 and from 0.337 to 0.327 (0.107–0.699) in Kuppe (Supplementary Figure S14C). Leaving out one donor at a time before nucleus exclusion yielded estimates of 0.978–1.241 across the 11 GSE270788 analyses and 0.262–0.440 across the 11 Kuppe analyses, all positive. Under the prespecified interpretation rules, the checks gave mixed results.

For *IL18*, *CASP1* and *GSDMD*, no class had an AMI-minus-control interval entirely above zero in both cohorts or entirely below zero in both cohorts (Supplementary Figure S15A). Pericyte intervals included zero for all three genes in both cohorts. Myeloid cells accounted for the largest share of *IL18* counts in GSE270788 (Supplementary Figure S15B). In Kuppe, cardiomyocytes accounted for the largest share, followed by myeloid cells. Endothelium in GSE270788 here excludes endocardium and lymphatic endothelium. Pericytes and endothelium each accounted for less than 5% in all four groups.

## Discussion

In the cell classes forming the human cardiac microvasculature, *IL18*, which encodes the IL-18 precursor, was relatively enriched in pericytes, whereas the transcripts *CASP1* and *CASP4*, encoding processing enzymes, and *GSDMD*, encoding the pore-forming molecule, were relatively enriched in endothelium. All four directions of this configuration, identified by paired comparisons within donors, were reproduced in all 22 separate donors under prespecified confirmation rules. On the basis of these opposing transcript distributions, we propose the configuration hypothesis of intercellular division of labour: pericytes supply the IL-18 precursor and endothelium carries out its processing and release.

In this hypothesis, the precursor supplied by pericytes is transferred to endothelium, where it is processed and released in its mature form. The reaction depends on the relationship between the cells supplying the precursor and those carrying out processing and release. Limited IL-18 secretion despite clear GSDMD cleavage has been reported in cultured human endothelial cells [40], giving reason to distinguish the action of molecules involved in processing and release from the supply of the precursor substrate. The experimental finding that pro-IL-18 released from human keratinocytes is activated extracellularly by the bacterial protease SpeB [22] also supports examining processes that link precursor production and maturation at different sites. Routes through which parts of a reaction pass between cells have been demonstrated experimentally: inflammatory responses propagate through membrane nanotubes connecting cardiac cells [9], and vesicles carry active caspase-1 or GSDMD pores [28,29]. These routes are candidates for mediating the proposed transfer.

The configuration hypothesis makes a concrete prediction: transfer between cells is itself a point at which the reaction is regulated. Even with the same precursor abundance and the same state of the processing side, generation of mature IL-18 is predicted to change if the conditions allowing the precursor to reach the processing side change. Thus, alongside altering molecular expression, altering transfer between the supplying and processing sides provides a point of intervention in the reaction. In contrast, if a subset of cells in each cell class co-expresses the necessary molecules and completes maturation and release within those cells, the proportion of cells carrying the full set of molecules and their responses to stimulation become important. The two explanations can be distinguished by combining same-cell co-expression measurements, tracing the origin of the precursor, and interventions that separately target the supplying and processing sides.

Another central element of the configuration hypothesis is an endothelial role not only in generating mature IL-18 but also in sequestering it and regulating its action. In the confirmation set, *IL18BP* and *CARD8*, which encodes an inflammasome sensor, also favoured endothelium in all 22 donors, alongside transcripts involved in processing and release, and met the prespecified criteria for prediction-based confirmation (Figure 5C, Supplementary Table S9). Endothelial IL-18BP has been reported in human intestinal submucosal vessels and cultured endothelial cells [34], and an *Il18bp* reporter has been detected in endothelium across several mouse organs, including the heart [35].

CARD8 functions as one of the main inflammasome sensors in cultured human endothelial cells and induces GSDMD cleavage and pyroptosis upon stimulation [40]. The dependence of free IL-18 abundance on its quantitative balance with IL-18BP [36,37] leads to the hypothesis that endothelium participates in both generation and sequestration of mature IL-18, and that the relationship between these processes determines the amount of IL-18 available to the receptor.

Species differences among the relevant molecules also make clear why this molecular configuration needs to be studied in the human heart. Rodents lack a *CARD8* orthologue [38,40], and caspase-11 in mice and rats, which corresponds to human caspase-4, directly cleaves pro-IL-18 only weakly [2]. The molecular composition and function involving *CASP4* and *CARD8* therefore cannot be extrapolated directly from standard mouse models. This study uses the transcript configuration reproduced in humans to ask about intercellular division of labour for IL-18 in the context of the human molecular composition.

Generation of mature IL-18 and its use by the receptor are also distinct molecular stages. In comparisons of deposited experimental structures, the arrangement of the same recognition residues differed between pro-IL-18 and mature IL-18, whereas the mature IL-18 backbone changed little across stages of receptor assembly, and IL-18BP and IL-18Rα shared some recognition residues on mature IL-18 (Figure 7). This comparison quantified the known processes of structural change accompanying maturation, sequestration by IL-18BP and receptor assembly [41–43] using the same rules. Applied to the interpretation of transcript distributions, this distinction identifies what needs to be related: the cells containing the precursor, the sites where the mature form is generated, the sites where it is sequestered, and the sites where it is used by the receptor. The configuration hypothesis seeks to explain the relationships between cells and molecules that link precursor supply to generation of mature IL-18 and determine whether the mature form is sequestered or leads to a receptor response.

**Figure 7.**
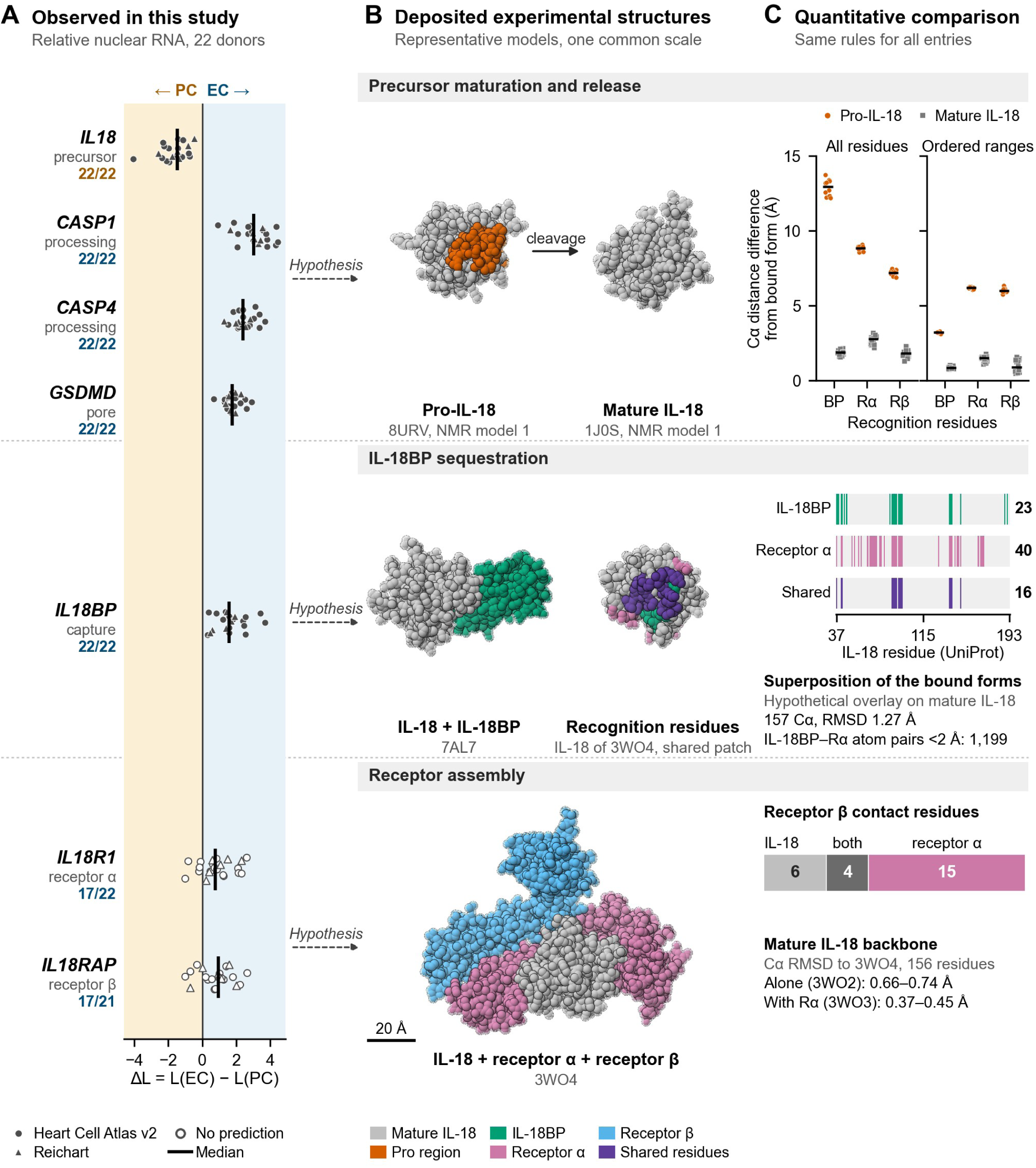
Transcript enrichment in the confirmation set and comparisons of IL-18 maturation, sequestration and receptor assembly in deposited experimental structures. (A) Endothelial-minus-pericyte differences, ΔL, for each transcript in the 22 confirmation donors, comprising 16 Heart Cell Atlas v2 donors and six Reichart donors, grouped by molecular stage, with ΔL defined as in Figure 3. Points represent individual donors, and vertical lines mark medians. Numbers give the donors on the side of the median, with non-tied donors as the denominator. Open symbols denote the two transcripts for which no prediction was made. (B) Deposited experimental structures shown at a common scale. Spheres represent heavy atoms. Grey denotes IL-18 residues 37–193 and orange the pro region, residues 1–36, with other colours indicating binding partners. Colours do not represent RNA abundance. NMR structures are represented by model 1. “Recognition residues” shows mature IL-18 from 3WO4 viewed from the side bearing recognition residues shared by IL-18BP and IL-18Rα. Green marks recognition residues for IL-18BP only, pink those for IL-18Rα only, and purple those shared by both. (C) The upper part shows the root-mean-square difference in pairwise Cα distances between recognition residues relative to the bound structures. Points represent individual models, and horizontal lines mark medians. “Ordered ranges” gives values restricted to residues within the well-ordered ranges reported for the precursor. The middle part shows the positions and numbers of recognition residues on mature IL-18, defined by heavy-atom distances of less than 4.5 Å, together with values from a hypothetical overlay of the IL-18BP and IL-18Rα binding poses superposed on mature IL-18. The lower part shows the breakdown of IL-18Rβ contact residues in 3WO4 and the RMSD of the mature IL-18 backbone. Dashed arrows from A to B indicate hypotheses. BP, IL-18BP; EC, endothelium; PC, pericyte; Rα, IL-18Rα; Rβ, IL-18Rβ.

An important question in ischaemic myocardial injury is how the relationship between the supplying side and the processing side varies with disease. The median directions of the four transcripts were preserved in the exploratory analyses of ischaemic hearts. In both acute myocardial infarction cohorts, however, *CASP4* relative RNA abundance was higher than in controls in the pericyte class, where relative *IL18* abundance exceeded that in endothelium, and the endothelial–pericyte difference was smaller (Figure 6A, 6B). Because activated human caspase-4 can directly process pro-IL-18 [1,2], this finding leads to the hypothesis that the conditions linking the precursor and its processing enzyme within pericytes change with myocardial injury. In Kuppe, the reduction in the *CASP4* contrast relative to controls in remote myocardium exceeded the median reduction for endothelial-biased reference genes, whereas in the infarct zone, the reduction for reference genes was similar to that for *CASP4* (Figure 6C, Supplementary Figure S13). Although both regions show a smaller contrast, the questions differ: the state of individual molecules, including *CASP4*, in remote myocardium, and how the contrast fits within broader differences in transcript distributions in the infarct zone. The relation to disease can be considered not only in terms of whether the direction of the distribution reverses, but also whether the state of processing-related molecules differs on the side relatively enriched in *IL18*.

Viewed in terms of intercellular division of labour, ischaemia-associated changes in the spatial relationship between pericytes and endothelium represent a factor that alters molecular transfer. In rat hearts subjected to ischaemia– reperfusion, capillary constriction by pericytes contributes to impaired reperfusion [23]. In mouse myocardial infarction, pericyte detachment from microvessels and migration to the site of injury have been reported together with increased vascular leakage [45]. If precursor supply, processing and sequestration depend on different cells, this leads to the hypothesis that such changes in the spatial relationship between cells alter precursor transfer and contact with molecules involved in sequestering mature IL-18. Adding IL-18BP improved contractile function after reperfusion in an ex vivo ischaemia–reperfusion model of human atrial tissue [7]. This also supports examining regulation of IL-18 action, alongside generation of the mature form, in relation to myocardial injury. Thus, connecting transcript configuration to microvascular function calls for joint examination of the differences from controls in *CASP4* within each cell class, the relationship between pericytes and endothelium, and sequestration by IL-18BP.

Establishing the relationship between cell classes with relative transcript enrichment and cells that contribute functionally to myocardial injury requires both expression measurements and interventions. Relative *IL18* RNA abundance was higher in smooth muscle as well as pericytes than in endothelium. Among eligible cell classes in the discovery cohorts, macrophages had the highest abundance in more than half of the donors. GSDMD-mediated IL-18 release has been demonstrated experimentally in mouse myeloid cells [24]. By contrast, *GSDMD* also showed relative endothelial enrichment in comparisons with cardiomyocytes (Supplementary Figure S12). Relevant reports describe caspase-4- and caspase-5-dependent cell death and GSDMD cleavage in human pulmonary microvascular endothelial cells [25], and reduction of ischaemia–reperfusion injury by cardiomyocyte-specific GSDMD deficiency in mice [26]. These findings make it important to compare vascular cells, immune cells and cardiomyocytes in terms of both transcript abundance rankings and the stages of precursor supply, processing and release at which intervention changes the reaction.

The next question in understanding how this configuration arises is whether transcription of the genes encoding the precursor, the molecules involved in processing and release, and the inhibitor is regulated in the same way. In the single-nucleus ATAC check planned after confirmation, accessibility around transcription start sites for the genes encoding the three molecules involved in processing and release also favoured endothelium, and the differences ranked in the top approximately 5% of reference genes. By contrast, *IL18* and *IL18BP*, whose RNA consistently favoured opposite sides in all 22 donors, showed accessibility biases that were not pronounced relative to reference genes (Supplementary Figure S16). The region around the *IL18* transcription start site was highly accessible in macrophages, and the discrepancy between RNA and accessibility was also large (Supplementary Table S10). The biases of these two transcripts may arise from differences in transcription rather than accessibility. This hypothesis can be tested by relating chromatin accessibility, newly synthesised RNA and RNA decay rates within each cell class.

The key to testing the configuration hypothesis is to connect precursor origin to receptor responses within the same tissue and experimental system. The starting point for comparison with the reported localisation of IL-18 protein in failing ischaemic hearts [18] is to distinguish endothelial cells, pericytes and macrophages and map *IL18* RNA, precursor and mature IL-18, active caspases and cleaved GSDMD in relation to one another in the same specimens. Combining precursor tracing with interventions directed separately at the supplying and processing sides and changes in the conditions of contact between cells can test whether generation of mature IL-18 depends on transfer between cells. Joint measurement of sequestration by IL-18BP and receptor responses can also distinguish the stage at which mature IL-18 is generated from the stage at which it produces its effects.

The measurements represent relative nuclear RNA abundance aggregated by cell class and may be affected by low counts, cell-state composition, ambient RNA or doublets, and differences in input processing. Independence of the confirmation donors was based on documented provenance rather than genetic matching. Atlas information had been inspected before the confirmation, and upstream processing of the Reichart input remains unresolved. Same-cell co-expression, inflammasome assembly and activation, GSDMD cleavage and pyroptotic cell death, protein maturation and release, neutralisation of IL-18 by IL-18BP, and intercellular transfer were not directly measured. The structural comparison describes the geometry of deposited structures and does not measure affinity, the feasibility of simultaneous binding or protein states in cardiac tissue. The protein reference is limited to two compartments from one donor. The single-nucleus ATAC checks involved six donors separate from those in the RNA analysis, and *IL18* and *IL18BP* signals were sparse.

The ischaemic-heart analyses include assessments added after inspecting results and are affected by small sample sizes, donor overlap with discovery data [27] or unresolved overlap, differences in depth, region and sampling time, shared controls and repeated measurements within patients. The intervals were not adjusted for multiplicity, and the q values for pericyte *CASP4* were 0.24 and 0.25. Because the contamination checks gave mixed results, determining whether the increase is intrinsic to pericytes requires measurements with cell identification in tissue.

We establish opposing endothelial–pericyte enrichment of IL-18-related transcripts in the human heart and propose the configuration hypothesis that pericytes and endothelium divide the tasks of precursor supply, processing and release. This hypothesis views IL-18-related reactions not only in terms of molecular expression within individual cells but also in terms of relationships between cells carrying out different stages. The relationship to ischaemic myocardial injury calls for testing both the abundance of individual molecules and the relationships between the cells that contain them. At the centre of the configuration hypothesis is the question of which combinations of cells and molecules connect precursor supply, maturation and release, sequestration, and use by the receptor.

## Supporting information

Supplementary Software 1

## Statements and Declarations

### Ethics approval

This secondary analysis used publicly available, de-identified data and involved no participant recruitment, intervention or tissue collection. Ethics and consent arrangements are reported by the source studies [11–13,16,17,19–21,32,33], including Washington University Institutional Review Board approval 201104172 for GSE183852.

### Data and code availability

The discovery resources are publicly available under SCP1303 and GSE183852. The GeoMx spatial data used for the endothelial–cardiomyocyte comparison are publicly available under GSE271676. The processed spatial data are available from Mendeley Data, version 1 (doi:10.17632/gmgcdhmmdx.1). Confirmation used the public processed Heart Cell Atlas v2 and Reichart resources [16,17]. File identities, matrix locations, source identifiers and analysis specifications are documented in Supplementary Software 1. Supplementary Data 1 contains donor-level and aggregate results, all 528 records from the discovery survey, confirmation and secondary-analysis outputs including the remaining survey transcripts, the positive-control values in Figure 1D, and the embedding coordinates and bin tables for Figure 1 and Supplementary Figures S8–S10. Analysis code, plotting code, saved resampling indices and provenance are supplied in Supplementary Software 1. The ischaemic-heart analyses used GSE270788 from GEO [19], the Kuppe et al. resources [20] and SCP1849 from the Single Cell Portal [21]. The Kuppe et al. resources comprise processed data from Zenodo (doi:10.5281/zenodo.6578047), single-nucleus ATAC fragments (doi:10.5281/zenodo.6578553), specimen metadata (doi:10.5281/zenodo.6578617) and Supplementary Table 1 of the original study. Processing-batch assignments for GSE270788 were obtained from the publicly available code for the original study (github.com/jamrute/immune-fibroblast-heart-failure). The protein reference used public Deep Visual Proteomics data from the Human Protein Atlas (proteinatlas.org) [30]. The single-nucleus ATAC checks used GSE184462 and GSE165837 from GEO [32,33]. The structural comparisons used PDB entries 7AL7, 3WO2, 3WO3, 3WO4, 8URV and 1J0S [41–44]. The comparison outputs and code are included in Supplementary Data 1 and Supplementary Software 1.

### Author contributions

D.O. conceived the study, designed the analyses, curated the public datasets, performed the computational analyses, generated the figures, interpreted the data and drafted the manuscript. G.D. contributed to study conceptualisation, interpretation of the transcriptomic findings and critical revision of the manuscript. S.F., N.N., K.O., A.K., M.A., Y.K., K.A. and H.N. contributed to clinical, immunological and cardiovascular interpretation of the findings and critically revised the manuscript for important intellectual content. H.M. supervised the study, contributed to study conceptualisation and interpretation, and critically revised the manuscript.

