## Supplementary Information for "*IL18* and pyroptosis-related genes show contrasting expression in human cardiac pericytes and endothelial cells"

#### Supplemental Methods

##### Study design, data sources and units of analysis

Source lymphocyte and T/NK annotations, whose granularity differs, were shown unchanged and separately from the 12 matched classes. B-cell and monocyte annotations available only in GSE183852 were not treated as matched classes.

For a protein reference, we used sample-level Deep Visual Proteomics data from the Human Protein Atlas [30]. The file dvp_sample_data.tsv was retrieved on 23 August 2026. We read GSDMD and CASP4 intensities in the two cardiac compartments, capillaries and cardiomyocytes, as provided in the source table. The five values per compartment represent samples from five sites within one donor. The values were summarised descriptively.

##### Shared embedding of endothelial and pericyte nuclei

For display, endothelial and pericyte nuclei used in the donor-level comparisons were placed in one shared two-dimensional embedding per cohort. The 42 SCP1303 donors contributed 101,307 endothelial and 69,304 pericyte nuclei. The 16 Heart Cell Atlas v2 donors contributed 19,740 endothelial and 42,718 pericyte nuclei. The six Reichart donors contributed 3,871 endothelial and 6,721 pericyte nuclei. The SCP1303 vascular endothelial class includes nuclei labelled endocardial (1,882 nuclei) or lymphatic (544 nuclei) in the source subpopulation annotations. Together, these account for 2.4% of the class. These nuclei were counted as vascular endothelium, as in the donor-level comparisons, and shown under their source subpopulation names in Figure 1B. The endocardial and lymphatic endothelial rows in Figure 5 represent nuclei belonging to the source primary classes. Nine genes (*IL18*, *CASP1*, *CASP4*, *GSDMD*, *IL1B*, *CASP5*, *IL18R1*, *IL18RAP* and *IL18BP*) were excluded from the SCP1303 embedding input. Counts were normalised to 10,000 per nucleus and log-transformed. We selected 2,000 highly variable genes using donor as the batch variable and scaled their expression to a maximum of 10. Expression was reduced to 50 principal components and integrated across donors with Harmony (Scanpy 1.11.5 and harmonypy 2.1.0). For confirmation resources, we used each resource's 50-dimensional latent representation as provided. These were the scArches representation for Heart Cell Atlas v2 and principal components for Reichart. UMAP coordinates were computed with 15 neighbours, a minimum distance of 0.3 and Euclidean distance (umap-learn 0.5.12, random state 1729).

The embedding was divided into hexagonal bins. Each bin was summarised using the same transformation as the donor–class aggregates: $L = \log_2(1 + 10^6 \times \text{summed gene count within the bin} / \text{summed library count within the bin})$. Bins with at least 30 nuclei, or 15 in the smaller Reichart set, were coloured. Three grid sizes were saved for each cohort. Before transcript values were displayed, we chose the finest grid placing at least 80% of nuclei in coloured bins. The horizontal ranges contained 80 bins for SCP1303, 40 for Heart Cell Atlas v2 and 25 for Reichart. No smoothing was applied. The embedding was used for display, and inference was based on donor-level comparisons.

##### Summaries, precision and discovery support criteria

Donor draws were shared across genes within each comparison. For reproducibility, we recorded the donor order, class-pair order and random-number stream.

The endothelial–cardiomyocyte comparison of *GSDMD* used different summary methods from the other contrasts. These comprised exact median intervals, sign tests, multiplicity families and assay-specific scales. GeoMx estimates remain in upper-quartile-normalised z-score units. Sequencing and spatial effect sizes were not pooled.

#### Transcriptomic calibration

We reconstructed endothelial–pericyte contrasts for the 21,932 genes with non-duplicated names shared by both nucleus datasets. Both the full common-gene denominator and the subset satisfying precision in both cohorts were reported. The genome-wide bootstrap used 1,000 draws, specifically the first 1,000 draws from the corresponding 10,000-draw sequence. The four selected transcripts were also evaluated with 10,000 bootstrap draws.

For expression-matched reference sets, we calculated each gene's donor median of the mean of the two class-specific L values. Reference genes had to fall within ±0.5 of the corresponding target in both cohorts. Nine genes, *IL18*, *CASP1*, *CASP4*, *GSDMD*, *IL1B*, *CASP5*, *IL18R1*, *IL18RAP* and *IL18BP*, were excluded from reference selection. Direction and effect size were not used to select these reference sets. The main reference denominator additionally required precision eligibility in both cohorts. A less restricted denominator was also calculated separately.

To describe the joint frequency, we sampled distinct reference genes into four labelled slots matching *IL18*, *CASP1*, *CASP4* and *GSDMD*. Across 200,000 draws, we counted simultaneous support for the specified negative/positive/positive/positive pattern.

#### Sensitivity analyses

All additional analyses were specified and documented after the discovery data had been seen but before calculation. We evaluated disease strata, every leave-one-donor-out sample, minimum class sizes of 50 and 100 nuclei, and alternative denominators excluding mitochondrial genes, the ten highest-count non-target genes in each cohort, or both.

Library-depth sensitivity was examined by binomial thinning of aggregated counts. Each setting used 200 replicates, with 1,000 donor-bootstrap draws within each replicate. The primary setting reduced each donor's larger class library to the expected depth of the smaller class library. A more stringent setting reduced all groups to the cohort-wide minimum library. The denominator in each replicate was the realised sum of thinned target counts and the thinned aggregate of all remaining genes.

Before association testing, we selected macrophage-associated reference genes from 16 predefined candidates using abundance and macrophage-enrichment criteria. In both cohorts, eligible candidates required a macrophage median CPM within fourfold of *IL18*. This also had to be at least fourfold above the highest median CPM in endothelium, pericytes or smooth muscle. Nine candidates met these criteria. We selected the five genes with the smallest mean absolute log2 abundance distance from macrophage *IL18* across cohorts. These were *C1QB*, *SPI1*, *TYROBP*, *FCER1G* and *C1QA*. We compared their class-specific abundance and detection with *IL18*. We tested donor-level Spearman associations between pericyte *IL18* L and each of the 16 candidate L measurements or the mean L of the five selected markers. Two-sided permutation P values used 9,999 permutations. Benjamini–Hochberg correction covered the 34 tests across both cohorts. Endothelial–smooth-muscle and pericyte–smooth-muscle contrasts were also evaluated with the same four target genes.

#### Prespecified independent confirmation

The primary regional definitions were LV/AX/SP in Heart Cell Atlas and LV, including FW-labelled samples, AP and S in Reichart. Eligible nuclei from these regions were combined within donor and cell class. Heart Cell Atlas contributed CellBender-processed counts. Gene-specific counts were summed within donor–class groups and normalised using the sum of all available features.

The six prespecified cell-class positive controls showed the expected direction in all contributing donors of both resources. Ties, defined as $|\Delta L| \leq 10^{-12}$, were excluded from directional fractions but included in medians and resampling. The same saved resampling indices were used across the four genes, and percentile endpoints used linear interpolation.

#### Evaluation of the remaining survey transcripts

This evaluation was planned after the initial confirmation results had been seen. The predictions, confirmation criteria and interpretation rules were specified in writing before values in the confirmation set were calculated. Features were identified by a unique stable-ID match in each matrix and a matching gene symbol at that position. Two matrices lacked either IDs or symbols. As in the initial confirmation, features were identified at uniquely matching positions through a reference ID–symbol mapping. Before reading values for the remaining survey transcripts, we verified the same import procedure against the four genes in the initial confirmation. It reproduced all their group counts, libraries, nucleus counts and decisions. After reading the values, we left the rules, indices and code unchanged and performed the aggregation only once. Because *IL1B* is outside the eight-transcript survey, it was excluded from this evaluation. It was assessed subsequently under Evaluation of other pyroptosis-related genes (planned after confirmation). Before confirmation-set values were calculated, values for *IL1B*, *CASP5*, *IL18R1*, *IL18RAP* and *IL18BP* had been calculated unintentionally in ischaemic-heart analyses. Those analyses covered only the ischaemic-heart cohorts and excluded the confirmation resources. Predictions were determined solely from the discovery records.

#### Evaluation of other pyroptosis-related genes (planned after confirmation)

This evaluation was planned after the initial confirmation and evaluation of the remaining survey transcripts. The scope covered molecules acting within a pyroptotic cell from sensing to plasma-membrane rupture, selected by functional category. We selected inflammasome sensors (*NLRP1*, *NLRP3*, *NLRC4*, *AIM2*, *CARD8*, *MEFV*), the adaptor *PYCARD*, the substrate *IL1B*, and molecules mediating pore formation and membrane rupture (*GSDME*, *NINJ1*). Inflammatory caspases (*CASP1*, *CASP4*, *CASP5*), *IL18* and *GSDMD* had already been evaluated. We checked the scope against human annotations for GO pyroptotic inflammatory response (GO:0070269) and pyroptotic cell death (GO:0141201), and Reactome Pyroptosis (R-HSA-5620971). Genes outside the scope were recorded by category.

Genes outside the scope and the reasons for excluding them were as follows. The product of *NAIP* recognises ligands in a complex with NLRC4, so we selected *NLRC4* to represent this inflammasome. Formation of inflammasomes by the products of *IFI16*, *NLRP6*, *NLRP9* and *NLRP12* in human cells has been demonstrated only in particular infections or cell types. In many humans, *CASP12* occurs only in a non-functional form. The gasdermins encoded by *GSDMA*–*GSDMC* are expressed mainly in epithelium. *CASP3*, *CASP6*, *CASP7*, *CASP8* and *CASP10* encode caspases central to the execution or initiation of apoptosis. Transcript abundance cannot distinguish their use in apoptosis from their use in pyroptosis. *CASP3* encodes a protease that cleaves GSDME. However, GSDME determines whether pyroptosis can proceed downstream of caspase-3, so we included *GSDME* and excluded *CASP3*. *CASP8* encodes a protease that cleaves GSDMD under particular conditions, but its principal role is initiating apoptosis. *GZMA* and *GZMB* in cytotoxic lymphocytes and *ELANE* in neutrophils encode proteases produced by other cells that cleave gasdermins in target cells. Their transcripts in endothelium or pericytes do not represent the machinery of those cells themselves. *DPP9*, *MAP3K20*, *TRIM21*, *APIP*, *DHX9*, *IFI27* and *TREM2* are involved in the regulation of activity. *IRF1*, *IRF2*, *TP53* and *TP63* are involved in the regulation of transcription. ESCRT-III (*CHMP* genes) repairs GSDMD pores and suppresses pyroptosis. *HMGB1* and *IL1A* encode released molecules, whereas *BAX*, *BAK1* and *CYCS* encode molecules in mitochondrial apoptosis.

Predictions were determined solely from the discovery records, using the same rules as for the remaining survey transcripts. Directional predictions were made only for genes whose endothelial–pericyte records in both cohorts were classified as supported in the discovery genome-wide calibration table (Supplementary Data 1). The intervals used the first 1,000 bootstrap draws from the 10,000-draw sequence. For the eight survey transcripts, the classifications in this table matched those in the survey table. Under this rule, only *CARD8* had a prediction, towards endothelium. *NLRP1* favoured endothelium in all non-tied donors in both cohorts. However, its interval width in

SCP1303 (1.017) exceeded the precision criterion of 1.0, so the record was classified as undeterminable. We therefore made no prediction. *GSDME* was recorded as *GSDME* in SCP1303 and under its former name *DFNA5* in GSE183852 nuclei. It therefore lacked a shared gene name and was absent from the calibration table, so we made no prediction. We did not change predictions by recalculating with 10,000 draws or searching again under alternative names.

We used the same confirmation set, donor–class groups, confirmation criteria, saved resampling indices and seven secondary analyses as in the initial confirmation. Features were identified using the same rules as for the remaining survey transcripts. Before reading the ten-gene values, we verified the same import procedure against the four genes in the initial confirmation. It reproduced all their group tables, ΔL values and decisions. We also tested the calculation on synthetic data with ten genes before reading the ten-gene values. After reading the values, we left the rules, indices and code unchanged and performed the aggregation only once. Summaries were recalculated with separately written code and checked for agreement. For the nine genes lacking predictions, we reported values and detection in endothelium and pericytes without assigning decisions. Detection summaries comprised the numbers of groups with zero counts and median fractions of positive nuclei.

Before this evaluation, confirmation-set ΔL values for the nine genes other than *IL1B* had been calculated during the single-nucleus ATAC checks. These calculations were for a genome-wide comparison of RNA and accessibility differences. Those values were used only for quantities summarising all genes and had not been displayed for individual genes. Confirmation-set values for *IL1B* were calculated for the first time in this evaluation.

##### Comparison of deposited experimental structures (planned after the transcript analyses)

Coordinates were obtained from RCSB PDB mmCIF files for 7AL7, 3WO2, 3WO3, 3WO4, 8URV and 1J0S. Residues were mapped to UniProt numbering: Q14116 for IL-18, O95998 for IL-18BP, Q13478 for IL-18Rα and O95256 for IL-18Rβ. We used heavy atoms with occupancy greater than zero, excluding hydrogen, deuterium, water, ions, glycans and expression tags. For atoms with alternative locations, we selected the location with the highest occupancy. Ties were resolved by preferring a blank identifier, then A, then lexicographic order. Atoms of modified residues (PCA and CSO) were included. We did not add missing residues or atoms. The structures included truncated or glycan-treated constructs and extracellular receptor regions. The crystal and solution conditions also differed.

Recognition residues were defined by protein heavy-atom distances below 4.5 Å. We assessed contacts between IL-18 and IL-18BP in 7AL7, and between IL-18 and IL-18Rα or IL-18Rβ in 3WO4. We used 4.0 Å and 5.0 Å as sensitivity thresholds. This set was defined solely from 7AL7 and 3WO4 and used unchanged for the comparison of pro-IL-18 and mature IL-18.

IL-18 in 7AL7 was superposed on IL-18 in 3WO4 using 157 common Cα atoms with equal weights (Kabsch method). No outliers were excluded. The same transformation was applied to IL-18BP. We counted protein atom pairs between IL-18BP and the receptors below each distance threshold: 2 Å, 2.5 Å and 3 Å. This is a hypothetical overlay of the binding poses from two experimental structures. IL-18 residues 72–77 in 7AL7 differ in conformation from IL-18 in 3WO2, 3WO3 and 3WO4. This is because Cys74 forms a disulphide bond with IL-18BP in a symmetry-related complex in the crystal [41]. This region lies outside the IL-18BP recognition residues.

For receptor stages, we superposed the four IL-18 copies in 3WO2 and six in 3WO3 on IL-18 in 3WO4. The alignment used 156 Cα atoms common to all copies (residues 37–192). IL-18Rβ contact residues were those less than 4.5 Å from IL-18 or IL-18Rα in 3WO4.

For pro-IL-18 and mature IL-18, we analysed each of the 10 models in 8URV and 20 in 1J0S. We compared Cα distances between recognition-residue pairs whose residue numbers differed by more than three with those in bound structures. We calculated the root-mean-square difference in these distances. The comparison structure was 7AL7 for IL-18BP recognition residues and 3WO4 for IL-18Rα and IL-18Rβ recognition residues. The well-ordered ranges were specified before calculation from the precursor report based on 15N R2 [43]. These were residues 11–28, 45–60, 80–88, 97–104, 107–112, 114–120, 133–145, 147–165, 170–179 and 185–190. Model values describe an ensemble of deposited structures, not biological samples. We therefore calculated neither tests nor intervals.

Solvent-accessible surface area was calculated by the Shrake–Rupley method with a 1.4 Å probe and 960 points per atom. Atomic radii were C 1.70, N 1.55, O 1.52 and S 1.80 Å. For each 8URV model, we calculated the difference between the complete structure and a state with pro-region atoms removed. The pro region comprised residues 1–36, and the remaining coordinates were unchanged. The state with the pro region removed is not an experimental structure obtained after cleavage.

Contacts, superpositions and distance differences were recalculated from the coordinates using separately written import and calculation code. We checked that the values agreed. Solvent-accessible surface area was recalculated with another implementation using a different point arrangement. The medians of the summed areas for each recognition-residue set differed by at most 3 Å$^2$.

Figure 7B depicts heavy atoms at their deposited coordinates as spheres with a display radius of 1.9 Å. Pro-IL-18 and mature IL-18 were superposed on the IL-18 Cα atoms in 3WO4 and displayed in the same orientation. The other views have different orientations. The rendering code is included in Supplementary Software 1.

##### Checks planned after confirmation (nucleus exclusion and sex)

We planned two checks after inspecting the confirmation results. Interpretation rules were specified before calculation. Nucleus exclusion used endothelial and pericyte nuclei from the primary confirmation analysis. For each nucleus, we summed counts for five myeloid markers (*C1QA*, *C1QB*, *FCER1G*, *SPI1*, *TYROBP*). We also calculated separate sums for three endothelial markers (*PECAM1*, *CDH5*, *VWF*) and three pericyte markers (*RGS5*, *KCNJ8*, *ABCC9*). Pericyte nuclei were excluded if the myeloid sum was at least one or the endothelial sum at least two. Endothelial nuclei were excluded if the myeloid sum was at least one or the pericyte sum at least two. The myeloid and endothelial markers and the pericyte nucleus-exclusion rule matched those used in the ischaemic-heart contamination checks. We rebuilt groups from the remaining nuclei and reassessed eligibility. All 22 donors remained eligible after exclusion. We therefore applied the prespecified confirmation criteria using the same saved resampling indices as in confirmation. Sex was obtained from each resource's public metadata. For each cohort and sex, we described median ΔL and the number of donors in the predicted direction. We calculated neither intervals nor tests. Direction was not interpreted in strata with fewer than five donors.

##### Single-nucleus ATAC checks (planned after confirmation)

From the CATlas cardiac single-nucleus ATAC resource [32], we used four CARE donors [33] and two ENTEx donors. Sample-to-donor assignments followed CATlas Supplementary Table S2. Endothelium was defined by the authors' label "Endothelial Cell (Myocardial)" and pericytes by "Cardiac Pericyte 1–4". We did not relabel nuclei. For the four CARE donors, we used the matrix of 5 kb tiles from GEO (GSE165837). For the two ENTEx donors, we assigned fragments from GEO (GSE184462) to the same tiles. Nuclei were matched by exact agreement between CATlas nucleus identifiers and GEO barcodes. We checked that the genome build was hg38 using chromosome lengths for CARE and overlap with transcription start sites for ENTEx. The primary analysis used ventricles from six donors with at least 30 nuclei in both endothelium and pericytes.

For each gene, we used tiles overlapping the region within 1 kb on either side of its MANE Select transcript's transcription start site. The presence or absence of signal was scored per nucleus. Within each donor and class, this sum N was divided by T, the summed number of signal-positive tiles across nuclei. We defined $A = \log2(10^6 \times N/T + 1)$. The difference was endothelial minus pericyte accessibility. The reference set comprised 19,169 MANE Select protein_coding genes after the following exclusions. We excluded the nine genes omitted from the embedding and genes whose transcription start sites lay within 50 kb of *IL1B*, *CASP5*, *IL18R1*, *IL18RAP* or *IL18BP*. We also excluded genes sharing tiles with the four genes. The same differences were calculated for reference genes. Each gene's median was expressed as a percentile in the predicted direction within the distribution of reference-gene medians. Values were summarised descriptively.

Figure 3C tracks were generated from left-ventricular fragments from the two ENTEx donors. For endothelial and pericyte nuclei, we counted fragments in 50-base intervals within 5 kb on either side of the transcription start sites of *IL18*, *CASP1*, *CASP4* and *GSDMD*. Counts were divided by the total fragments in each class. These were descriptive tracks for display and were not used for comparisons.

Interpretation rules and predictions were fixed before the four genes' values were inspected. We changed the measurement unit twice before inspecting the values. We initially specified a fixed region around the transcription start site, then the authors' peaks. The final unit was the tile because the public CARE data were a matrix of 5 kb tiles, rather than peaks.

After inspecting the results around the *IL18* transcription start site, we planned the following three checks in sequence. For each check, interpretation and rules for reading the data were fixed before the values were viewed. First, we assessed gene-body accessibility using tiles overlapping each MANE Select transcript from 2 kb upstream of its transcription start site to the transcript end. We counted signal-positive tiles per nucleus, summed these counts and calculated A with the same formula. Nuclei labelled "Macrophage (General)" by the authors from the same six donors were positive controls. We calculated macrophage-minus-endothelial and macrophage-minus-pericyte differences around transcription start sites and across the gene body. We had specified that *IL18* accessibility would be recorded as measurable if both differences around its transcription start site were positive in all six donors. Second, we calculated genome-wide ΔL in the 22 confirmation donors in the same way as in the confirmation analysis. Before calculation, we excluded *IL1B*, *CASP5*, *IL18R1*, *IL18RAP* and *IL18BP*, whose values had not been inspected at that time. We fitted a least-squares line for 18,564 genes matched one-to-one by name to single-nucleus ATAC genes. Median ΔL was regressed on median gene-body ΔA, and residual ranks were reported. To examine macrophage RNA contamination, we selected genes before inspecting RNA values. Both median gene-body differences, macrophage-minus-endothelial and macrophage-minus-pericyte, had to rank in the top 1% of reference genes. Of these, 107 matched RNA genes. We had specified that, if their median ΔL lay within ±0.2 of the genome-wide median, we would read this as no indication of macrophage RNA preferentially entering pericyte profiles. Third, we assessed 41 tiles overlapping the region within 100 kb on either side of the *IL18* transcription start site. We classified tiles as having pericyte-specific accessibility if the endothelial-minus-pericyte difference was negative in all six donors, with a median of −0.5 or lower. The median macrophage-minus-pericyte difference also had to be below 0.5. Across the genome, we selected tiles with combined endothelial and pericyte signal in the same range as tiles around *IL18*. We calculated the expected number from the proportion of these tiles meeting the same criteria. Values are shown in Supplementary Table S10.

We added *IL18BP* using the same six donors and measurement method after confirming its endothelial RNA bias in the confirmation set. This addition also followed inspection of the single-nucleus ATAC results for the four genes. The endothelial prediction was fixed before the *IL18BP* values were counted. The criterion for an endothelial direction, specified before the values were counted, required at least five of six donors on the endothelial side and a positive median difference. This donor-count criterion did not apply to the four-gene checks. We used the same T values as for the four genes and the reference genes. These values had been calculated with *IL18BP* tiles excluded. Tiles around the *IL18BP* transcription start site also overlap the 3′ end of the neighbouring *RNF121* transcript. The second gene-body tile also overlaps the 3′ end of the *NUMA1* transcript.

#### Ischaemic-heart data and groups

GSE270788 [19] included five donors with acute myocardial infarction (AMI), five with ischaemic cardiomyopathy (ICM) and seven controls. Of these, one ICM donor and one control donor had fewer than 30 pericyte nuclei. Six donors with non-ischaemic cardiomyopathy from the same cohort were excluded from the ischaemic-heart comparisons. From Kuppe [20], we used seven AMI, eight ICM and four control donors, excluding one donor whose clinical group was coronary artery disease and heart failure. Clinical group, region and days after infarction followed Supplementary Table 1 of the original study and the published specimen metadata. Specimens were collected 2–11 days after infarction in AMI and 31–166 days after infarction in ICM. SCP1849 [21] included seven ICM and eight non-failing control donors. Controls were the groups annotated as Donor in GSE270788, control in Kuppe and NF (non-failing) in SCP1849. Donors were defined by public sample IDs (MA numbers) in GSE270788, patient IDs in Kuppe and donor IDs in SCP1849. In Kuppe, counts from all specimens from each donor were summed, except in region-specific analyses. For GSE270788, we used gene-expression features from the public filtered matrix and 46,384 nuclei with public annotations. We used raw counts from the processed data in Kuppe and Cell Ranger raw counts in SCP1849. Cell classes followed the published annotations. The endothelial and pericyte classes were Endothelium and Pericyte in GSE270788, and Endothelial and Pericyte in Kuppe. Kuppe's Endothelial class included lymphatic endothelium and endocardium. In SCP1849, endothelium comprised the EC-General capillary, EC-KCNIP4, EC-Angiogenic and EC-Arterial subpopulations. Nuclei annotated with the cell type pericyte (CL:0000669) constituted the pericyte class. Class-level comparisons were made between AMI and controls in GSE270788 and Kuppe. Nine classes were matched across both cohorts: endothelium, pericytes, myeloid cells, fibroblasts, cardiomyocytes, T/NK cells (lymphocytes in Kuppe), smooth muscle, mast cells and adipocytes. For row labels in Figure 6B and Supplementary Figure S15A, Endothelium denotes Endothelium in GSE270788 and Endothelial in Kuppe. The Smooth muscle label denotes SMC in GSE270788 and vSMCs in Kuppe. The "T/NK, lymphoid" label denotes TNKCells in GSE270788 and Lymphoid in Kuppe.

#### Measures and summaries in ischaemic hearts

The definitions of L, ΔL, ties and donor–class eligibility were the same as in the main Methods. Each group was summarised by the median ΔL, the number of donors in the predicted direction and the number of ties. The prespecified criterion was that at least 75% of non-tied donors had the predicted direction. For all quantities, percentile 95% bootstrap intervals were obtained by resampling donors with replacement 10,000 times within groups, or units within strata for region-specific analyses. Count-pooled L was calculated after separately summing the counts and library totals of eligible donors for each group and class. Count-pooled ΔL was calculated using donors eligible for both endothelium and pericytes, and its difference between AMI and controls was calculated. The change in each class was the AMI-minus-control difference in count-pooled L among donors eligible for that class. Intervals for this change were calculated only when both groups contained at least two donors. Changes in *IL18*, *CASP1* and *GSDMD* for each class were calculated in the same way. For each class-level change, two-sided P values were obtained by enumerating all donor-label assignments, and Benjamini–Hochberg q values used all classes within each cohort and gene as the family, comprising 14 classes in GSE270788 and 11 in Kuppe. Classes for which P could not be calculated were included in the family with P = 1. The fraction of *IL18* counts was calculated using all nuclei from all donors in each group. Each class's counts were divided by the total across all classes. For gene-to-gene comparisons and selection of specific genes, candidates comprised all features in each cohort's count matrix. The same nine genes as in transcriptomic calibration were excluded. Expression-matched reference genes had counts in pericytes in both AMI and controls. Their count-pooled pericyte L across the two groups had to lie within ±0.5 of *CASP4*. Among these genes, we reported the fraction with pericyte changes at least as large as that of *CASP4*. We also reported the fraction whose difference in count-pooled ΔL was at most that of *CASP4*.

#### Region-specific analyses in ischaemic hearts

In Kuppe, the unit was the patient–region combination, with specimens within the same unit pooled. Units were divided into five strata according to the recorded specimen region: control, AMI remote myocardium, border zone, infarct zone and ICM. Remote myocardium comprised specimens annotated as RZ, border zone those annotated as BZ, and infarct zone those annotated as IZ. Each ICM patient contributed one specimen, which was assigned to the ICM stratum regardless of region. Units eligible for both endothelium and pericytes were used to calculate count-pooled L within each stratum and its difference from controls. Intervals were calculated only for strata with at least three units. The contrast ratio κ was defined as the stratum's count-pooled ΔL divided by the control count-pooled ΔL. Endothelial-biased reference genes were defined in controls. Endothelial counts were at least 20, and count-pooled ΔL was at least 1. The mean of endothelial and pericyte count-pooled L lay within ±0.5 of *CASP4*. We calculated the median ratio of these genes and the fraction with ratios at most that of *CASP4*. Endothelial and pericyte changes in each stratum were compared with genes with counts in that class in both the stratum and controls. Their count-pooled L across the stratum and controls had to lie within ±0.5 of *CASP4*. We reported the fraction with changes at most that of *CASP4* in endothelium and at least that of *CASP4* in pericytes. For each unit, ΔL was divided by the control count-pooled ΔL. This ratio was compared with the median ratio of the endothelial-biased reference genes in the same unit.

##### Checks for contamination in pericyte *CASP4*

The first check used a contamination model based on five myeloid markers. These were the macrophage markers selected for the sensitivity analyses. *CASP4* counts attributable to contamination were estimated for each specimen. Each GSE270788 donor contributed one specimen. We multiplied the myeloid-marker counts in pericyte nuclei by the *CASP4*-to-marker count ratio in the contamination source. The contamination source was either all nuclei with public annotations from the specimen, including pericytes, or its myeloid nuclei. Observed and estimated CPM were calculated for each group by pooling donors with eligible pericytes. We calculated the AMI-minus-control difference for the observed values, the estimates and the remaining values after subtraction. The second check subtracted contributions from other classes. Four classes were used as contamination sources: myeloid cells, endothelium, fibroblasts and cardiomyocytes. GSE270788 endothelium included endocardium and lymphatic endothelium. Genes specific to each class were identified by pooling nuclei from all control donors. Each gene required a CPM of at least 50 in that class. It also required at least four times the CPM of every other class with at least 100 control nuclei. Contamination sources with at least five specific genes were used for estimation. Estimates used the first check's formula, replacing the markers with the summed counts of specific genes. The estimates were summed across contamination sources. When a specimen had fewer than 30 nuclei in the source class or zero specific-gene counts, a donor-level ratio was used. That ratio pooled all specimens from the donor. The third check excluded pericyte nuclei with total counts of at least 1 across the five myeloid markers or at least 2 across the endothelial markers *PECAM1*, *CDH5* and *VWF*. The fraction of excluded nuclei was calculated for each group. Donor–class groups and eligibility were reassessed using the remaining nuclei. The pericyte-class change was then calculated. The pericyte-class change before nucleus exclusion was also estimated after leaving out one AMI or control donor at a time. In GSE270788, we also calculated point estimates after excluding samples assigned to processing batch 3 in the original study's public code. This batch included additional sequencing. These estimates used the remaining three controls (MA5, MA6, MA13) and two AMI donors (MA9, MA11). Interpretation rules were specified before calculation. For the contamination model, the remaining difference had to have an interval entirely above zero in all four combinations. These combinations comprised two cohorts and two contamination sources. If this condition held, higher pericyte *CASP4* in AMI could not be explained by contamination alone. Otherwise, explanation by contamination alone remained possible. The conditions for the other checks were assessed within each cohort. Subtraction of contributions from other classes required an interval above zero for the remaining difference. This condition was not evaluated in a cohort if any contamination source could not be estimated. Nucleus exclusion required an interval above zero after exclusion. Leave-one-donor-out estimates had to be positive in every case. If every evaluable condition held, higher pericyte *CASP4* in AMI was classified as persisting. If none held, the conditions were classified as unmet. Otherwise, the interpretation was that the checks gave mixed results.

##### Single-nucleus ATAC in ischaemic hearts (planned after the RNA results)

This analysis was planned after inspecting the ischaemic-heart RNA results. Calculation methods were specified before the ATAC values were read. We used 17 libraries of Kuppe [20] single-nucleus ATAC fragments (cellranger-atac 2.0.0, GRCh38). They represented four controls and patient–region units from five AMI patients. The AMI units comprised four in remote myocardium, two in the border zone and three in the infarct zone. Nuclei were matched by exact agreement between identifiers in the processed data (library and barcode) and fragment barcodes. Endothelium and pericytes followed the authors' labels, Endothelial and Pericyte. These labels were not reassigned. Regions followed the single-nucleus RNA specimen regions for the same patient–region combinations. Units required at least 30 nuclei in both endothelium and pericytes. Three other infarct-zone units with few nuclei were not used.

Accessibility A used the same 5 kb tiles and formula as in Single-nucleus ATAC checks (planned after confirmation). Fragments were counted in every tile they overlapped by at least one base. N and T were summed within each unit and class. For each stratum, eligible units were pooled. For overall AMI versus controls, all eligible units were first pooled within patients, then within groups. The change in each class was A in the stratum or overall AMI minus A in controls. For the endothelial–pericyte difference, ΔA = A(endothelium) − A(pericyte), we also calculated the difference between each stratum and controls. Percentile 95% bootstrap intervals used 10,000 resamples with replacement of units within each group. For overall AMI versus controls, patients were resampled instead. Intervals were calculated only when both groups contained at least three units. *CASP4* changes were expressed as percentiles in the predicted direction within distributions of the same quantities for 19,169 reference genes. The predicted directions were positive for pericyte changes and negative for changes in ΔA. We also examined reference genes whose pericyte A for controls and AMI combined lay within ±0.5 of *CASP4*. We reported the fraction with changes equal to that of *CASP4* or larger in the predicted direction. No tests were performed, and values were summarised descriptively.

##### Reproducibility and ethics

Independent implementations reconstructed the eight central cohort–gene summaries from integer pseudobulk arrays and compared them with the original results. Counts, medians, signs, ties and bootstrap endpoints were checked. Additional scalar or direct-resampling implementations checked the calibration and simulation calculations.

#### Measurement conventions

EC denotes vascular endothelium and PC pericytes. The principal difference is ΔL = L(EC) − L(PC), with L = log2(1 + CPM). Negative differences favour pericytes and positive differences favour endothelium. Counts are summed within donor and cell class, using all available features for the library denominator. Eligible donor–class groups contain at least 30 nuclei and have a positive library total. Ties are included in medians and bootstrap resampling and excluded from directional fractions.

#### Prior-knowledge disclosure

Before the confirmation directions and rules were fixed, the investigators had inspected all-gene pseudobulk summaries of both discovery cohorts (SCP1303 and GSE183852 nuclei) and the endothelial–cardiomyocyte *GSDMD* analyses. They had inspected the class-marker and differential-expression supplementary tables of Litviňuková et al. (Supplementary Tables 1–35). In these tables, *IL18* appears among markers of mesothelial cells and LYVE1-positive macrophages. *CASP4* appears among markers of a venous endothelial state (EC6_ven), and *GSDMD* among markers of an immune-associated endothelial state (EC4_immune). *CASP1* appears among markers of CD16-positive monocytes, *IL18R1* among mast-cell markers and *IL18RAP* among NK and NKT-cell markers. They had also inspected Supplementary Tables 1–9 of the HeartMap integrated atlas, in which *IL18* appears among epicardial-lineage markers and *IL18R1* among mast-cell markers. Other information inspected comprised the supplementary tables of the discovery cohort publications, the main text and figures of Reichart et al., and Human Protein Atlas summaries for GSDMD and CASP4. They had inspected metadata-level information on the candidate confirmation resources, including donor numbers, annotations and library depth, and the immunohistochemical localisation of IL-18 protein to endothelial cells and macrophages in failing ischaemic myocardium reported by Mallat et al. No target-gene values, differential-expression tables or gene pages from the confirmation resources were opened before the rules were fixed, and the Reichart supplementary tables S1–S71 were not inspected. The prediction document dated 10 October 2026 and the confirmation specification, with their SHA-256 values and English summaries, are provided in Supplementary Software 1.

#### Limits of the spatial marker-score analysis

Marker-score sensitivity does not establish causal mediation, cellular purity or an endothelial–pericyte spatial interaction.

#### Accompanying electronic files

Supplementary Data 1 provides numerical tables and metadata in UTF-8 format. Supplementary Software 1 contains analysis and plotting code, analysis specifications, saved resampling indices and source-data records. Large original matrices are not redistributed. Re-execution requires the source resources and the recorded environment. A codebook in Supplementary Data 1 gives English equivalents for Japanese annotation strings in the numerical tables.

The embedding coordinates, bin tables at all three grid sizes, cell-state positions and grid-selection records for Figure 1 and Supplementary Figures S8–S10 are in Supplementary Data 1. The code that produced them and the plotting code are in Supplementary Software 1.

#### Supplementary Table S1. Discovery endothelial–pericyte contrasts

| Gene | Resource | N | Median ΔL | 95% interval | Predicted / non-tie | Ties |
|---|---|---|---|---|---|---|
| *IL18* | SCP1303 | 42 | −2.121 | −2.438 to −1.763 | 39/41 | 1 |
| *IL18* | GSE183852 nuclei | 38 | −1.317 | −1.751 to −0.850 | 33/34 | 4 |
| *CASP1* | SCP1303 | 42 | 2.677 | 2.396–2.830 | 42/42 | 0 |
| *CASP1* | GSE183852 nuclei | 38 | 3.097 | 2.879–3.436 | 38/38 | 0 |
| *CASP4* | SCP1303 | 42 | 2.016 | 1.796–2.325 | 42/42 | 0 |
| *CASP4* | GSE183852 nuclei | 38 | 2.358 | 2.071–2.438 | 38/38 | 0 |
| *GSDMD* | SCP1303 | 42 | 2.325 | 2.212–2.694 | 41/42 | 0 |
| *GSDMD* | GSE183852 nuclei | 38 | 1.688 | 1.496–1.873 | 38/38 | 0 |

Predicted direction is negative for *IL18* and positive for *CASP1*, *CASP4* and *GSDMD*. Directional fractions exclude ties, whereas total N includes them. The four directions were selected after the full discovery survey. Exact source values and all other class pairs are provided in Supplementary Data 1.

#### Supplementary Table S2. Primary confirmation estimates

| Gene | Resource | N | Median ΔL | 95% interval | Predicted / non-tie | Ties |
|---|---|---|---|---|---|---|
| *IL18* | Heart Cell Atlas v2 | 16 | −1.565 | −2.094 to −1.179 | 16/16 | 0 |
| *IL18* | Reichart | 6 | −1.419 | −2.161 to −0.651 | 6/6 | 0 |
| *IL18* | Pooled | 22 | −1.451 | −1.977 to −1.255 | 22/22 | 0 |
| *CASP1* | Heart Cell Atlas v2 | 16 | 2.957 | 2.474–3.810 | 16/16 | 0 |
| *CASP1* | Reichart | 6 | 3.174 | 2.304–3.482 | 6/6 | 0 |
| *CASP1* | Pooled | 22 | 3.020 | 2.773–3.561 | 22/22 | 0 |
| *CASP4* | Heart Cell Atlas v2 | 16 | 2.416 | 2.168–3.026 | 16/16 | 0 |
| *CASP4* | Reichart | 6 | 2.194 | 1.631–2.680 | 6/6 | 0 |
| *CASP4* | Pooled | 22 | 2.395 | 2.160–2.749 | 22/22 | 0 |
| *GSDMD* | Heart Cell Atlas v2 | 16 | 1.762 | 1.525–2.045 | 16/16 | 0 |
| *GSDMD* | Reichart | 6 | 1.792 | 1.456–2.116 | 6/6 | 0 |
| *GSDMD* | Pooled | 22 | 1.762 | 1.558–1.937 | 22/22 | 0 |

Each gene met the prespecified pooled criteria, and neither contributing dataset had an opposing median. The formal decision concerns the pooled donor set. All four marginal intervals are unadjusted for multiplicity. Source: Supplementary Data 1.

#### Supplementary Table S3. Prespecified secondary analyses

| Analysis | Gene | N | Median | 95% interval | Predicted / non-tie | Ties |
|---|---|---|---|---|---|---|
| Capillary endothelium and pericytes | *IL18* | 22 | −1.478 | −1.807 to −1.063 | 22/22 | 0 |
| Capillary endothelium and pericytes | *CASP1* | 22 | 3.068 | 2.463–3.631 | 22/22 | 0 |
| Capillary endothelium and pericytes | *CASP4* | 22 | 2.217 | 2.012–2.624 | 22/22 | 0 |
| Capillary endothelium and pericytes | *GSDMD* | 22 | 1.722 | 1.584–2.027 | 22/22 | 0 |
| Exclusion of predefined '-like' states | *IL18* | 22 | −1.463 | −1.967 to −1.266 | 22/22 | 0 |
| Exclusion of predefined '-like' states | *CASP1* | 22 | 3.003 | 2.885–3.531 | 22/22 | 0 |
| Exclusion of predefined '-like' states | *CASP4* | 22 | 2.360 | 2.137–2.748 | 22/22 | 0 |
| Exclusion of predefined '-like' states | *GSDMD* | 22 | 1.734 | 1.570–1.912 | 22/22 | 0 |
| Right ventricular | *IL18* | 19 | −1.884 | −2.291 to −1.448 | 18/18 | 1 |
| Right ventricular | *CASP1* | 19 | 3.308 | 1.975–4.436 | 15/15 | 4 |
| Right ventricular | *CASP4* | 19 | 2.207 | 1.890–2.784 | 19/19 | 0 |
| Right ventricular | *GSDMD* | 19 | 1.543 | 1.223–2.222 | 19/19 | 0 |
| Atrial | *IL18* | 15 | −1.191 | −1.597 to −1.060 | 12/14 | 1 |
| Atrial | *CASP1* | 15 | 2.550 | 1.298–4.643 | 14/14 | 1 |
| Atrial | *CASP4* | 15 | 1.859 | 1.233–2.284 | 15/15 | 0 |
| Atrial | *GSDMD* | 15 | 1.685 | 1.443–2.493 | 15/15 | 0 |
| Reichart cardiomyopathy ventricles | *IL18* | 52 | −1.693 | −1.802 to −1.466 | 51/52 | 0 |
| Reichart cardiomyopathy ventricles | *CASP1* | 52 | 2.347 | 2.099–2.683 | 52/52 | 0 |
| Reichart cardiomyopathy ventricles | *CASP4* | 52 | 1.893 | 1.766–2.052 | 52/52 | 0 |
| Reichart cardiomyopathy ventricles | *GSDMD* | 52 | 1.574 | 1.435–1.716 | 52/52 | 0 |

| Analysis | Gene | N | Median | 95% interval | Predicted / non-tie | Ties |
|---|---|---|---|---|---|---|
| Differences in fractions of positive nuclei | *IL18* | 22 | −0.028 | −0.043 to −0.019 | 22/22 | 0 |
| Differences in fractions of positive nuclei | *CASP1* | 22 | 0.018 | 0.010–0.021 | 22/22 | 0 |
| Differences in fractions of positive nuclei | *CASP4* | 22 | 0.068 | 0.059–0.083 | 22/22 | 0 |
| Differences in fractions of positive nuclei | *GSDMD* | 22 | 0.051 | 0.042–0.058 | 22/22 | 0 |

The predefined '-like' annotations were EC9_FB-like, EC10_CMC-like and PC4_CMC-like. Dataset-specific primary summaries are provided in Supplementary Table S2 and Supplementary Figure S7. The rows for differences in fractions of positive nuclei use Δp, to which the ΔL magnitude threshold does not apply. A discrepancy in right ventricular sex metadata was documented. Donors with reverse or tied differences are included in the summaries. All secondary analyses are descriptive.

#### Supplementary Table S4. Endothelial–cardiomyocyte comparison of *GSDMD*

| Resource | N | Positive | Median | Exact 95% CI | Scale |
|---|---|---|---|---|---|
| SCP1303 | 42 | 42/42 | 3.738 | 3.430–3.995 | log2(CPM + 1) |
| GSE183852 nuclei | 38 | 38/38 | 3.272 | 3.089–3.540 | log2(CPM + 1) |
| GSE271676 | 32 | 27/32 | 1.591 | 0.666–2.044 | Upper-quartile-normalised z-score units |

These contrasts use cardiomyocytes as the comparator. GeoMx values are shown on their original upper-quartile-normalised z-score scale, separately from log2(CPM + 1) values. The five negative GeoMx contrasts are shown in Supplementary Figure S12. The within-cohort signed-contrast percentiles are 94.9, 95.1 and 97.4 in the same resource order. These are descriptive transcriptome positions, not P values.

#### GeoMx measured-composition diagnostic

| Coefficient | Estimate | HC3 95% interval |
|---|---|---|
| Intercept | 0.467 | −0.896 to 1.830 |
| Myeloid | 0.492 | 0.097–0.887 |
| Stromal | 0.307 | −0.452 to 1.066 |

All 32 complete spatial donors contribute. These are OLS coefficients with HC3 normal-Wald intervals. Source: Supplementary Data 1.

#### Supplementary Table S5. Check of the primary confirmation analysis after exclusion of nuclei with suspected contamination

(a) Excluded nuclei

| Dataset | Class | Nuclei before exclusion | Excluded nuclei | With myeloid markers | With opposite-class markers | With both | Nuclei after exclusion | Fraction excluded | Fraction per donor (minimum/median/maximum) |
|---|---|---|---|---|---|---|---|---|---|
| A (Heart Cell Atlas v2) | Endothelium | 19,740 | 1,346 | 189 | 1,184 | 27 | 18,394 | 6.82% | 0.77%/5.36%/14.01% |
| A (Heart Cell Atlas v2) | Pericytes | 42,718 | 218 | 123 | 96 | 1 | 42,500 | 0.51% | 0.12%/0.30%/4.18% |
| B (Reichart) | Endothelium | 3,871 | 216 | 133 | 86 | 3 | 3,655 | 5.58% | 2.73%/5.32%/11.11% |
| B (Reichart) | Pericytes | 6,721 | 180 | 163 | 18 | 1 | 6,541 | 2.68% | 0.93%/3.18%/4.44% |

The myeloid-marker and opposite-class-marker columns both include nuclei carrying both sets of markers (excluded nuclei = myeloid + opposite class − both). Opposite-class markers are endothelial markers for pericyte nuclei and pericyte markers for endothelial nuclei.

(b) Summaries before and after exclusion

| Gene | Pooled median before exclusion (95% interval) | Pooled median after exclusion (95% interval) | Donors in the predicted direction after exclusion | Medians in A and B after exclusion | Decision after exclusion |
|---|---|---|---|---|---|
| *IL18* | −1.451 (−1.977 to −1.255) | −1.588 (−2.108 to −1.239) | 22/22 | −1.630/−1.283 | Met confirmation criteria |
| *CASP1* | 3.020 (2.773–3.561) | 3.035 (2.701–3.596) | 22/22 | 2.981/3.228 | Met confirmation criteria |
| *CASP4* | 2.395 (2.160–2.749) | 2.467 (2.208–2.749) | 22/22 | 2.492/2.208 | Met confirmation criteria |
| *GSDMD* | 1.762 (1.558–1.937) | 1.755 (1.609–1.949) | 22/22 | 1.755/1.843 | Met confirmation criteria |

The difference is $\Delta L = L(EC) - L(PC)$. Percentile intervals used 10,000 donor resamples stratified by dataset, with the same saved resampling indices as in confirmation. Decisions follow the prespecified confirmation criteria.

#### Supplementary Table S6. Endothelial–pericyte differences (ΔL) stratified by sex

| Cohort | Sex | N | *IL18* | *CASP1* | *CASP4* | *GSDMD* |
|---|---|---|---|---|---|---|
| SCP1303 | Female | 19 | −2.128 (19/19) | 2.449 (19/19) | 1.774 (19/19) | 2.193 (19/19) |
| SCP1303 | Male | 23 | −2.113 (20/22, 1 tie) | 2.806 (23/23) | 2.317 (23/23) | 2.553 (22/23) |
| GSE183852 nuclei | Female | 19 | −0.871 (15/16, 3 ties) | 3.049 (19/19) | 2.340 (19/19) | 1.726 (19/19) |
| GSE183852 nuclei | Male | 19 | −1.708 (18/18, 1 tie) | 3.218 (19/19) | 2.376 (19/19) | 1.561 (19/19) |
| Confirmation (22 donors pooled) | Female | 9 | −1.935 (9/9) | 2.686 (9/9) | 2.393 (9/9) | 1.806 (9/9) |
| Confirmation (22 donors pooled) | Male | 13 | −1.442 (13/13) | 3.426 (13/13) | 2.436 (13/13) | 1.718 (13/13) |
| Confirmation A (Heart Cell Atlas v2) | Female | 8 | −1.956 (8/8) | 2.387 (8/8) | 2.383 (8/8) | 1.832 (8/8) |
| Confirmation A (Heart Cell Atlas v2) | Male | 8 | −1.433 (8/8) | 3.667 (8/8) | 2.729 (8/8) | 1.643 (8/8) |
| Confirmation B (Reichart) | Female | 1 (direction not interpreted because of the small sample) | −0.473 (1/1) | 3.048 (1/1) | 2.816 (1/1) | 1.549 (1/1) |
| Confirmation B (Reichart) | Male | 5 | −1.460 (5/5) | 3.300 (5/5) | 2.126 (5/5) | 1.934 (5/5) |

Each entry gives the donor median of ΔL = L(EC) − L(PC) (number of donors in the predicted direction/number of non-tied donors). Predicted direction is negative for *IL18* and positive for *CASP1*, *CASP4* and *GSDMD*. Neither intervals nor tests were calculated. Directions in strata with fewer than five donors were left uninterpreted. Values are from before nucleus exclusion, with all donors from both discovery cohorts and the 22 primary confirmation donors.

#### Supplementary Table S7. Endothelial-minus-pericyte differences for the remaining survey transcripts in the confirmation set and secondary analyses

The difference is ΔL = L(EC) − L(PC). Rows for fractions of positive nuclei show Δp in percentage points. Percentile intervals used 10,000 donor resamples stratified by dataset, with the same saved resampling indices as in the initial confirmation. Only the primary *IL18BP* analysis had a prediction and was evaluated against the prespecified confirmation criteria. The other rows are descriptive.

| Transcript | Analysis | N | Endothelial side | Pericyte side | Ties | Median (95% interval) |
|---|---|---|---|---|---|---|
| *IL18BP* | Primary analysis (22 donors) | 22 | 22 | 0 | 0 | 1.575 (1.377–1.867) |
| *IL18BP* | Capillary endothelium and pericytes | 22 | 21 | 1 | 0 | 1.236 (0.978–1.753) |
| *IL18BP* | Exclusion of predefined '-like' states | 22 | 22 | 0 | 0 | 1.502 (1.248–1.862) |
| *IL18BP* | Right ventricular | 19 | 12 | 6 | 1 | 1.477 (−0.240 to 2.706) |
| *IL18BP* | Atrial | 15 | 13 | 1 | 1 | 0.856 (0.221–1.318) |
| *IL18BP* | Reichart cardiomyopathy | 52 | 43 | 9 | 0 | 0.801 (0.676–0.994) |
| *IL18BP* | Difference in positive-nucleus fraction (percentage points) | 22 | 21 | 1 | 0 | 0.688 (0.386–0.895) |
| *IL18BP* | Heart Cell Atlas v2 | 16 | 16 | 0 | 0 | 1.584 (1.377–2.453) |
| *IL18BP* | Reichart | 6 | 6 | 0 | 0 | 1.409 (0.530–1.820) |
| *IL18R1* | Primary analysis (22 donors) | 22 | 17 | 5 | 0 | 0.763 (0.428–1.264) |
| *IL18R1* | Capillary endothelium and pericytes | 22 | 18 | 4 | 0 | 0.834 (0.531–1.254) |
| *IL18R1* | Exclusion of predefined '-like' states | 22 | 17 | 5 | 0 | 0.792 (0.425–1.188) |
| *IL18R1* | Right ventricular | 19 | 16 | 3 | 0 | 0.598 (0.237–1.673) |
| *IL18R1* | Atrial | 15 | 13 | 2 | 0 | 1.704 (1.366–2.288) |
| *IL18R1* | Reichart cardiomyopathy | 52 | 49 | 3 | 0 | 1.200 (0.883–1.438) |
| *IL18R1* | Difference in positive-nucleus fraction (percentage points) | 22 | 17 | 5 | 0 | 1.444 (0.505–6.290) |
| *IL18R1* | Heart Cell Atlas v2 | 16 | 11 | 5 | 0 | 0.763 (−0.119 to 1.264) |
| *IL18R1* | Reichart | 6 | 6 | 0 | 0 | 0.824 (0.315–1.898) |

| Transcript | Analysis | N | Endothelial side | Pericyte side | Ties | Median (95% interval) |
|---|---|---|---|---|---|---|
| *IL18RAP* | Primary analysis (22 donors) | 22 | 17 | 4 | 1 | 0.961 (0.326–1.248) |
| *IL18RAP* | Capillary endothelium and pericytes | 22 | 12 | 4 | 6 | 0.563 (0.000–1.350) |
| *IL18RAP* | Exclusion of predefined '-like' states | 22 | 17 | 4 | 1 | 0.964 (0.336–1.294) |
| *IL18RAP* | Right ventricular | 19 | 2 | 6 | 11 | 0.000 (−0.159 to 0.000) |
| *IL18RAP* | Atrial | 15 | 8 | 4 | 3 | 0.874 (−0.491 to 1.762) |
| *IL18RAP* | Reichart cardiomyopathy | 52 | 35 | 11 | 6 | 0.569 (0.043–0.753) |
| *IL18RAP* | Difference in positive-nucleus fraction (percentage points) | 22 | 17 | 4 | 1 | 0.084 (0.049–0.152) |
| *IL18RAP* | Heart Cell Atlas v2 | 16 | 13 | 3 | 0 | 0.851 (0.326–1.248) |
| *IL18RAP* | Reichart | 6 | 4 | 1 | 1 | 1.048 (−0.348 to 1.795) |
| *CASP5* | Primary analysis (22 donors) | 22 | 4 | 7 | 11 | 0.000 (−0.100 to 0.000) |
| *CASP5* | Capillary endothelium and pericytes | 22 | 4 | 6 | 12 | 0.000 (0.000–0.000) |
| *CASP5* | Exclusion of predefined '-like' states | 22 | 4 | 7 | 11 | 0.000 (−0.102 to 0.000) |
| *CASP5* | Right ventricular | 19 | 3 | 1 | 15 | 0.000 (0.000–0.000) |
| *CASP5* | Atrial | 15 | 4 | 3 | 8 | 0.000 (0.000–0.030) |
| *CASP5* | Reichart cardiomyopathy | 52 | 31 | 5 | 16 | 0.443 (0.000–0.702) |
| *CASP5* | Difference in positive-nucleus fraction (percentage points) | 22 | 4 | 7 | 11 | 0.000 (−0.010 to 0.000) |
| *CASP5* | Heart Cell Atlas v2 | 16 | 2 | 7 | 7 | 0.000 (−0.268 to 0.000) |
| *CASP5* | Reichart | 6 | 2 | 0 | 4 | 0.000 (0.000–1.273) |

**Supplementary Table S8. Accessibility around the transcription start sites of four genes in single-nucleus ATAC data from ischaemic hearts (Kuppe)**

(A) Accessibility by class and differences from controls

| Gene | Group/ stratum | Units | Endothelial N | Endothelial T | Endothelial A | Endothelial difference (95% interval) | Pericyte N | Pericyte T | Pericyte A | Pericyte difference (95% interval) | ΔA | Difference in ΔA (95% interval) |
|---|---|---|---|---|---|---|---|---|---|---|---|---|
| *IL18* | Controls | 4 | 26 | 13,846,048 | 1.525 | — | 16 | 9,385,088 | 1.436 | — | 0.089 | — |
| *IL18* | Overall AMI | 5 | 84 | 37,076,657 | 1.707 | 0.182 (−0.126 to 0.505) | 27 | 19,380,734 | 1.259 | −0.177 (−0.520 to 0.125) | 0.448 | 0.359 (−0.212 to 0.983) |
| *IL18* | Remote myocardium | 4 | 45 | 17,826,516 | 1.817 | 0.292 (0.049–0.568) | 12 | 10,681,107 | 1.086 | −0.349 (−0.962 to 0.125) | 0.731 | 0.641 (0.092–1.256) |
| *IL18* | Border zone | 2 | 9 | 7,568,672 | 1.130 | −0.395 (—) | 8 | 4,949,362 | 1.388 | −0.048 (—) | −0.257 | −0.347 (—) |
| *IL18* | Infarct zone | 3 | 30 | 11,681,469 | 1.835 | 0.310 (−0.765 to 0.664) | 7 | 3,750,265 | 1.519 | 0.084 (−1.398 to 0.618) | 0.316 | 0.226 (−1.415 to 2.020) |
| *CASP1* | Controls | 4 | 130 | 13,846,048 | 3.377 | — | 38 | 9,385,088 | 2.336 | — | 1.041 | — |
| *CASP1* | Overall AMI | 5 | 253 | 37,076,657 | 2.968 | −0.409 (−0.615 to −0.164) | 69 | 19,380,734 | 2.189 | −0.147 (−0.733 to 0.560) | 0.779 | −0.262 (−1.157 to 0.466) |
| *CASP1* | Remote myocardium | 4 | 116 | 17,826,516 | 2.908 | −0.469 (−0.741 to −0.194) | 34 | 10,681,107 | 2.065 | −0.271 (−0.927 to 0.455) | 0.844 | −0.197 (−1.080 to 0.588) |
| *CASP1* | Border zone | 2 | 57 | 7,568,672 | 3.093 | −0.284 (—) | 23 | 4,949,362 | 2.498 | 0.162 (—) | 0.595 | −0.446 (—) |
| *CASP1* | Infarct zone | 3 | 80 | 11,681,469 | 2.972 | −0.405 (−0.709 to −0.146) | 12 | 3,750,265 | 2.070 | −0.266 (−2.159 to 0.993) | 0.902 | −0.139 (−1.599 to 1.548) |
| *CASP4* | Controls | 4 | 152 | 13,846,048 | 3.582 | — | 46 | 9,385,088 | 2.561 | — | 1.021 | — |
| *CASP4* | Overall AMI | 5 | 365 | 37,076,657 | 3.439 | −0.143 (−0.288 to 0.080) | 100 | 19,380,734 | 2.623 | 0.062 (−0.255 to 0.462) | 0.816 | −0.205 (−0.640 to 0.227) |
| *CASP4* | Remote myocardium | 4 | 161 | 17,826,516 | 3.326 | −0.256 (−0.476 to 0.091) | 50 | 10,681,107 | 2.506 | −0.055 (−0.407 to 0.459) | 0.820 | −0.201 (−0.770 to 0.270) |
| *CASP4* | Border zone | 2 | 98 | 7,568,672 | 3.802 | 0.220 (—) | 27 | 4,949,362 | 2.690 | 0.129 (—) | 1.112 | 0.090 (—) |
| *CASP4* | Infarct zone | 3 | 106 | 11,681,469 | 3.333 | −0.250 (−0.680 to −0.103) | 23 | 3,750,265 | 2.834 | 0.273 (−0.106 to 0.920) | 0.498 | −0.523 (−1.569 to −0.002) |
| *GSDMD* | Controls | 4 | 264 | 13,846,048 | 4.327 | — | 101 | 9,385,088 | 3.556 | — | 0.771 | — |
| *GSDMD* | Overall AMI | 5 | 585 | 37,076,657 | 4.069 | −0.258 (−0.671 to 0.193) | 173 | 19,380,734 | 3.311 | −0.245 (−0.582 to 0.188) | 0.757 | −0.013 (−0.327 to 0.394) |

| Gene | Group/ stratum | Units | Endothelial N | Endothelial T | Endothelial A | Endothelial difference (95% interval) | Pericyte N | Pericyte T | Pericyte A | Pericyte difference (95% interval) | ΔA | Difference in ΔA (95% interval) |
|---|---|---|---|---|---|---|---|---|---|---|---|---|
| *GSDMD* | Remote myocardium | 4 | 322 | 17,826,516 | 4.253 | −0.074 (−0.491 to 0.293) | 92 | 10,681,107 | 3.265 | −0.291 (−0.665 to 0.187) | 0.988 | 0.217 (−0.242 to 0.635) |
| *GSDMD* | Border zone | 2 | 156 | 7,568,672 | 4.434 | 0.107 (—) | 47 | 4,949,362 | 3.392 | −0.164 (—) | 1.042 | 0.271 (—) |
| *GSDMD* | Infarct zone | 3 | 107 | 11,681,469 | 3.345 | −0.982 (−1.334 to −0.405) | 34 | 3,750,265 | 3.331 | −0.225 (−0.725 to 0.212) | 0.013 | −0.757 (−1.126 to 0.051) |

(B) Reference-gene calibration of *CASP4*

| Measure | Comparison | *CASP4* value | Percentile among 19,169 reference genes (predicted direction) | Fraction of 2,317 accessibility-matched reference genes with changes equal to that of *CASP4* or larger in the predicted direction |
|---|---|---|---|---|
| Pericyte difference | Overall AMI | 0.062 | 67.1 | 0.315 |
| Pericyte difference | Remote myocardium | −0.055 | 42.4 | 0.433 |
| Difference in ΔA | Overall AMI | −0.205 | 77.5 | 0.260 |
| Difference in ΔA | Remote myocardium | −0.201 | 79.0 | 0.266 |

A = log2($10^6$ × N/T + 1). N is the number of nuclei with fragments in 5 kb tiles overlapping the region within 1,000 bases on either side of each gene's MANE Select transcription start site. T is the sum across nuclei of the number of tiles containing fragments. The unit is the patient for controls and overall AMI, and the patient–region combination within strata. Controls comprise four patients and overall AMI five, with four units in remote myocardium, two in the border zone and three in the infarct zone. Differences are overall AMI or stratum values minus controls. ΔA = A(endothelium) − A(pericyte). Percentile 95% bootstrap intervals used 10,000 within-group resamples of units with replacement, only when both groups had at least three units. Units were patients for the overall AMI–control comparison, and the border zone had two units, hence '—'. The predicted directions are a positive pericyte *CASP4* difference and a negative difference in *CASP4* ΔA. Intervals are unadjusted for multiplicity, and no tests were performed.

#### Supplementary Table S9. Discovery records and endothelial-minus-pericyte differences in the confirmation set for ten other pyroptosis-related genes

The difference is ΔL = L(EC) − L(PC). Rows for fractions of positive nuclei show Δp in percentage points. In (A), discovery values come from the discovery genome-wide calibration table (Supplementary Data 1), with intervals from 1,000 bootstrap resamples, compared with 10,000 for the manuscript's main summaries. Directional fractions are as recorded in the calibration table: zero for records with a median of zero. Discovery records failing the precision criteria were classified as undeterminable. Those meeting the precision criteria but failing the remaining conditions were classified as not supported. In (B), percentile intervals used 10,000 donor resamples stratified by dataset, with the same saved resampling indices as in the initial confirmation. Only *CARD8* had a prediction and was assessed against the prespecified confirmation criteria, which it met. The other nine genes were described without predictions. In (C), values are medians across donor–class groups in the primary analysis, and fractions of positive nuclei are fractions with count ≥1.

(A) Discovery records and predictions

| Gene | Category | SCP1303 median | SCP1303 interval width | SCP1303 directional fraction | GSE183852 median | GSE183852 interval width | GSE183852 directional fraction | Discovery classification | Prediction |
|---|---|---|---|---|---|---|---|---|---|
| *CARD8* | Sensor | 2.646 | 0.384 | 1.000 | 2.881 | 0.339 | 1.000 | Supported | Endothelial side |
| *NLRP1* | Sensor | 4.867 | 1.017 | 1.000 | 5.050 | 0.828 | 1.000 | Undeterminable | None |
| *NLRP3* | Sensor | 0.000 | 0.000 | 0.000 | −0.102 | 0.428 | 0.700 | Not supported | None |
| *NLRC4* | Sensor | 0.499 | 1.042 | 0.694 | 0.111 | 1.017 | 0.594 | Undeterminable | None |
| *AIM2* | Sensor | 2.517 | 1.053 | 0.952 | 3.826 | 1.829 | 0.973 | Undeterminable | None |
| *MEFV* | Sensor | 0.000 | 0.000 | 0.000 | 0.000 | 0.017 | 0.000 | Not supported | None |
| *PYCARD* | Adaptor | 0.000 | 0.320 | 0.000 | 0.834 | 0.686 | 0.763 | Not supported | None |
| *IL1B* | Substrate | 0.000 | 0.000 | 0.000 | 0.000 | 0.000 | 0.000 | Undeterminable | None |
| *GSDME* | Pore formation and membrane rupture | — | — | — | — | — | — | Absent from calibration table | None |
| *NINJ1* | Pore formation and membrane rupture | 0.000 | 0.305 | 0.000 | 0.219 | 0.540 | 0.622 | Not supported | None |

(B) Primary and secondary analyses in the confirmation set

| Gene | Analysis | N | Endothelial side | Pericyte side | Ties | Median (95% interval) |
|---|---|---|---|---|---|---|
| *CARD8* | Primary analysis (22 donors) | 22 | 22 | 0 | 0 | 2.728 (2.564–3.001) |
| *CARD8* | Capillary endothelium and pericytes | 22 | 22 | 0 | 0 | 2.960 (2.663–3.136) |
| *CARD8* | Exclusion of predefined '-like' states | 22 | 22 | 0 | 0 | 2.724 (2.566–3.003) |

| Gene | Analysis | N | Endothelial side | Pericyte side | Ties | Median (95% interval) |
|---|---|---|---|---|---|---|
| *CARD8* | Right ventricular | 19 | 19 | 0 | 0 | 2.990 (2.572–3.299) |
| *CARD8* | Atrial | 15 | 15 | 0 | 0 | 2.592 (2.308–3.076) |
| *CARD8* | Reichart cardiomyopathy | 52 | 52 | 0 | 0 | 2.773 (2.565–2.918) |
| *CARD8* | Difference in positive-nucleus fraction (percentage points) | 22 | 22 | 0 | 0 | 18.549 (12.801–23.469) |
| *CARD8* | Heart Cell Atlas v2 | 16 | 16 | 0 | 0 | 2.728 (2.474–3.012) |
| *CARD8* | Reichart | 6 | 6 | 0 | 0 | 2.850 (2.518–3.128) |
| *NLRP1* | Primary analysis (22 donors) | 22 | 22 | 0 | 0 | 4.285 (3.887–4.747) |
| *NLRP1* | Capillary endothelium and pericytes | 22 | 22 | 0 | 0 | 4.416 (4.001–4.829) |
| *NLRP1* | Exclusion of predefined '-like' states | 22 | 22 | 0 | 0 | 4.277 (3.920–4.737) |
| *NLRP1* | Right ventricular | 19 | 19 | 0 | 0 | 4.373 (3.848–5.514) |
| *NLRP1* | Atrial | 15 | 15 | 0 | 0 | 3.047 (2.234–4.015) |
| *NLRP1* | Reichart cardiomyopathy | 52 | 52 | 0 | 0 | 4.290 (4.035–4.450) |
| *NLRP1* | Difference in positive-nucleus fraction (percentage points) | 22 | 22 | 0 | 0 | 19.078 (17.586–22.449) |
| *NLRP1* | Heart Cell Atlas v2 | 16 | 16 | 0 | 0 | 4.218 (3.760–4.990) |
| *NLRP1* | Reichart | 6 | 6 | 0 | 0 | 4.359 (3.939–5.219) |
| *NLRP3* | Primary analysis (22 donors) | 22 | 4 | 11 | 7 | −0.065 (−0.552 to 0.000) |
| *NLRP3* | Capillary endothelium and pericytes | 22 | 5 | 10 | 7 | 0.000 (−0.552 to 0.000) |
| *NLRP3* | Exclusion of predefined '-like' states | 22 | 4 | 11 | 7 | −0.065 (−0.565 to 0.000) |
| *NLRP3* | Right ventricular | 19 | 1 | 6 | 12 | 0.000 (−0.705 to 0.000) |
| *NLRP3* | Atrial | 15 | 5 | 4 | 6 | 0.000 (0.000–0.675) |
| *NLRP3* | Reichart cardiomyopathy | 52 | 16 | 22 | 14 | 0.000 (−0.307 to 0.000) |
| *NLRP3* | Difference in positive-nucleus fraction (percentage points) | 22 | 4 | 11 | 7 | −0.011 (−0.067 to 0.000) |
| *NLRP3* | Heart Cell Atlas v2 | 16 | 4 | 6 | 6 | 0.000 (−0.526 to 0.296) |
| *NLRP3* | Reichart | 6 | 0 | 5 | 1 | −0.549 (−0.803 to −0.065) |
| *NLRC4* | Primary analysis (22 donors) | 22 | 14 | 8 | 0 | 0.212 (−0.206 to 0.733) |

| Gene | Analysis | N | Endothelial side | Pericyte side | Ties | Median (95% interval) |
|---|---|---|---|---|---|---|
| *NLRC4* | Capillary endothelium and pericytes | 22 | 10 | 11 | 1 | −0.056 (−0.479 to 0.298) |
| *NLRC4* | Exclusion of predefined '-like' states | 22 | 15 | 7 | 0 | 0.329 (−0.172 to 0.751) |
| *NLRC4* | Right ventricular | 19 | 8 | 8 | 3 | 0.000 (−1.492 to 0.080) |
| *NLRC4* | Atrial | 15 | 7 | 5 | 3 | 0.000 (−0.151 to 1.154) |
| *NLRC4* | Reichart cardiomyopathy | 52 | 29 | 23 | 0 | 0.132 (−0.223 to 0.378) |
| *NLRC4* | Difference in positive-nucleus fraction (percentage points) | 22 | 12 | 10 | 0 | 0.048 (−0.050 to 0.147) |
| *NLRC4* | Heart Cell Atlas v2 | 16 | 11 | 5 | 0 | 0.632 (−0.105 to 0.858) |
| *NLRC4* | Reichart | 6 | 3 | 3 | 0 | −0.154 (−0.814 to 0.456) |
| *AIM2* | Primary analysis (22 donors) | 22 | 22 | 0 | 0 | 2.271 (1.914–2.806) |
| *AIM2* | Capillary endothelium and pericytes | 22 | 9 | 9 | 4 | 0.000 (−0.500 to 0.659) |
| *AIM2* | Exclusion of predefined '-like' states | 22 | 22 | 0 | 0 | 2.290 (1.913–2.823) |
| *AIM2* | Right ventricular | 19 | 13 | 2 | 4 | 2.162 (0.000–3.505) |
| *AIM2* | Atrial | 15 | 11 | 2 | 2 | 1.115 (0.000–2.776) |
| *AIM2* | Reichart cardiomyopathy | 52 | 50 | 1 | 1 | 2.947 (2.352–3.477) |
| *AIM2* | Difference in positive-nucleus fraction (percentage points) | 22 | 21 | 1 | 0 | 0.422 (0.344–0.754) |
| *AIM2* | Heart Cell Atlas v2 | 16 | 16 | 0 | 0 | 2.271 (1.914–2.863) |
| *AIM2* | Reichart | 6 | 6 | 0 | 0 | 2.141 (1.189–3.461) |
| *MEFV* | Primary analysis (22 donors) | 22 | 5 | 5 | 12 | 0.000 (0.000–0.000) |
| *MEFV* | Capillary endothelium and pericytes | 22 | 3 | 5 | 14 | 0.000 (0.000–0.000) |
| *MEFV* | Exclusion of predefined '-like' states | 22 | 5 | 5 | 12 | 0.000 (0.000–0.000) |
| *MEFV* | Right ventricular | 19 | 3 | 2 | 14 | 0.000 (0.000–0.000) |
| *MEFV* | Atrial | 15 | 2 | 1 | 12 | 0.000 (0.000–0.000) |
| *MEFV* | Reichart cardiomyopathy | 52 | 16 | 12 | 24 | 0.000 (0.000–0.000) |
| *MEFV* | Difference in positive-nucleus fraction (percentage points) | 22 | 5 | 5 | 12 | 0.000 (0.000–0.000) |
| *MEFV* | Heart Cell Atlas v2 | 16 | 5 | 5 | 6 | 0.000 (−0.248 to 0.326) |

| Gene | Analysis | N | Endothelial side | Pericyte side | Ties | Median (95% interval) |
|---|---|---|---|---|---|---|
| *MEFV* | Reichart | 6 | 0 | 0 | 6 | 0.000 (0.000–0.000) |
| *PYCARD* | Primary analysis (22 donors) | 22 | 18 | 1 | 3 | 1.205 (0.663–1.558) |
| *PYCARD* | Capillary endothelium and pericytes | 22 | 13 | 3 | 6 | 0.474 (0.000–1.311) |
| *PYCARD* | Exclusion of predefined '-like' states | 22 | 18 | 1 | 3 | 1.214 (0.670–1.516) |
| *PYCARD* | Right ventricular | 19 | 12 | 2 | 5 | 0.514 (0.000–1.565) |
| *PYCARD* | Atrial | 15 | 9 | 3 | 3 | 1.210 (0.000–2.147) |
| *PYCARD* | Reichart cardiomyopathy | 52 | 48 | 3 | 1 | 1.185 (0.976–1.307) |
| *PYCARD* | Difference in positive-nucleus fraction (percentage points) | 22 | 17 | 2 | 3 | 0.215 (0.094–0.535) |
| *PYCARD* | Heart Cell Atlas v2 | 16 | 13 | 0 | 3 | 1.229 (0.850–1.790) |
| *PYCARD* | Reichart | 6 | 5 | 1 | 0 | 0.487 (−0.374 to 1.471) |
| *IL1B* | Primary analysis (22 donors) | 22 | 3 | 2 | 17 | 0.000 (0.000–0.000) |
| *IL1B* | Capillary endothelium and pericytes | 22 | 3 | 2 | 17 | 0.000 (0.000–0.000) |
| *IL1B* | Exclusion of predefined '-like' states | 22 | 3 | 2 | 17 | 0.000 (0.000–0.000) |
| *IL1B* | Right ventricular | 19 | 0 | 0 | 19 | 0.000 (0.000–0.000) |
| *IL1B* | Atrial | 15 | 2 | 0 | 13 | 0.000 (0.000–0.000) |
| *IL1B* | Reichart cardiomyopathy | 52 | 6 | 7 | 39 | 0.000 (0.000–0.000) |
| *IL1B* | Difference in positive-nucleus fraction (percentage points) | 22 | 3 | 2 | 17 | 0.000 (0.000–0.000) |
| *IL1B* | Heart Cell Atlas v2 | 16 | 2 | 2 | 12 | 0.000 (0.000–0.000) |
| *IL1B* | Reichart | 6 | 1 | 0 | 5 | 0.000 (0.000–0.788) |
| *GSDME* | Primary analysis (22 donors) | 22 | 1 | 20 | 1 | −1.840 (−2.482 to −1.296) |
| *GSDME* | Capillary endothelium and pericytes | 22 | 0 | 21 | 1 | −1.937 (−2.548 to −1.199) |
| *GSDME* | Exclusion of predefined '-like' states | 22 | 1 | 20 | 1 | −1.854 (−2.511 to −1.272) |
| *GSDME* | Right ventricular | 19 | 3 | 14 | 2 | −2.091 (−2.801 to −0.878) |
| *GSDME* | Atrial | 15 | 3 | 9 | 3 | −0.418 (−1.212 to 0.000) |
| *GSDME* | Reichart cardiomyopathy | 52 | 2 | 50 | 0 | −2.256 (−2.500 to −1.875) |

| Gene | Analysis | N | Endothelial side | Pericyte side | Ties | Median (95% interval) |
|---|---|---|---|---|---|---|
| *GSDME* | Difference in positive-nucleus fraction (percentage points) | 22 | 1 | 20 | 1 | −0.634 (−0.788 to −0.347) |
| *GSDME* | Heart Cell Atlas v2 | 16 | 1 | 14 | 1 | −1.611 (−2.471 to −1.162) |
| *GSDME* | Reichart | 6 | 0 | 6 | 0 | −2.516 (−2.913 to −1.239) |
| *NINJ1* | Primary analysis (22 donors) | 22 | 10 | 11 | 1 | −0.002 (−0.335 to 0.640) |
| *NINJ1* | Capillary endothelium and pericytes | 22 | 9 | 12 | 1 | −0.087 (−0.628 to 0.276) |
| *NINJ1* | Exclusion of predefined '-like' states | 22 | 9 | 11 | 2 | −0.006 (−0.379 to 0.452) |
| *NINJ1* | Right ventricular | 19 | 7 | 6 | 6 | 0.000 (−0.958 to 0.083) |
| *NINJ1* | Atrial | 15 | 6 | 7 | 2 | 0.000 (−0.688 to 0.169) |
| *NINJ1* | Reichart cardiomyopathy | 52 | 31 | 20 | 1 | 0.179 (−0.041 to 0.398) |
| *NINJ1* | Difference in positive-nucleus fraction (percentage points) | 22 | 10 | 11 | 1 | −0.016 (−0.118 to 0.087) |
| *NINJ1* | Heart Cell Atlas v2 | 16 | 8 | 7 | 1 | 0.024 (−0.202 to 0.723) |
| *NINJ1* | Reichart | 6 | 2 | 4 | 0 | −0.379 (−0.995 to 0.577) |

(C) Detection in primary-analysis groups

| Gene | Dataset | Groups per class | Endothelium: groups with 0 counts | Endothelium: median count | Endothelium: positive-nucleus fraction (%) | Endothelium: median L | Pericytes: groups with 0 counts | Pericytes: median count | Pericytes: positive-nucleus fraction (%) | Pericytes: median L |
|---|---|---|---|---|---|---|---|---|---|---|
| *CARD8* | Pooled | 22 | 0 | 214.5 | 23.56 | 8.019 | 0 | 75 | 4.88 | 5.243 |
| *CARD8* | Heart Cell Atlas v2 | 16 | 0 | 187.5 | 16.43 | 7.759 | 0 | 121.5 | 4.62 | 5.262 |
| *CARD8* | Reichart | 6 | 0 | 263 | 33.28 | 8.086 | 0 | 71.5 | 6.56 | 5.243 |
| *NLRP1* | Pooled | 22 | 0 | 221.5 | 20.59 | 7.851 | 0 | 26 | 1.47 | 3.446 |
| *NLRP1* | Heart Cell Atlas v2 | 16 | 0 | 221.5 | 18.96 | 7.872 | 0 | 30.5 | 1.24 | 3.417 |
| *NLRP1* | Reichart | 6 | 0 | 214.5 | 29.51 | 7.838 | 0 | 15.5 | 1.64 | 3.467 |
| *NLRP3* | Pooled | 22 | 17 | 0 | 0.00 | 0.000 | 9 | 1 | 0.03 | 0.359 |
| *NLRP3* | Heart Cell Atlas v2 | 16 | 12 | 0 | 0.00 | 0.000 | 8 | 0.5 | 0.01 | 0.100 |
| *NLRP3* | Reichart | 6 | 5 | 0 | 0.00 | 0.000 | 1 | 1 | 0.12 | 0.727 |
| *NLRC4* | Pooled | 22 | 1 | 3 | 0.33 | 1.960 | 2 | 5 | 0.31 | 1.855 |
| *NLRC4* | Heart Cell Atlas v2 | 16 | 1 | 2 | 0.28 | 1.804 | 2 | 4.5 | 0.20 | 1.419 |

| Gene | Dataset | Groups per class | Endothelium: groups with 0 counts | Endothelium: median count | Endothelium: positive-nucleus fraction (%) | Endothelium: median L | Pericytes: groups with 0 counts | Pericytes: median count | Pericytes: positive-nucleus fraction (%) | Pericytes: median L |
|---|---|---|---|---|---|---|---|---|---|---|
| *NLRC4* | Reichart | 6 | 0 | 5 | 0.59 | 2.174 | 0 | 7.5 | 0.70 | 2.350 |
| *AIM2* | Pooled | 22 | 0 | 6 | 0.52 | 2.708 | 5 | 1 | 0.08 | 0.628 |
| *AIM2* | Heart Cell Atlas v2 | 16 | 0 | 5.5 | 0.52 | 2.776 | 4 | 1.5 | 0.07 | 0.645 |
| *AIM2* | Reichart | 6 | 0 | 6 | 0.52 | 2.475 | 1 | 1 | 0.11 | 0.614 |
| *MEFV* | Pooled | 22 | 17 | 0 | 0.00 | 0.000 | 17 | 0 | 0.00 | 0.000 |
| *MEFV* | Heart Cell Atlas v2 | 16 | 11 | 0 | 0.00 | 0.000 | 11 | 0 | 0.00 | 0.000 |
| *MEFV* | Reichart | 6 | 6 | 0 | 0.00 | 0.000 | 6 | 0 | 0.00 | 0.000 |
| *PYCARD* | Pooled | 22 | 4 | 2 | 0.36 | 2.051 | 7 | 2 | 0.14 | 0.897 |
| *PYCARD* | Heart Cell Atlas v2 | 16 | 3 | 3 | 0.36 | 2.163 | 7 | 1.5 | 0.09 | 0.756 |
| *PYCARD* | Reichart | 6 | 1 | 1.5 | 0.42 | 1.782 | 0 | 2 | 0.17 | 0.958 |
| *IL1B* | Pooled | 22 | 19 | 0 | 0.00 | 0.000 | 20 | 0 | 0.00 | 0.000 |
| *IL1B* | Heart Cell Atlas v2 | 16 | 14 | 0 | 0.00 | 0.000 | 14 | 0 | 0.00 | 0.000 |
| *IL1B* | Reichart | 6 | 5 | 0 | 0.00 | 0.000 | 6 | 0 | 0.00 | 0.000 |
| *GSDME* | Pooled | 22 | 9 | 1 | 0.07 | 0.640 | 1 | 13.5 | 0.75 | 2.508 |
| *GSDME* | Heart Cell Atlas v2 | 16 | 5 | 1 | 0.08 | 0.843 | 1 | 14.5 | 0.51 | 2.427 |
| *GSDME* | Reichart | 6 | 4 | 0 | 0.00 | 0.000 | 0 | 8 | 0.83 | 2.575 |
| *NINJ1* | Pooled | 22 | 3 | 2 | 0.33 | 1.682 | 1 | 6 | 0.39 | 2.055 |
| *NINJ1* | Heart Cell Atlas v2 | 16 | 3 | 3 | 0.27 | 1.807 | 1 | 6 | 0.28 | 2.054 |
| *NINJ1* | Reichart | 6 | 0 | 2 | 0.34 | 1.563 | 0 | 7 | 0.73 | 2.246 |

#### Supplementary Table S10. Additional single-nucleus ATAC checks in six donors (planned after confirmation)

(A) Endothelial–pericyte differences in gene-body accessibility

| Gene | Predicted direction | Ventricles: donors in the predicted direction | Ventricles: median difference | Ventricles: reference-gene percentile (predicted direction) | All sampled heart regions: donors in the predicted direction | All sampled heart regions: median difference |
|---|---|---|---|---|---|---|
| *IL18* | Pericytes | 2/6 | 0.276 | 20.8 | 1/6 | 0.282 |
| *CASP1* | Endothelium | 6/6 | 0.726 | 93.4 | 6/6 | 0.795 |
| *CASP4* | Endothelium | 6/6 | 0.847 | 94.8 | 6/6 | 0.821 |
| *GSDMD* | Endothelium | 6/6 | 0.942 | 95.6 | 5/6 | 1.001 |
| *IL18BP* | Endothelium | 3/6 | −0.113 | 43.9 | 1/6 | −0.366 |

(B) *IL18* accessibility differences involving macrophages

| Measurement region | Comparison | Ventricles: donors with positive differences | Ventricles: median difference | Ventricles: reference-gene percentile | All sampled heart regions: donors with positive differences | All sampled heart regions: median difference |
|---|---|---|---|---|---|---|
| Around transcription start sites | Macrophage-minus-endothelial | 6/6 | 2.769 | 99.3 | 6/6 | 2.594 |
| Around transcription start sites | Macrophage-minus-pericyte | 6/6 | 2.444 | 98.8 | 6/6 | 2.486 |
| Gene body | Macrophage-minus-endothelial | 6/6 | 2.268 | 99.1 | 6/6 | 2.216 |
| Gene body | Macrophage-minus-pericyte | 6/6 | 2.477 | 99.3 | 6/6 | 2.487 |

(C) Relationship between RNA differences in the confirmation set and gene-body accessibility differences (18,564 genes, Spearman $\rho = 0.502$, line $\Delta L = 0.194 + 1.297 \times \Delta A$)

| Gene | RNA median ΔL (22 donors) | Accessibility median ΔA (6 donors) | Residual from the fitted line | Residual rank |
|---|---|---|---|---|
| *IL18* | −1.451 | 0.276 | −2.003 | Top 2.3% towards pericytes |
| *CASP1* | 3.020 | 0.726 | 1.883 | Top 3.7% towards endothelium |
| *CASP4* | 2.395 | 0.847 | 1.102 | Top 8.9% towards endothelium |
| *GSDMD* | 1.762 | 0.942 | 0.346 | Top 26.7% towards endothelium |

(D) RNA differences in the confirmation set for genes with macrophage-specific accessibility

| Measure | 107 genes with macrophage-specific accessibility | All RNA genes (32,657) |
|---|---|---|
| Median ΔL | 0.000 | 0.000 |
| Fraction of genes favouring pericytes in at least 18 of 22 donors | 4.7% | 10.8% |

(E) Tiles with pericyte-specific accessibility within 100 kb on either side of the *IL18* transcription start site (among 41 tiles)

| Tile (GRCh38) | Location | Median endothelial-minus-pericyte difference | Median macrophage-minus-pericyte difference |
|---|---|---|---|
| chr11:112,170,001–112,175,000 | Approximately 6–11 kb upstream of the transcription start site, overlapping the 3′ region of *TEX12* | −0.872 | −0.021 |
| chr11:112,230,001–112,235,000 | Approximately 66–71 kb upstream of the transcription start site, within the *PTS* gene | −0.570 | −0.523 |

Accessibility in 5 kb tiles was defined as $A = \log_2(10^6 \times N/T + 1)$. For accessibility around transcription start sites, N is the number of nuclei with signal in tiles overlapping the region within 1,000 bases on either side of the MANE Select transcription start site. For gene-body accessibility, N is the summed number across nuclei of signal-positive tiles overlapping the region from 2,000 bases upstream of the transcription start site to the transcript end. T is the summed number of signal-positive tiles across nuclei. Median differences are across six donors, with positive values favouring endothelium, or macrophages in (B). The reference set comprises 19,169 genes. Values are descriptive, and no tests were performed. (B) Under the rule specified before viewing the values, *IL18* accessibility was recorded as measurable because both differences around the transcription start site were positive in all six donors. (C) RNA ΔL was calculated in the same way as in the confirmation analysis and is the median across 22 donors. Residuals are observed ΔL minus the line's prediction. (D) The genes with macrophage-specific accessibility are the 107 matched to RNA among 108 genes with both macrophage-minus-endothelial and macrophage-minus-pericyte median gene-body differences in the top 1% of reference genes. The interpretation specified before viewing the values was 'no indication' if the median for the 107 genes lay within ±0.2 of the genome-wide median. *IL18* ΔL lay further towards pericytes than that of 97.2% of these 107 genes. (E) Tiles with pericyte-specific accessibility were defined by a negative endothelial-minus-pericyte difference in all six donors, a median ≤−0.5, and a median macrophage-minus-pericyte difference <0.5. The tile at the *IL18* transcription start site and the tile upstream of the second exon failed to meet these criteria. Of 516,617 genome-wide tiles matched for signal range, 4.2% met the criteria, giving an expectation of 1.7 among 41 tiles. Tile positions and neighbouring genes follow MANE Select 1.4.

#### Supplementary Table S11. Comparison of deposited experimental structures

(A) Structures used

| PDB | Molecule | Method | Copies/models used | Reference |
|---|---|---|---|---|
| 8URV | Pro-IL-18 (residues 1–193) | Solution NMR | 10 models | [43] |
| 1J0S | Mature IL-18 | Solution NMR | 20 models | [44] |
| 7AL7 | IL-18–IL-18BP | X-ray, 1.80 Å | 1 copy | [41] |
| 3WO2 | IL-18 | X-ray, 2.33 Å | 4 copies | [42] |
| 3WO3 | IL-18–IL-18Rα | X-ray, 3.10 Å | 6 copies | [42] |
| 3WO4 | IL-18–IL-18Rα–IL-18Rβ | X-ray, 3.10 Å | 1 copy | [42] |

(B) Numbers of recognition residues on mature IL-18

| Threshold | IL-18BP | IL-18Rα | IL-18Rβ | Shared by IL-18BP and IL-18Rα | In either or both sets |
|---|---|---|---|---|---|
| 4.0 Å | 21 | 32 | 9 | 12 | 41 |
| 4.5 Å (primary) | 23 | 40 | 11 | 16 | 47 |
| 5.0 Å | 25 | 42 | 11 | 17 | 50 |

The 16 shared residues (UniProt numbering) are 37, 41, 42, 87, 88, 89, 90, 91, 93, 94, 95, 96, 139, 140, 141 and 149.

(C) Hypothetical overlay of binding poses superposed using mature IL-18 (157 Cα atoms, RMSD 1.27 Å)

| Protein pair | Atom pairs below 2 Å | Below 2.5 Å | Below 3 Å | Minimum distance |
|---|---|---|---|---|
| IL-18BP and IL-18Rα | 1,199 | 2,249 | 3,835 | 0.25 Å |
| IL-18BP and IL-18Rβ | 23 | 57 | 100 | 0.14 Å |

Of atom pairs separated by less than 2 Å, 520 were backbone–backbone pairs, all between IL-18BP and IL-18Rα.

(D) Receptor stages

| Item | Value |
|---|---|
| RMSD after superposition on 3WO4, IL-18 alone (3WO2, 4 copies, 156 Cα atoms) | 0.66–0.74 Å |
| RMSD after superposition on 3WO4, complex with IL-18Rα (3WO3, 6 copies, 156 Cα atoms) | 0.37–0.45 Å |
| IL-18Rβ contact residues in 3WO4 | IL-18 only 6, IL-18Rα only 15, both 4 |

(E) Differences in Cα distances between recognition residues (root-mean-square differences relative to bound structures, Å)

| Recognition residues | Range | Residues | Residue pairs | Pro-IL-18 8URV, median (minimum–maximum) | Mature IL-18 1J0S, median (minimum–maximum) |
|---|---|---|---|---|---|
| IL-18BP | All | 23 | 219 | 12.948 (12.199–13.736) | 1.885 (1.559–2.147) |
| IL-18BP | Well-ordered ranges | 9 | 30 | 3.208 (3.108–3.291) | 0.842 (0.746–1.038) |
| IL-18Rα | All | 40 | 722 | 8.834 (8.565–9.077) | 2.786 (2.182–3.177) |
| IL-18Rα | Well-ordered ranges | 15 | 96 | 6.199 (6.070–6.284) | 1.496 (1.082–1.773) |
| IL-18Rβ | All | 11 | 40 | 7.205 (6.870–7.468) | 1.812 (1.307–2.303) |
| IL-18Rβ | Well-ordered ranges | 7 | 14 | 6.002 (5.738–6.338) | 0.878 (0.480–1.585) |

(F) Solvent-accessible surface area of recognition residues ($Å^2$, median across models)

| Recognition residues | Pro-IL-18 8URV | Pro-IL-18 with the pro region removed | Mature IL-18 1J0S | Median per-model difference (minimum–maximum) |
|---|---|---|---|---|
| IL-18BP | 1,305.0 | 1,614.4 | 1,568.9 | 306.3 (217.0–375.9) |
| IL-18Rα | 3,374.5 | 3,815.2 | 3,601.9 | 422.5 (375.5–477.2) |
| IL-18Rβ | 817.0 | 867.6 | 873.1 | 39.7 (30.6–195.9) |

The difference is the value for the state with the pro region removed minus that for pro-IL-18. The difference was positive in all 10 models.

#### Supplementary Figure Legends

##### Supplementary Figure S1

Capillary-restricted endothelial–pericyte differences (ΔL) for *IL18*, *CASP1*, *CASP4* and *GSDMD* in the 22 primary confirmation donors (Heart Cell Atlas v2, n = 16, and Reichart, n = 6). Points represent donors. Summary points and bars are dataset-specific medians and 95% donor-bootstrap intervals. The dashed line marks zero. All four genes show their predicted direction in every donor.

##### Supplementary Figure S2

Endothelial–pericyte differences after exclusion of the predefined EC9_FB-like, EC10_CMC-like and PC4_CMC-like annotations in the 22 primary confirmation donors (Heart Cell Atlas v2, n = 16, and Reichart, n = 6). Plot elements are as in Supplementary Figure S1. All four genes show their predicted direction in every donor.

##### Supplementary Figure S3

Right-ventricular endothelial–pericyte differences in 19 donors (Heart Cell Atlas v2, n = 16, and Reichart, n = 3). *IL18* is negative in 18 donors with one tie. *CASP1* is positive in 15 with four ties. *CASP4* and *GSDMD* are positive in all 19. Ties lie on the dashed zero line. Plot elements are as in Supplementary Figure S1.

##### Supplementary Figure S4

Atrial endothelial–pericyte differences in 15 Heart Cell Atlas v2 donors. No Reichart donor met the atrial eligibility criteria. *IL18* is negative in 12 donors, positive in two and tied in one. *CASP1* is positive in 14 with one tie; *CASP4* and *GSDMD* are positive in all 15. Plot elements are as in Supplementary Figure S1.

##### Supplementary Figure S5

Endothelial–pericyte differences in the 52 Reichart cardiomyopathy donors, a set separate from the 22 primary confirmation donors. *IL18* is negative in 51 donors and positive in one. *CASP1*, *CASP4* and *GSDMD* are positive in all 52. Plot elements are as in Supplementary Figure S1.

##### Supplementary Figure S6

Difference in the fraction of nuclei with at least one count (Δp, endothelium minus pericyte) in the 22 primary confirmation donors (Heart Cell Atlas v2, n = 16, and Reichart, n = 6). All four genes show their predicted direction in every donor. Plot elements are as in Supplementary Figure S1.

##### Supplementary Figure S7

Dataset-specific primary results for the 22 primary confirmation donors, shown separately for Heart Cell Atlas v2 (n = 16) and Reichart (n = 6). All four genes show their predicted direction in every donor of each dataset. Plot elements are as in Supplementary Figure S1.

##### Supplementary Figure S8

Endothelial and pericyte nuclei of the six Reichart confirmation donors, 3,871 endothelial and 6,721 pericyte nuclei, in a shared embedding of the principal components distributed with the resource. Plot elements and colour scales are as in Figure 1B and 1C. Grey bins contain fewer than 15 nuclei.

##### Supplementary Figure S9

Bin values from the embeddings in Figure 1B and 1C at three grid sizes, 40, 60 and 80 bins across the horizontal range, for SCP1303 (A) and Heart Cell Atlas v2 (B). Each row gives the grid size, the number of coloured bins and the coverage, the fraction of nuclei in coloured bins. Bold rows mark the grid used in Figure 1B or 1C, the finest with coverage of at least 0.80. Grey bins contain fewer than 30 nuclei. Colour scales are shared by Supplementary Figures S9 and S10.

#### Supplementary Figure S10

Bin values of Supplementary Figure S8 at 25, 35 and 50 bins across the horizontal range, shown as in Supplementary Figure S9. Grey bins contain fewer than 15 nuclei.

#### Supplementary Figure S11. Sensitivity analyses and comparisons with smooth muscle

(A) Endothelial-minus-pericyte medians and 95% bootstrap intervals for *IL18*, *CASP1*, *CASP4* and *GSDMD* by source disease group. Numbers in parentheses are donor counts. Background shading marks the side of zero, with orange favouring pericytes and blue favouring endothelium. (B) Medians and 95% bootstrap intervals of endothelial-minus-smooth-muscle and pericyte-minus-smooth-muscle differences for the four genes in both cohorts. Numbers in parentheses are donor counts. For the pericyte-minus-smooth-muscle *CASP1* contrast in SCP1303, 24 of 42 donors were tied, giving a median and interval of zero. (C) Frequency of meeting the single-cohort precision and support criteria across 200 within-donor count-thinning replicates. Each donor's two libraries were reduced to the expected depth of the smaller library. The frequency for *IL18* in GSE183852 was 55.5%, and the other displayed frequencies were 100%. Results for disease strata, leave-one-donor-out analyses, nucleus-count thresholds, denominators, markers and mural-cell comparisons are provided in Supplementary Data 1. DCM, dilated cardiomyopathy; EC, endothelium; HCM, hypertrophic cardiomyopathy; PC, pericyte; SMC, smooth muscle.

#### Supplementary Figure S12. Endothelial–cardiomyocyte comparison of *GSDMD*

(A–C) Donor-level endothelial-minus-cardiomyocyte *GSDMD* differences in SCP1303 (42 donors), the GSE183852 nuclei subset (38 donors) and GSE271676 GeoMx (32 donors). Points represent individual donors with uniform vertical offsets. Diamonds and horizontal bars below the points show medians and exact 95% CIs. Sequencing panels show differences in log2(CPM + 1), and the GeoMx panel shows differences in upper-quartile-normalised z-score units. Positive differences occurred in 42/42, 38/38 and 27/32 donors, in the same cohort order. The annotated percentiles are within-cohort signed-contrast ranks. (D) GeoMx composition diagnostic, showing the intercept and regression coefficients for the myeloid and stromal marker scores, with HC3 95% confidence intervals. (E) GSDMD protein abundance in one healthy human heart from Human Protein Atlas Deep Visual Proteomics, expressed as the base-10 logarithm of the intensities reported by the source. Points represent five sites in each laser-microdissected compartment, and vertical lines mark medians. The capillary compartment is endothelial-rich and was dissected using CD34 as a marker. CASP4 was not detected at any of the five sites in either compartment.

#### Supplementary Figure S13. Ischaemic-heart *CASP4* changes compared with expression-matched genes

(A) Acute myocardial infarction-minus-control differences (AMI − control) in GSE270788 and Kuppe. GSE270788 included six control and five AMI donors, and Kuppe included four control and seven AMI donors. The left panel ("Pericyte level, L(PC)") shows the difference in pericyte L. The right panel ("ΔL = L(EC) − L(PC)") shows the AMI-minus-control difference in the endothelial–pericyte contrast. Both were calculated from the same count-pooled L values as in Figure 6B. Navy diamonds show the *CASP4* values. Box plots show the distributions of the corresponding quantities for reference genes. Boxes span the 25th–75th percentiles, the line within each box marks the median, and whiskers span the 5th–95th percentiles. Reference genes were selected for pericyte abundance close to that of *CASP4* (Supplemental Methods). There were 2,364 reference genes in GSE270788 and 3,066 in Kuppe, with the same genes used in the left and right panels. Numbers to the right of the left panel ("% ≥ *CASP4*") give the percentage of reference genes with changes at least as large as that of *CASP4*. Numbers to the right of the right panel ("% ≤ *CASP4*") give the percentage with changes at most as large as that of *CASP4*. Percentages were rounded to whole numbers. (B) Changes from controls in the AMI regions and ICM in Kuppe, using the same values as the left part of Figure 6C. Rows and patient counts in parentheses in the row labels are the same as in Figure 6C. The left panel ("Pericyte level, L(PC)") shows changes in pericyte L, and the right panel ("Endothelial level, L(EC)") shows changes in endothelial L. Navy diamonds and box plots are displayed as in A. Reference genes were selected separately for each row and class for abundance close to that of *CASP4* in that class (Supplemental Methods). Reference-gene counts, in pericyte/endothelial order, were 3,063/1,968 in remote myocardium, 3,094/2,014 in the border zone, 3,088/2,149 in the infarct zone and 3,024/1,860 in ICM. Numbers at right give the same percentages as in A, with "% ≥ *CASP4*" for pericyte panels and "% ≤ *CASP4*" for endothelial panels. The horizontal-axis range is shared by A and B, and the vertical line marks zero. (C) The contrast ratio κ for ΔL = L(EC) − L(PC) relative to controls in each Kuppe unit. A unit is a patient–region combination in AMI and one specimen from one individual in ICM and controls. The ratio κ was calculated by dividing the unit's ΔL by the ΔL calculated from count-pooled L for the four controls. The four control units were divided by the same value. Navy diamonds show *CASP4* κ. Grey open circles show the median κ calculated in the same unit for the 307 endothelial-biased reference genes in Figure 6C ("Matched EC-biased genes, median"). A grey vertical line joins the diamond and circle for each unit. The horizontal dashed line at κ = 1 marks a difference equal to that of the pooled controls. Units are grouped into control, remote myocardium, border zone, infarct zone and ICM sections, ordered within each section by the numerical part of the original identifiers. Numbers in parentheses in the section labels give the number of units. The 23 units eligible for both endothelium and pericytes are shown. Pale grey vertical lines mark section boundaries. AMI, acute myocardial infarction; EC, endothelium; ICM, ischaemic cardiomyopathy; PC, pericyte.

#### Supplementary Figure S14. Contamination checks for pericyte *CASP4* changes in acute myocardial infarction

The left column shows GSE270788 and the right column Kuppe. The labels "AMI 5 vs control 6" and "AMI 7 vs control 4" above the columns give the numbers of AMI and control donors with eligible pericytes, requiring at least 30 nuclei and a positive library total. In Kuppe, nuclei from all regions and specimens of each donor were pooled, as in Figure 6B. The same donors were used in A, B and the two "all donors" rows in C. The vertical line marks zero. (A) Marker-based contamination model. Specimen-level *CASP4* counts attributable to contamination were estimated from the counts of five myeloid markers in pericyte nuclei and the ratio of *CASP4* to marker counts in the contamination source from the same specimen. In GSE270788, each donor contributed one specimen (Supplemental Methods). The upper set of rows ("Source: all nuclei") used all nuclei with public annotations from that specimen as the contamination source. The lower set of rows ("Source: myeloid nuclei") used the specimen's myeloid nuclei. Grey open circles ("Observed (ΔO)") show the AMI-minus-control difference in *CASP4* CPM across pooled pericyte nuclei. Black diamonds ("Observed − expected (Δ)") show the AMI-minus-control difference after subtracting the estimated contamination-derived CPM from the observed CPM in each group. (B) Subtraction of contributions from other classes. Genes specific to each of four classes, myeloid cells, endothelium, fibroblasts and cardiomyocytes, were used to estimate the contribution of each class to pericyte *CASP4* (Supplemental Methods). Endothelium in GSE270788 included endocardium and lymphatic endothelium. The label "3 classes (myeloid n/a)" above Kuppe indicates estimation from the three classes other than myeloid cells: endothelium, fibroblasts and cardiomyocytes (Supplemental Methods). Grey open circles ("Observed (ΔO)") show the observed difference. Grey squares ("Expected from other classes (ΔE)") show the difference expected from contamination by these classes. Black diamonds ("Observed − expected (Δ2)") show the observed difference minus the expected difference. The horizontal axes in A and B show CPM differences (AMI − control). Their ranges differ between cohorts but are shared by A and B within each cohort. (C) Nucleus exclusion and leave-one-donor-out estimates. Values are AMI-minus-control differences in count-pooled pericyte *CASP4* L, where L = log2(1 + CPM). "All nuclei, all donors" gives the estimate using all nuclei and all donors, with the same value and interval as the pericyte row in Figure 6B. The thick grey band for "All nuclei, leave one donor out" spans the minimum and maximum of the 11 estimates obtained by leaving out one donor at a time. The open diamond for "All nuclei, batch 3 excluded" gives the estimate for five GSE270788 donors, three controls and two AMI donors, after excluding donors from processing batch 3, and is shown without an interval. This row is not applicable to Kuppe ("n/a"). "After nucleus filter, all donors" gives the estimate after excluding pericyte nuclei with a total count of at least 1 across the five myeloid markers or at least 2 across the three endothelial markers and reassessing eligibility. The horizontal-axis range in C is shared by the two cohorts. Horizontal bars in A–C are percentile 95% bootstrap intervals from 10,000 donor resamples with replacement within groups. In A and B, values in the same set of rows used the same resamples. AMI, acute myocardial infarction; CPM, counts per million library counts; n/a, not available.

#### Supplementary Figure S15. Class-level changes in *IL18*, *CASP1* and *GSDMD* in acute myocardial infarction and class contributions to *IL18* counts

(A) Class-level changes, L(AMI) − L(control), in *IL18*, *CASP1* and *GSDMD* in GSE270788 and Kuppe. L is the same count-pooled L used in Figure 6B. Row-to-class assignments, interval calculation and eligibility for interval calculation, and the pale grey band on the pericyte row are as in Figure 6B (Methods, Supplemental Methods). "too few nuclei" indicates that at least one group had no donor with 30 or more nuclei in that class. For each class, the upper row shows GSE270788 in black and the lower row Kuppe in grey. Diamonds and horizontal bars show changes and 95% bootstrap intervals. An open diamond denotes the estimate without an interval for GSE270788 adipocytes, for which there was one control donor. The horizontal-axis range is shared by the three genes. The dark vertical line marks zero, pale vertical lines mark other tick positions, and thin horizontal lines separate classes. (B) Bar lengths show each class's share of *IL18* counts. All donors in each group were included: seven controls and five AMI donors in GSE270788, and four controls and seven AMI donors in Kuppe. *IL18* counts were summed across all nuclei in each class, regardless of nucleus-count eligibility, and expressed as a fraction of the group total. Bar colours indicate classes, with the colour key below the figure. The epicardial class is present only in GSE270788. In addition to pericytes, endothelium and myeloid cells, classes contributing at least 10% of any of the four bars are named, and the remaining classes are combined as "Other". Endocardium and lymphatic endothelium in GSE270788 are included in "Other". Kuppe's Endothelium corresponds to the source annotation Endothelial and includes endocardium and lymphatic endothelium. The right-hand column ("*IL18* counts") gives total *IL18* counts for each group. AMI, acute myocardial infarction; T/NK, T and NK cells.

#### Supplementary Figure S16. Comparison of endothelial and pericyte chromatin accessibility around transcription start sites in single-nucleus ATAC data from six donors

Cardiac single-nucleus ATAC data from six CATlas donors, comprising four CARE and two ENTEx donors. For each gene, the presence or absence of signal in 5 kb tiles overlapping the region within 1 kb on either side of the transcription start site was scored per nucleus and summed within each donor and class. The difference is endothelial-minus-pericyte accessibility on a log2 scale. (A) Ventricular median differences for all genes, comprising 19,169 reference genes and five target genes, ordered from smallest to largest. Shading marks the 5% at each end. (B) Median differences and the number of donors favouring endothelium out of six for the same genes. Grey points represent reference genes and were offset vertically. The 247 reference genes outside the horizontal-axis range are not shown. The dotted line marks the 95th percentile of median differences across all genes. (C, D) Donor-level differences for the five genes in ventricles (C) and all sampled heart regions (D). Circles represent CARE donors and triangles ENTEx donors, filled blue for positive differences and orange for negative differences. Open symbols denote donors with zero signal-positive nuclei in either endothelium or pericytes, and pale fills denote donors with 1–3 signal-positive nuclei in either class. Vertical lines mark medians, and grey bands span the central 50% of reference-gene medians. Numbers to the right of gene names give the number of donors in the predicted direction divided by six, with EC indicating the endothelial direction and PC the pericyte direction. The predicted direction for *IL18BP* is towards endothelium. EC, endothelium; PC, pericyte.

**Supplementary Figure S1**

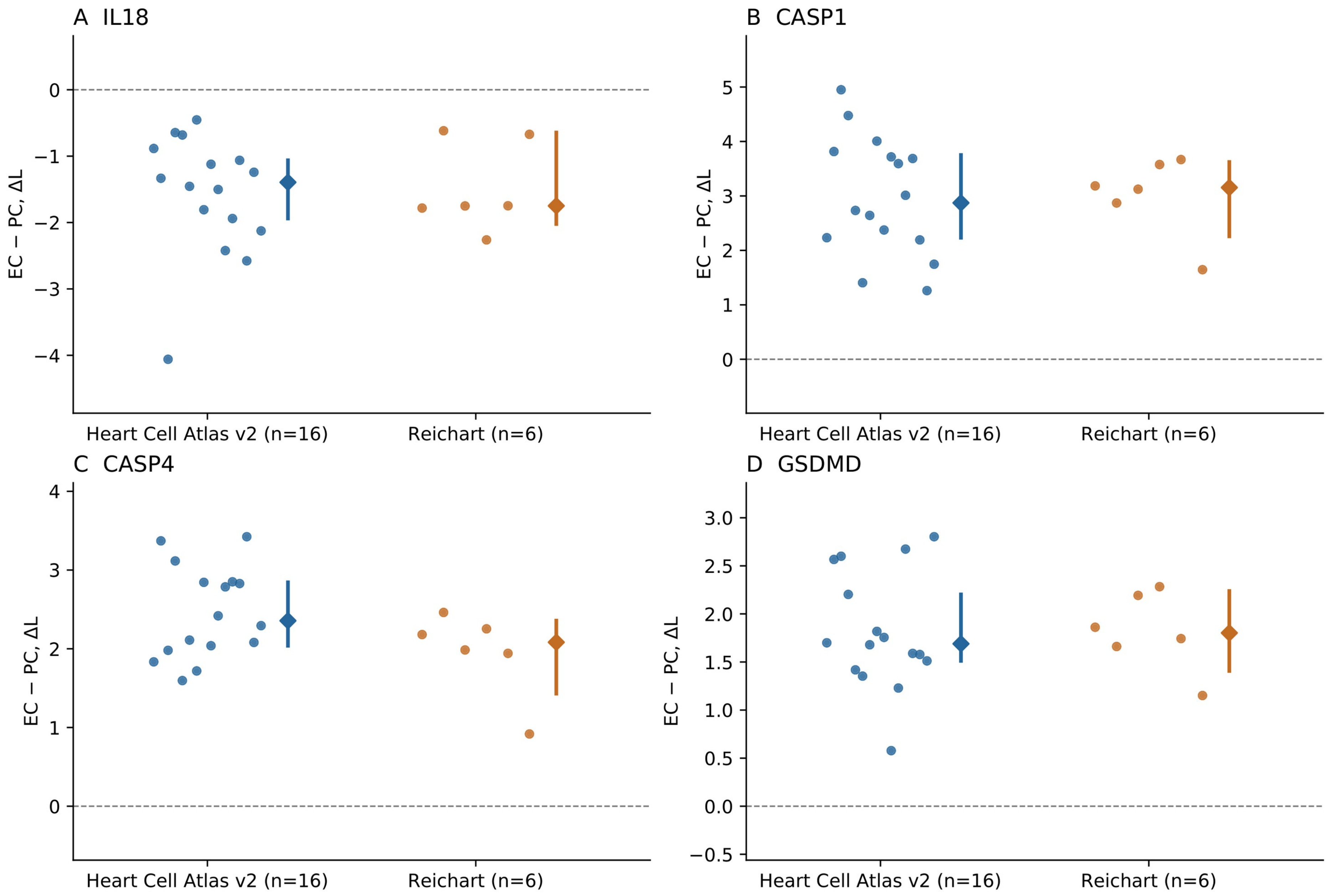

**Supplementary Figure S2**

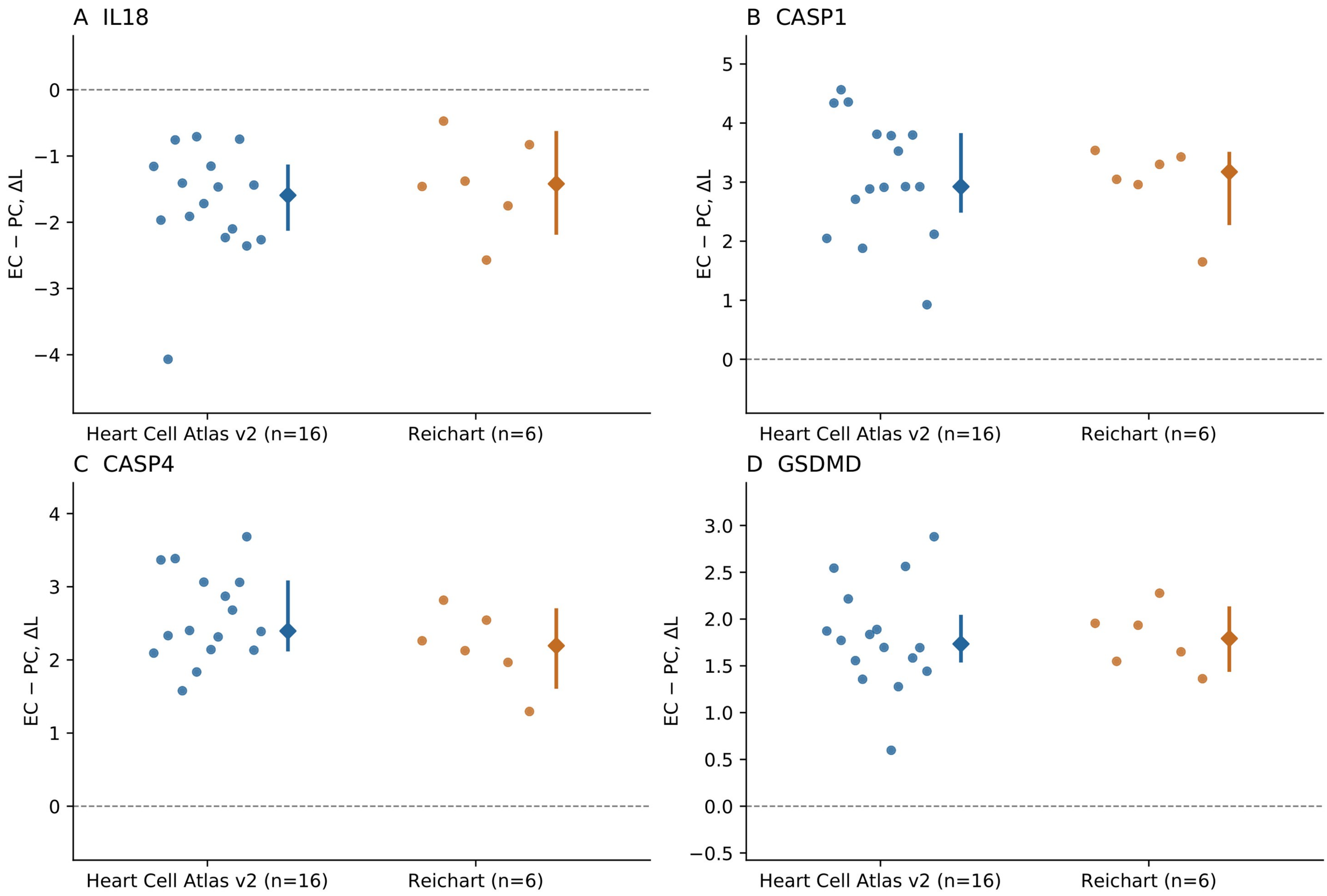

**Supplementary Figure S3**

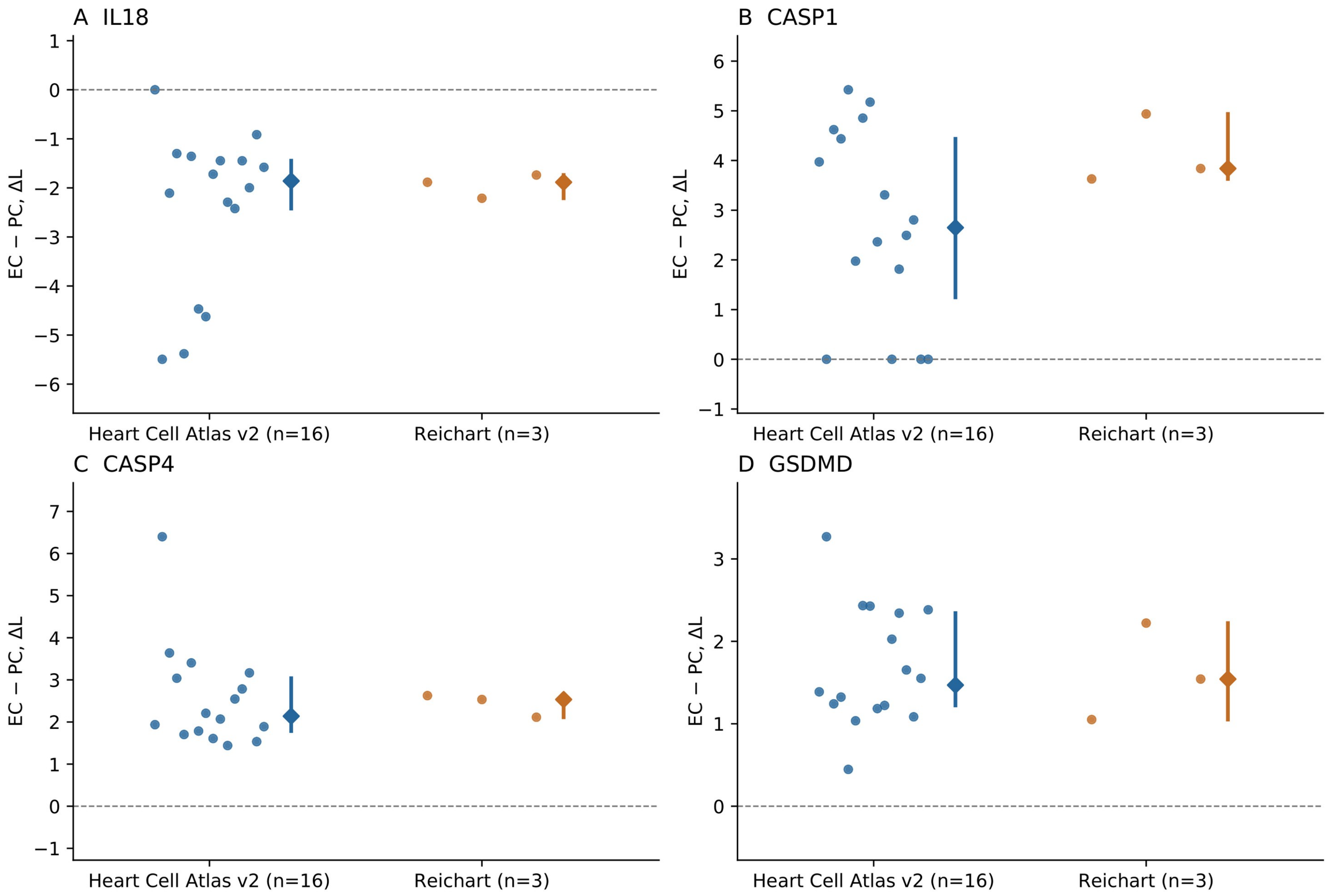

**Supplementary Figure S4**

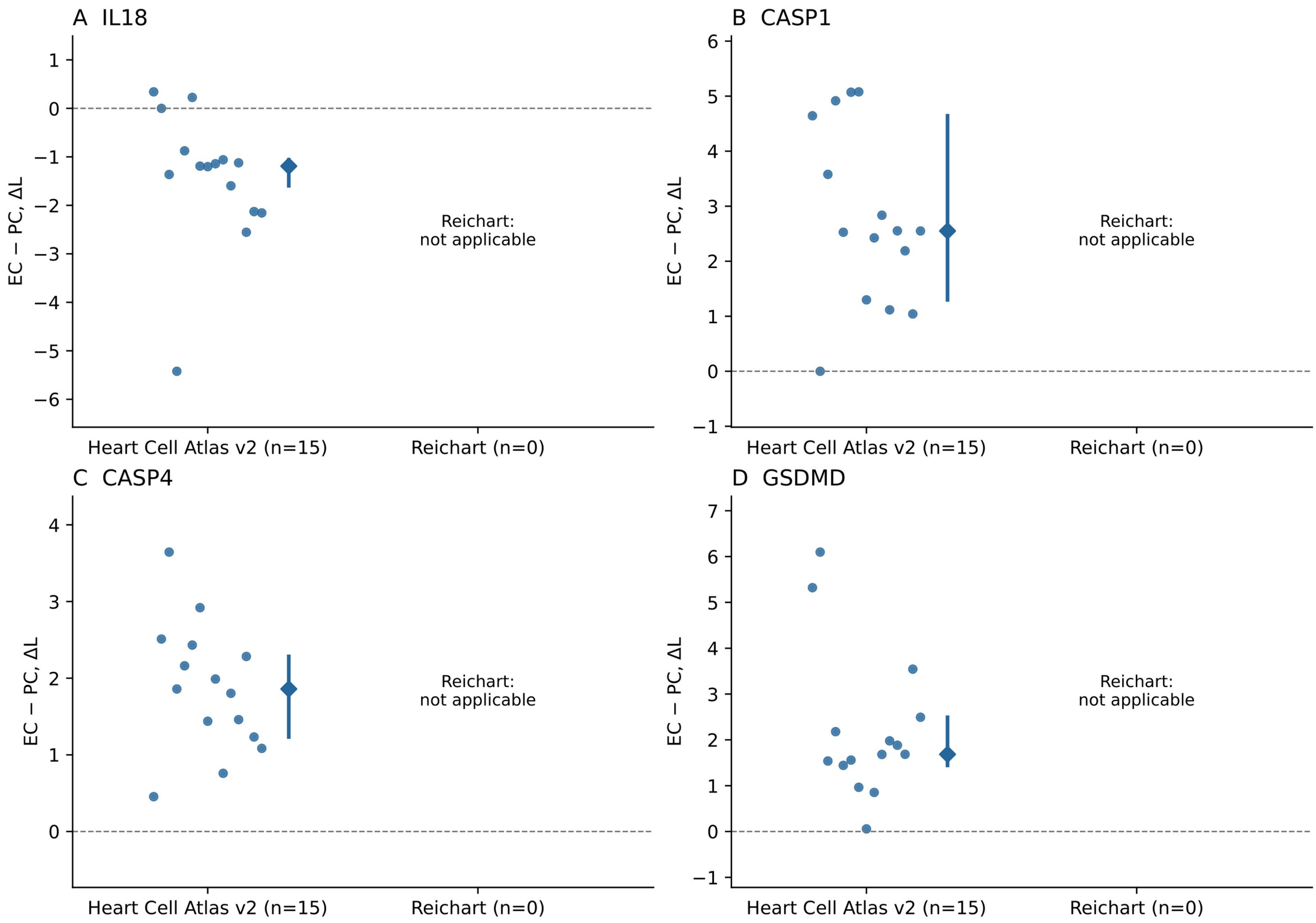

**Supplementary Figure S5**

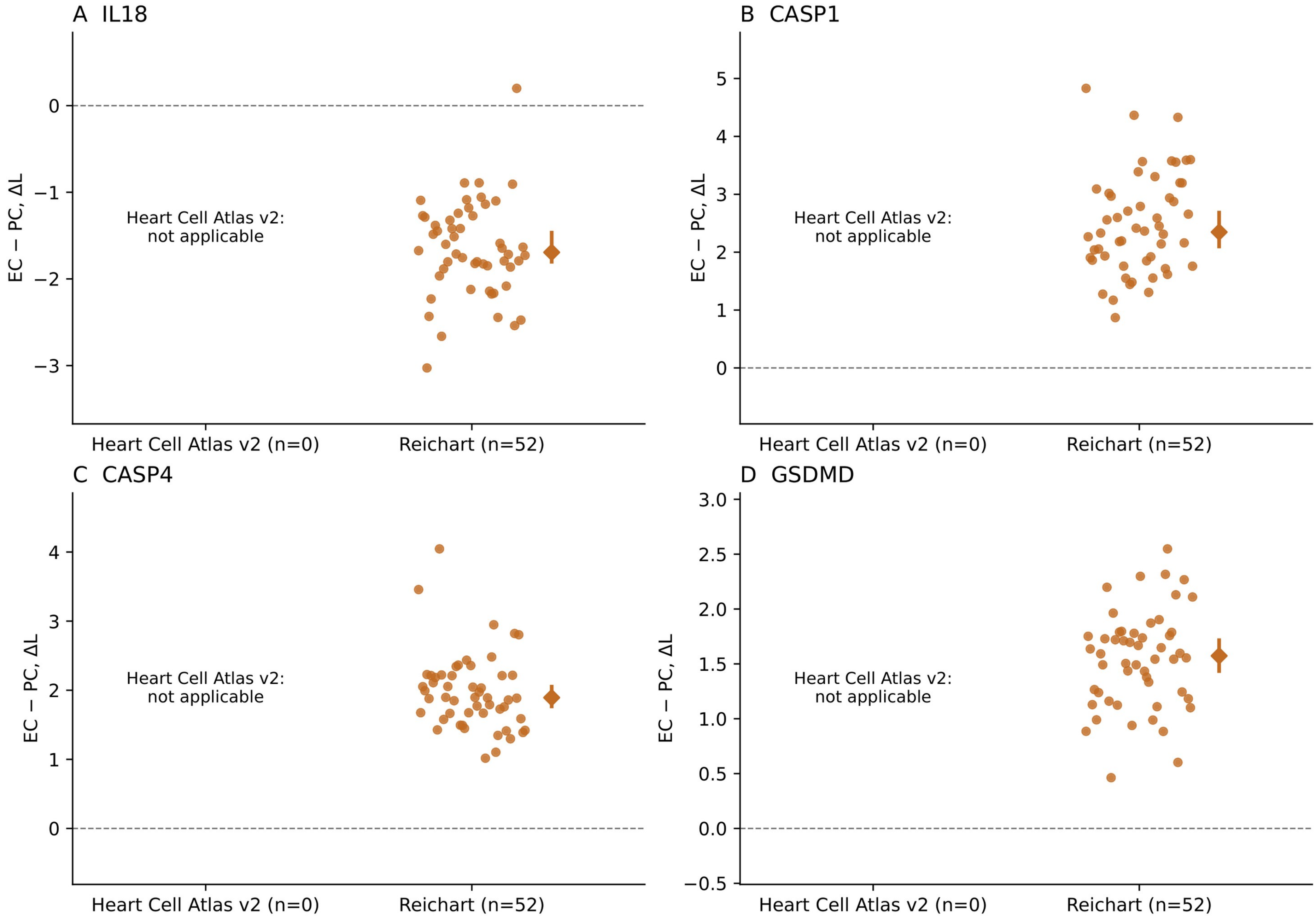

#### Supplementary Figure S6

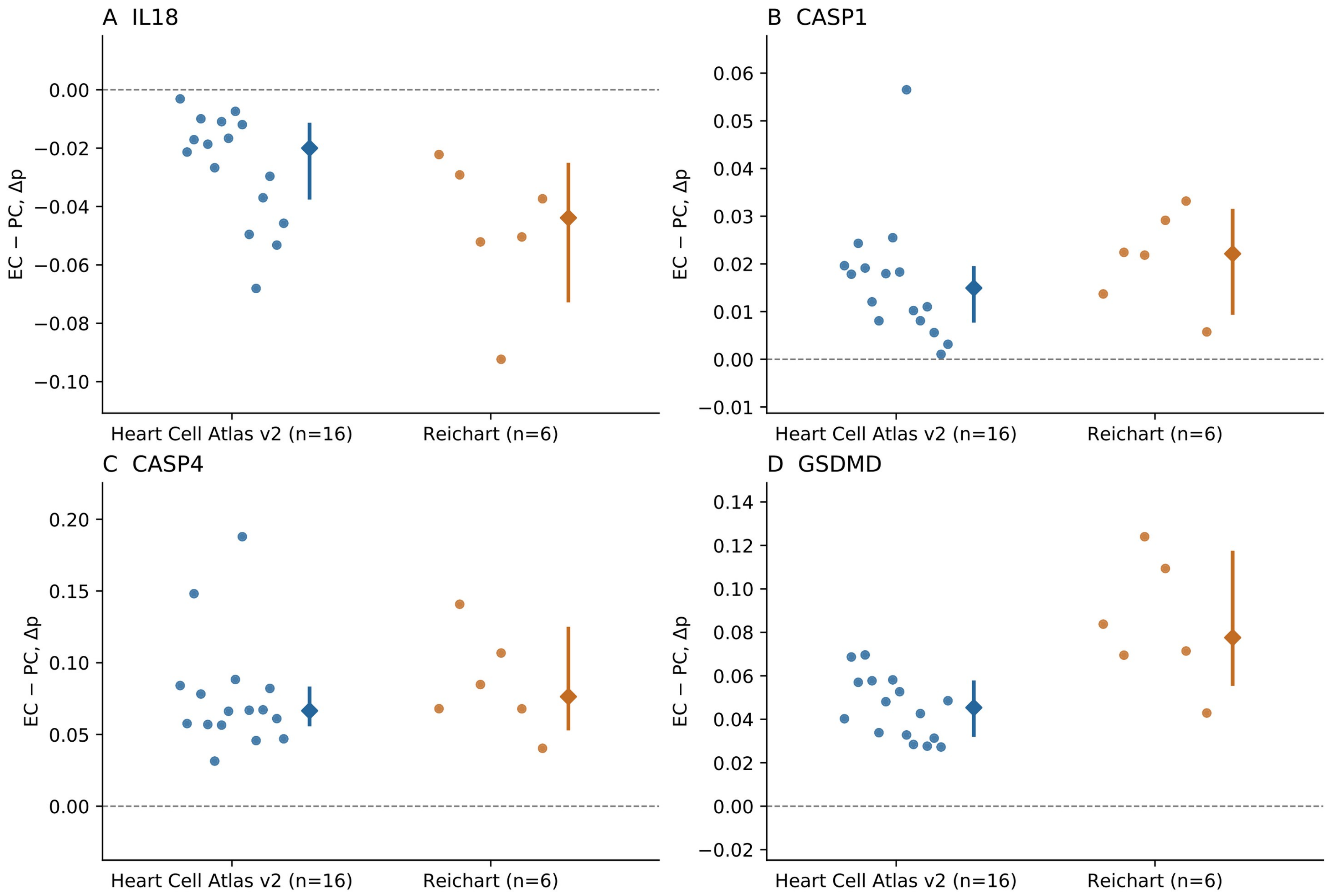

**Supplementary Figure S7**

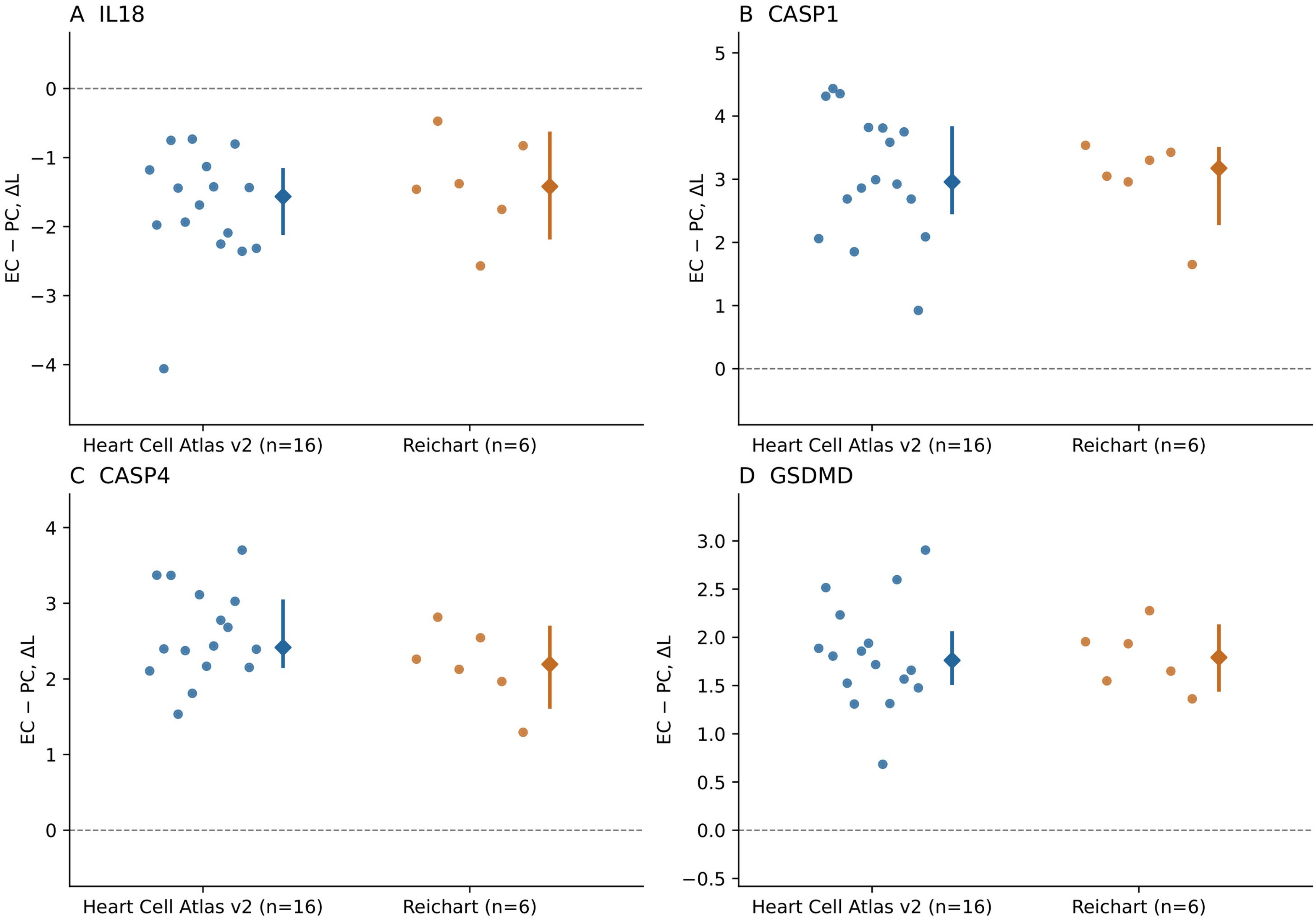

#### Supplementary Figure S8

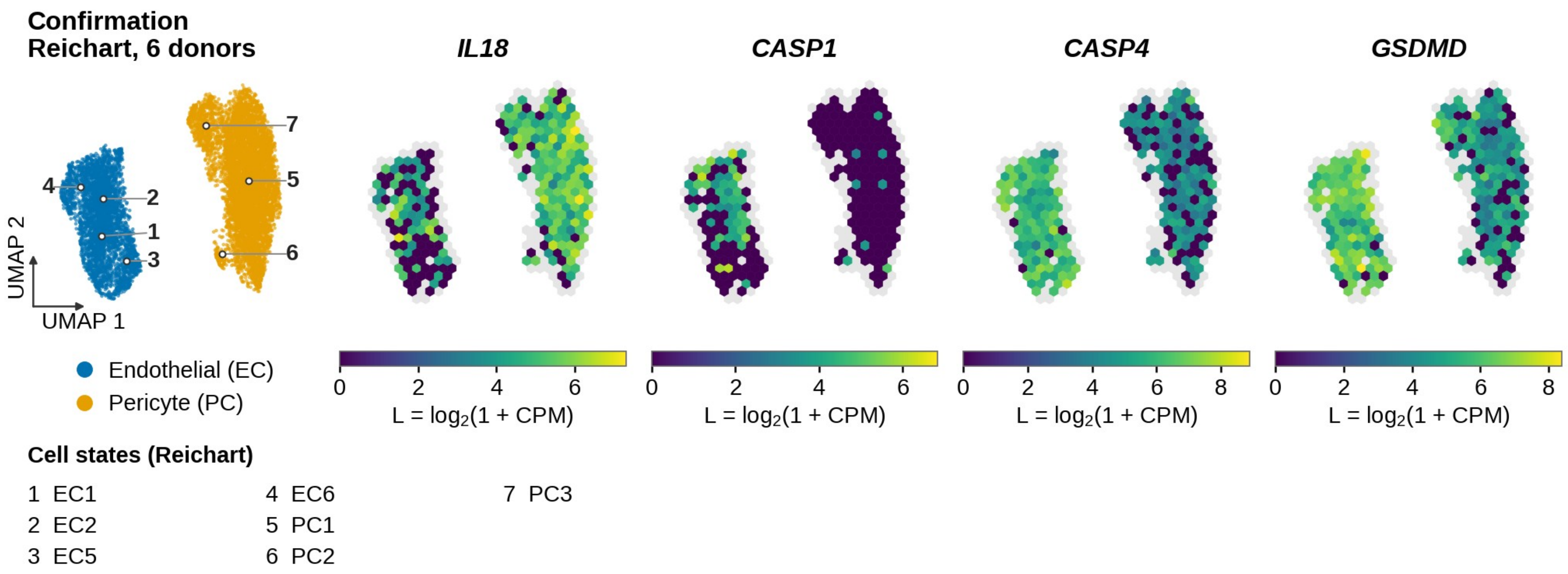

### Supplementary Figure S9

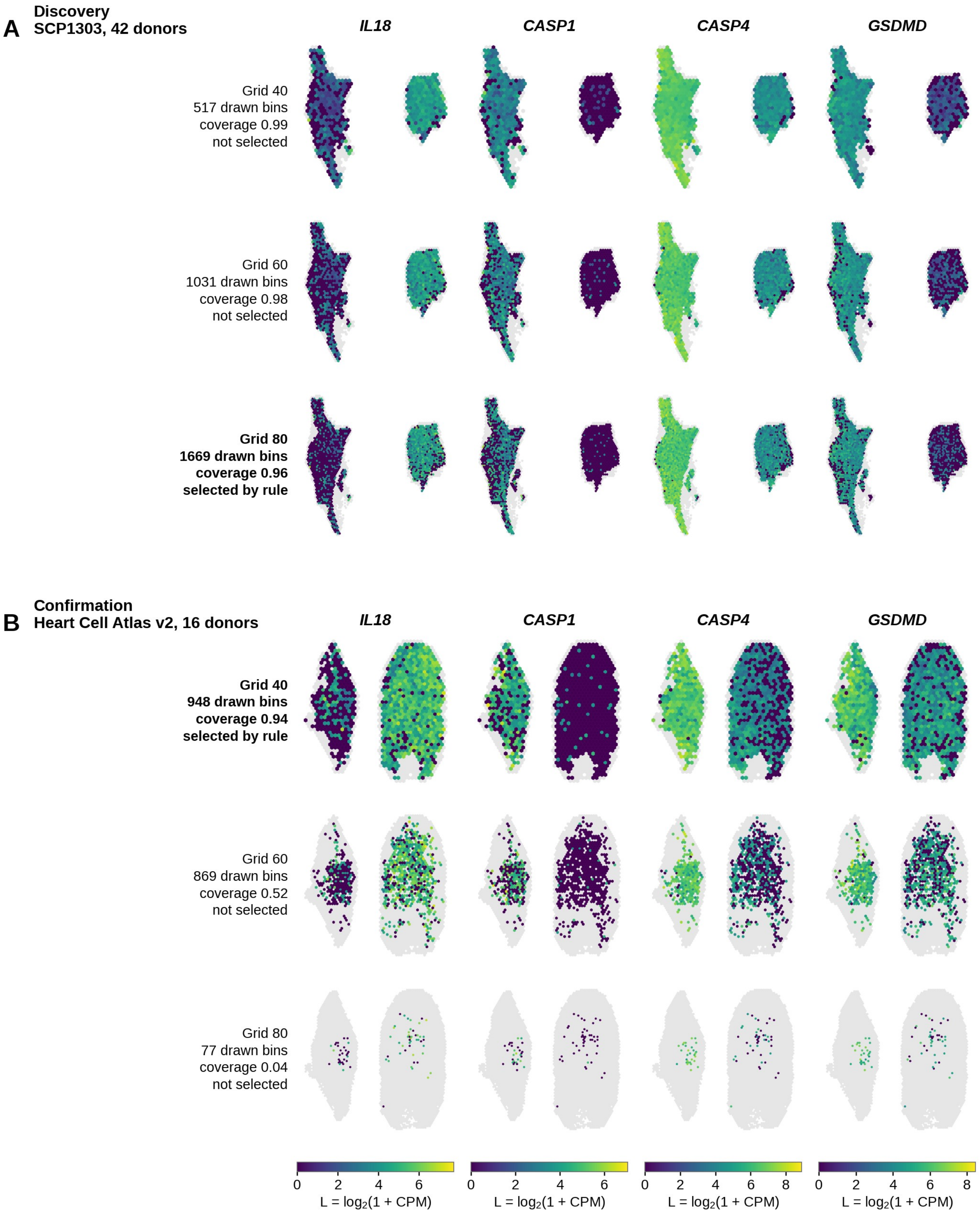

**Supplementary Figure S10**

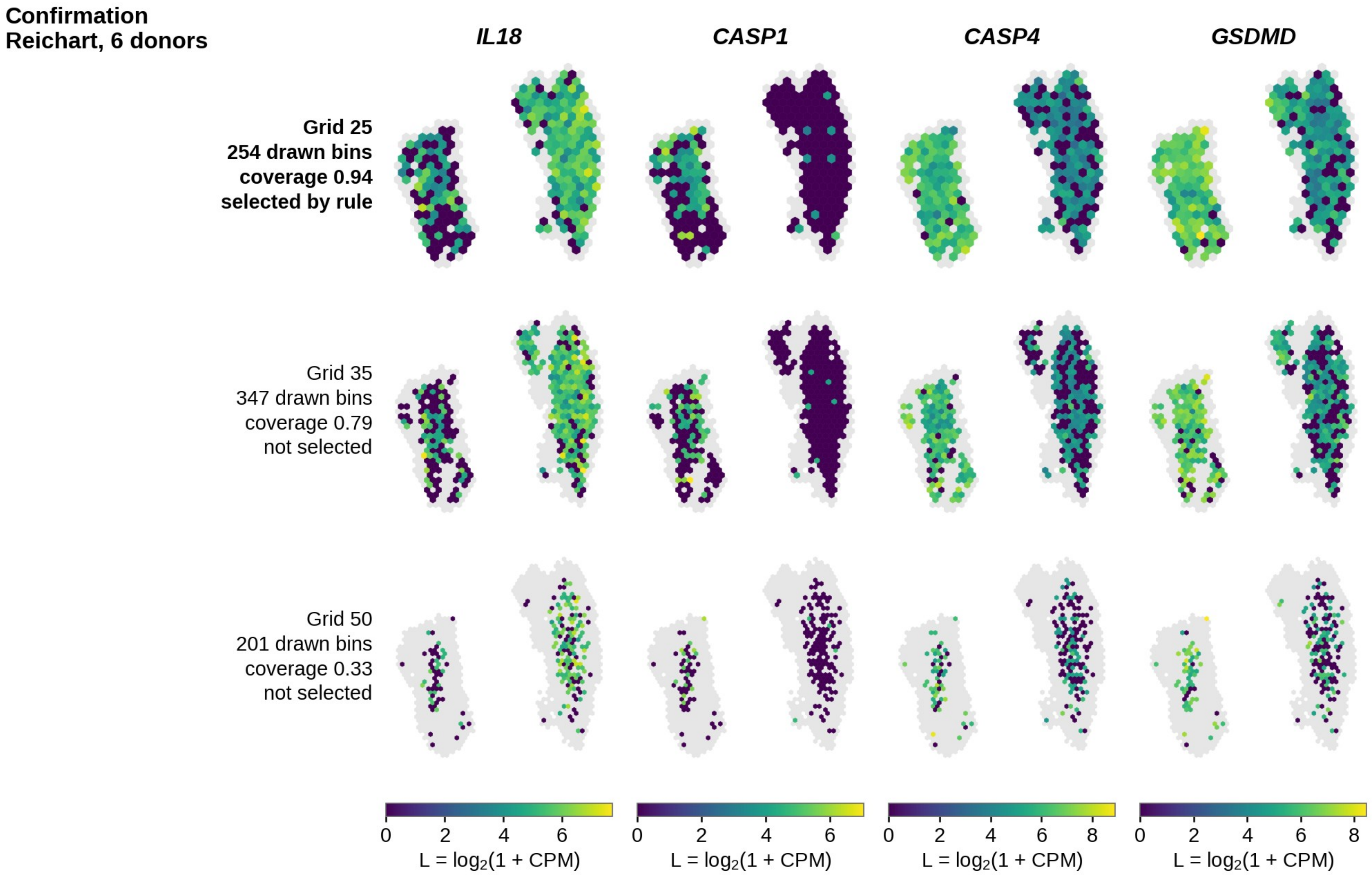

#### Supplementary Figure S11

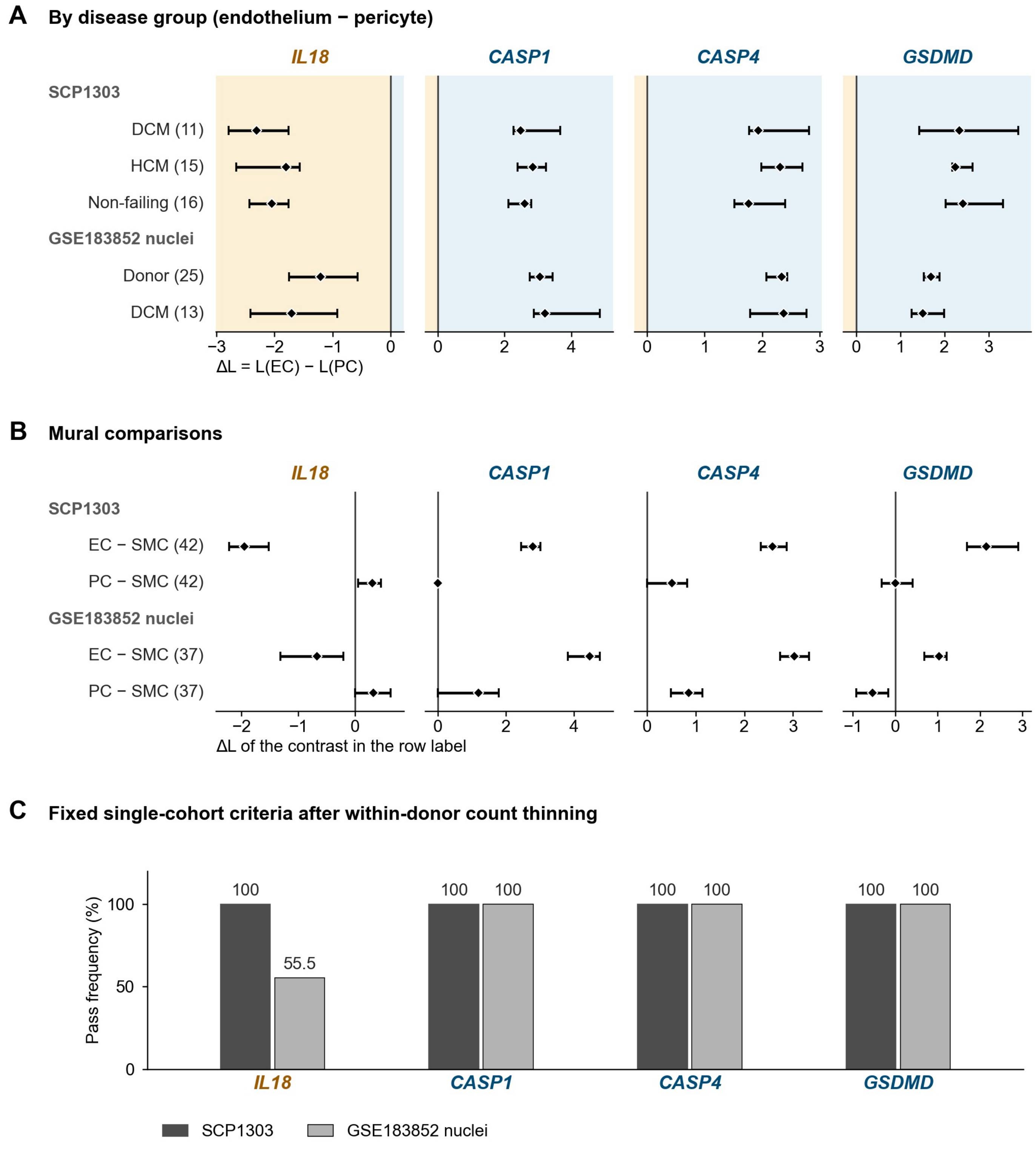

#### Supplementary Figure S12

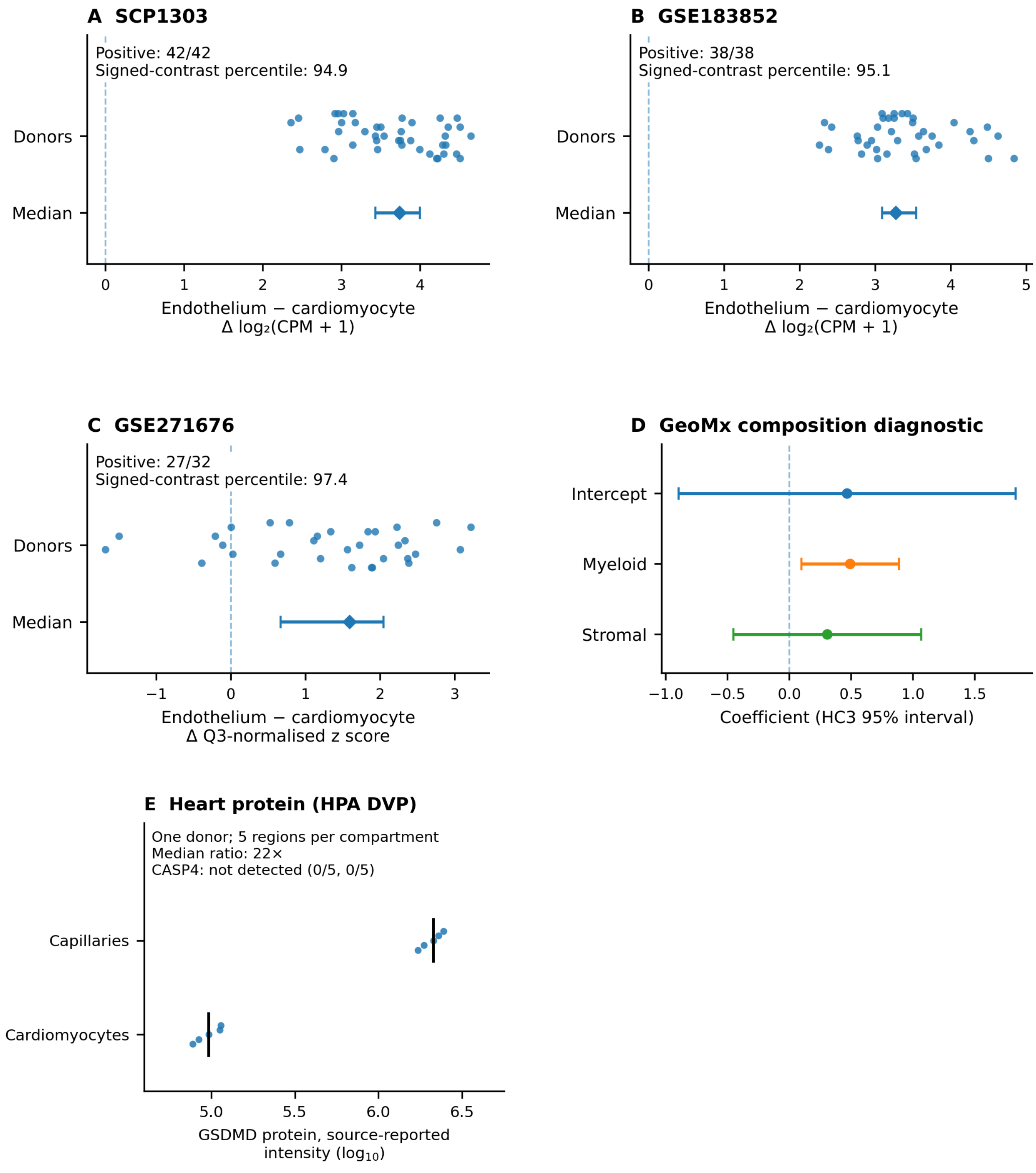

### Supplementary Figure S13

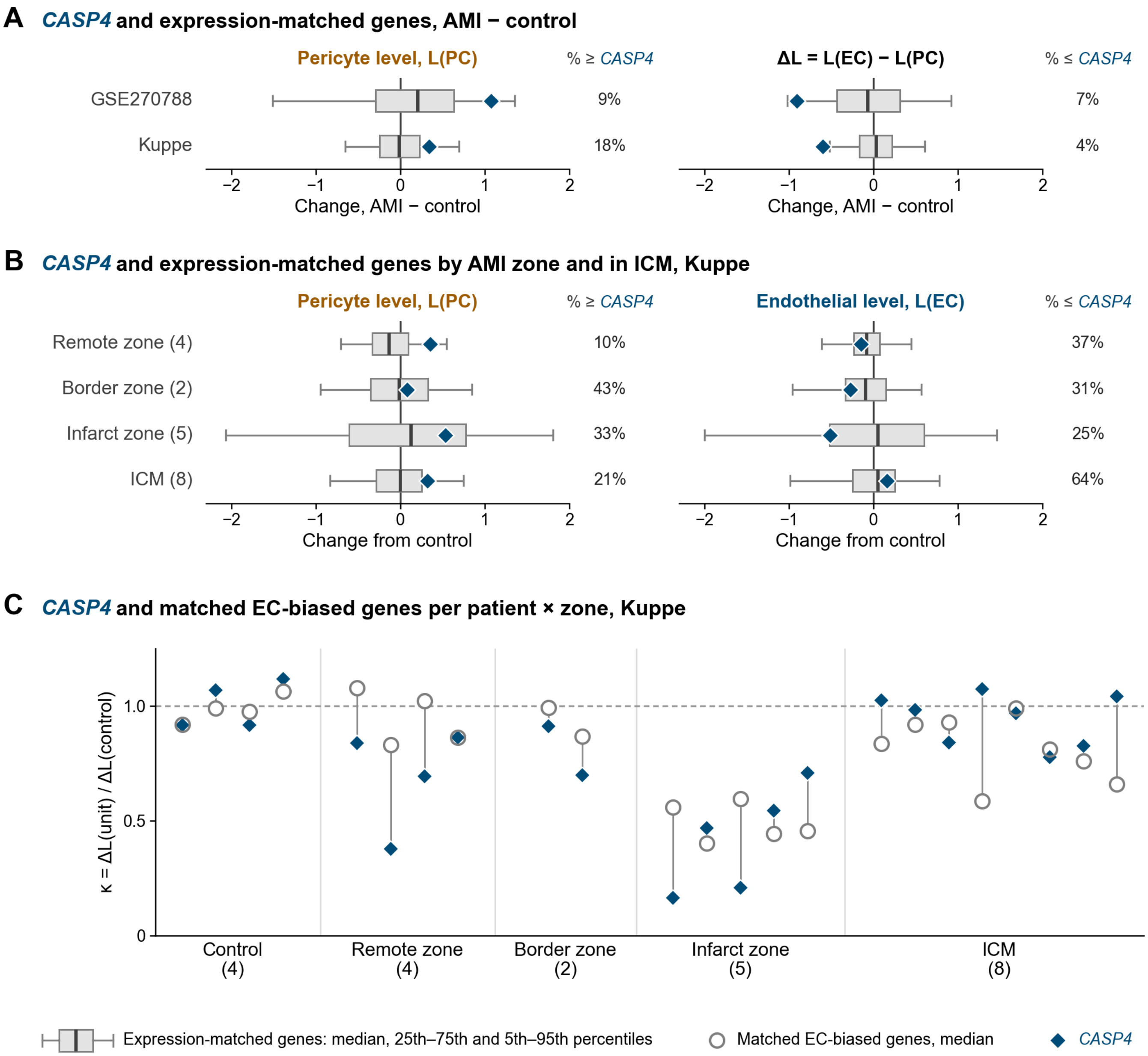

#### Supplementary Figure S14

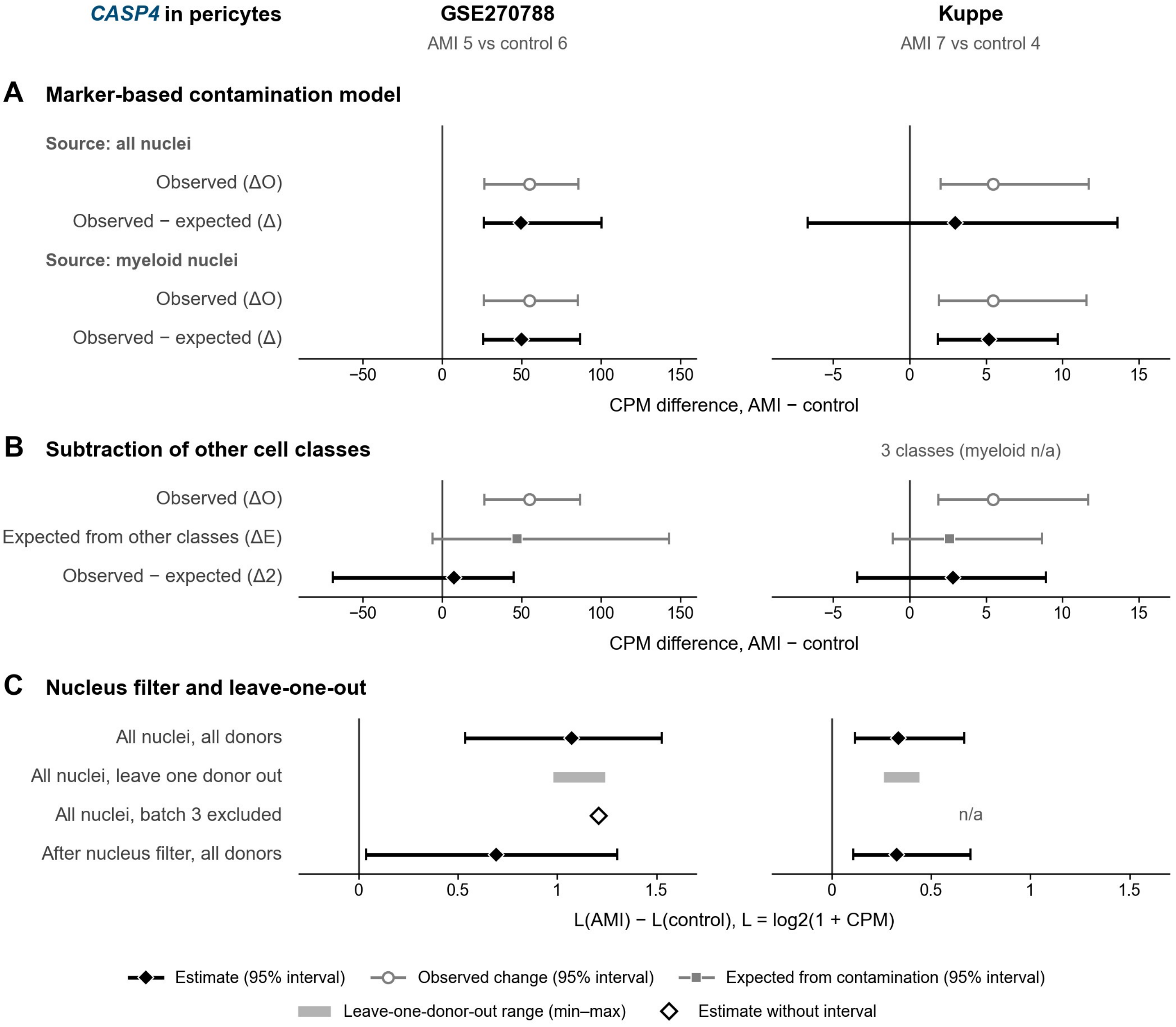

#### Supplementary Figure S15

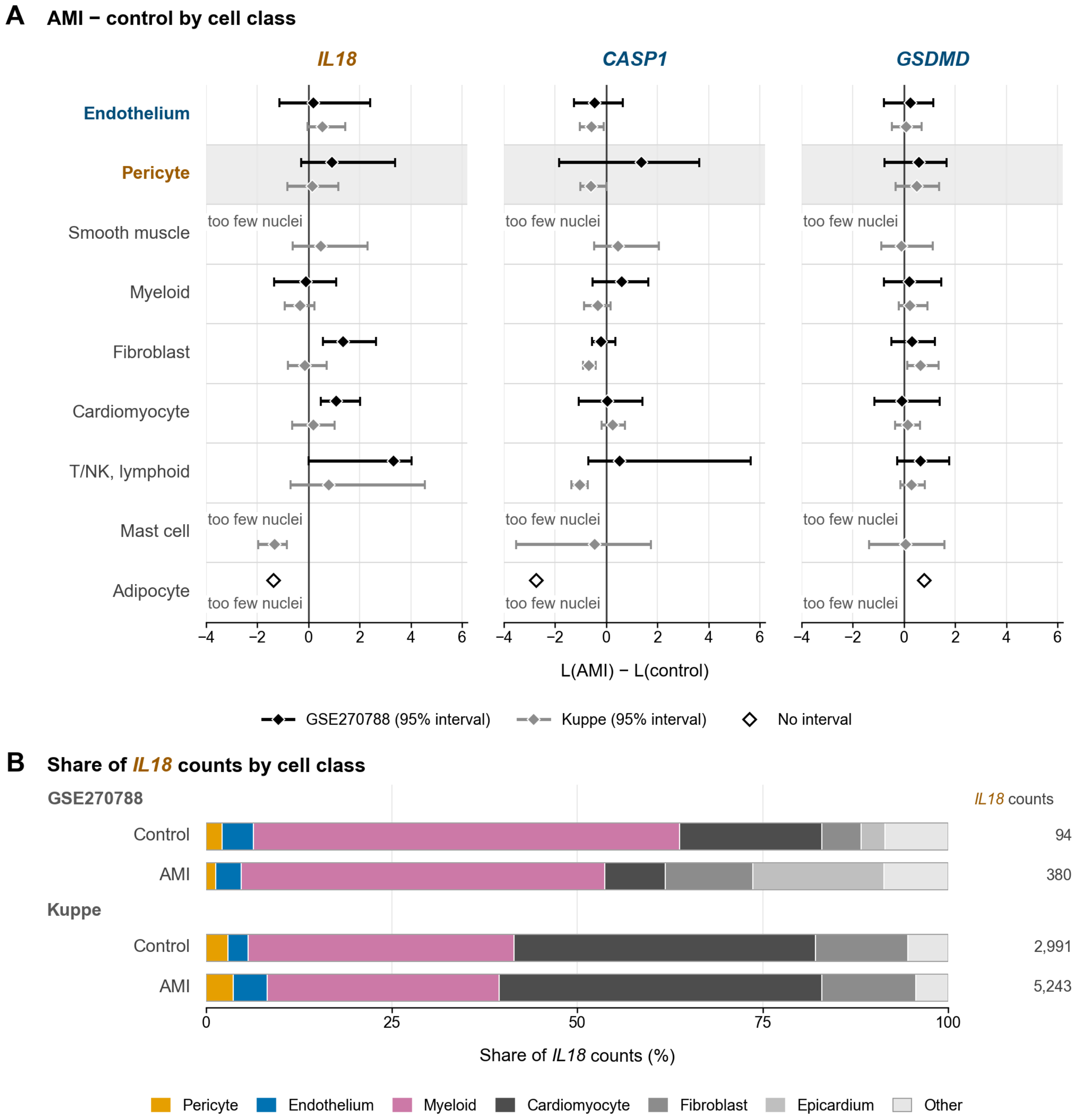

#### Supplementary Figure S16

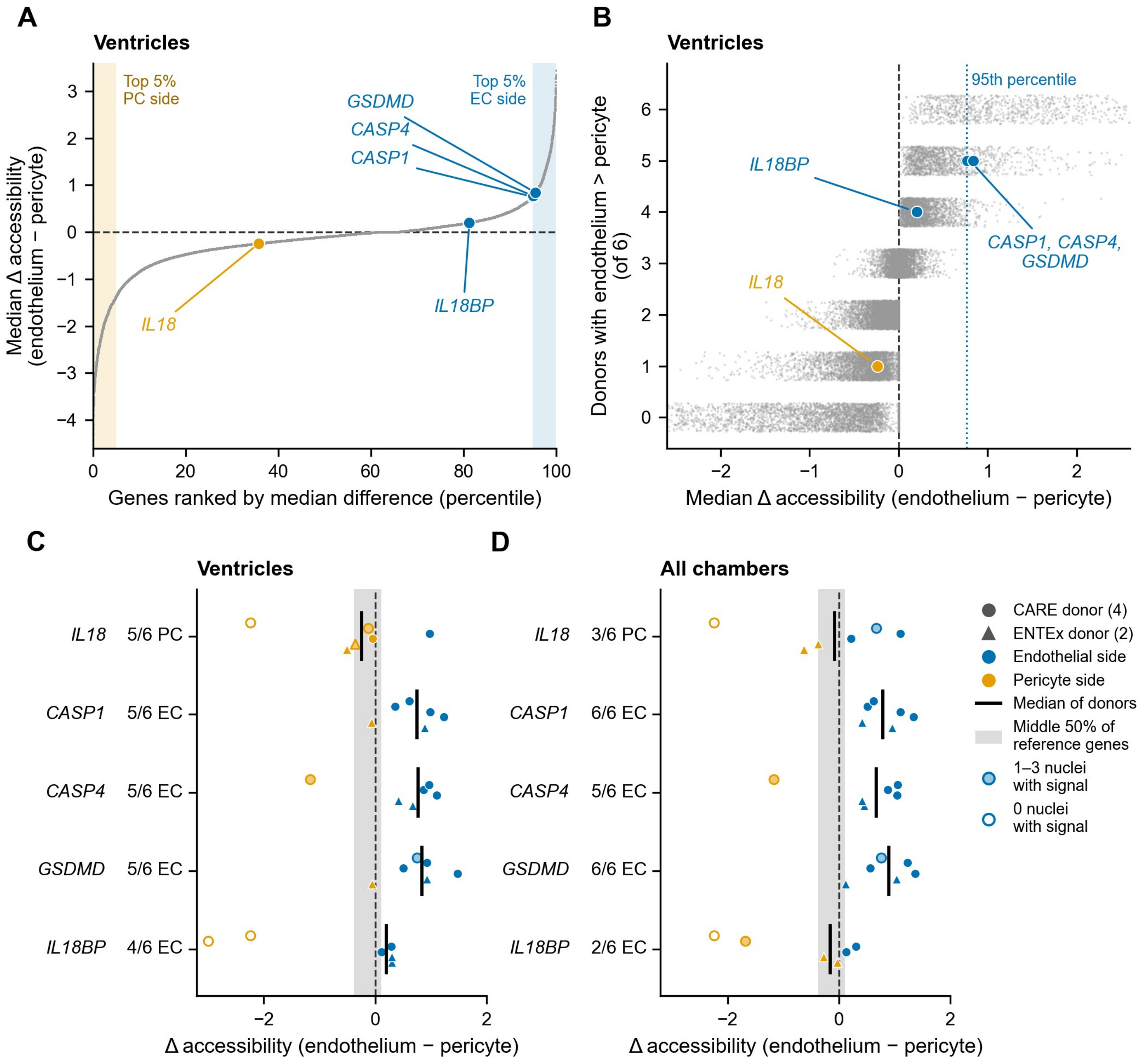
